# Compositional Inductive Biases in Function Learning

**DOI:** 10.1101/091298

**Authors:** Eric Schulz, Joshua B. Tenenbaum, David Duvenaud, Maarten Speekenbrink, Samuel J. Gershman

## Abstract

How do people recognize and learn about complex functional structure? Taking inspiration from other areas of cognitive science, we propose that this is achieved by harnessing compositionality: complex structure is decomposed into simpler building blocks. We formalize this idea within the framework of Bayesian regression using a grammar over Gaussian process kernels, and compare this approach with other structure learning approaches. Participants consistently chose compositional (over non-compositional) extrapolations and interpolations of functions. Experiments designed to elicit priors over functional patterns revealed an inductive bias for compositional structure. Compositional functions were perceived as subjectively more predictable than non-compositional functions, and exhibited other signatures of predictability, such as enhanced memorability and reduced numerosity. Taken together, these results support the view that the human intuitive theory of functions is inherently compositional.

## Introduction

Recognizing functional patterns is a ubiquitous problem in everyday cognition, underlying the perception of time, space and number. How much food should you cook to satisfy every guest at a party? How far do you have to turn the faucet handle to get the right temperature? Should you invest in a particular stock that seems to be going up? Since the space of such mappings is theoretically infinite, inductive biases are necessary to constrain the plausible inferences (Griffiths, Chater, Kemp, Perfors, & Tenenbaum, 2010; Mitchell, 1980). But what is the nature of these inductive biases over functions?

The major theoretical accounts of function learning, which we review below, offer two different answers to this question. Rule-based accounts posit a fixed set of parametric forms (or rules) that serve as a “vocabulary” for functions; these accounts imply strong inductive biases for the rules in the functional vocabulary. By contrast, similarity-based accounts posit a nonparametric representation of functions, implying relatively weak inductive biases.

A major challenge for humans is how to accommodate the virtually infinite diversity of functions in the real world. Rule-based models can only represent linear combinations of a fixed set of parametric functions. Similarity-based models can in principle represent an infinite variety of functions, but their typically weak inductive biases do not support strong inferences from small amounts of data—an important characteristic of human learning (Lake, Ullman, Tenenbaum, & Gershman, 2016).

A ubiquitous strategy in many areas of cognition, from language (Chomsky, 1965) to concept learning (Goodman, Tenenbaum, Feldman, & Griffiths, 2008; Kemp, 2012; Piantadosi, Tenenbaum, & Goodman, 2016) and visual perception (Biederman, 1987; Lake, Salakhutdinov, & Tenenbaum, 2015), is to divide and conquer: construct complex representations out of simpler building blocks using a set of compositional rules. Compositional systems support strong inferences from small amounts of data by imposing structural constraints, without sacrificing the capacity for representing an infinite variety of forms. The primary claim in this paper is that human function learning is structurally constrained by compositional inductive biases.

To formalize this idea, we need a theoretical framework for function learning that can represent and reason about compositional function spaces. Lucas, Griffiths, Williams, and Kalish (2015) recently presented a normative theory of function learning using the formalism of Gaussian processes (GPs). As we will describe more formally, GPs are distributions over functions that can encode properties such as smoothness, linearity, periodicity, symmetry, and many other inductive biases found by past research on human function learning (Brehmer, 1974b; DeLosh, Busemeyer, & McDaniel, 1997). Lucas et al. (2015) showed how Bayesian inference with GP priors can be expressed in both parametric (rule-based) and nonparametric (similarity-based) forms. GPs can therefore serve as a computational-level theory of function learning that bridges different mechanistic implementations.

In this paper, we build on the GP formalism to study, both theoretically and experimentally, the compositional nature of inductive biases in human function learning. Our extensions of the GP formalism not only bridge the “rules” and “similarity” perspectives on learning, but can also explain how people are able to learn much more complex kinds of functional relationships that are not well described by either traditional notions of rules or traditional kinds of similarity metrics.

Our main theoretical contribution is to extend the GP approach to modeling human function learning with a prior that obeys compositionally structured constraints. We do this using a compositional grammar for intuitive functions introduced in the machine learning literature by Duvenaud, Lloyd, Grosse, Tenenbaum, and Ghahramani (2013). We then test the predictive and explanatory power of this compositional GP model in 10 function learning and reasoning experiments, comparing the compositional prior to a flexible non-compositional prior (the spectral mixture representation proposed by Wilson & Adams, 2013, which we will describe later). Both models use Bayesian inference to reason about functions, but differ in their inductive biases.

Our experiments begin by comparing these different models of human function learning on five functional pattern completion tasks, two of which ask participants to choose among different completions, two of which assess a restricted posterior distribution over compositional kernels, and one of which asks participants to manually complete sampled functions within a graphical user interface.

Throughout all of these experiments, we find that participants’ completions are better described by the compositional prior as compared to non-compositional alternatives. We then generate a set of 40 similar functions, 20 of which are compositional and 20 of which are non-compositional, and compare these functions by asking participants how predictable they are and letting them learn and predict these functions in two experiments using a trial-by-trial function learning paradigm. We find that participants not only perceive compositional functions as more predictable and learn them more easily, but also that a compositional model of both predictability and function learning provides a quantitatively accurate description of participants’ behavior. Finally, we investigate how compositional functions influence memory, change detection, and the perception of numerosity in three additional experiments. To understand these results computationally, we propose a compositional Bayesian model of pattern encoding, chunking and retrieval, and compare this model to other alternatives. We conclude by discussing the implications as well as possible limitations of our proposed model and spell out future directions of compositional function learning research.

## Prior research on human function learning

The general problem of inferring how one variable depends functionally on another is important for many aspects of cognition. Traditionally, it has been studied in paradigms assessing how people learn about input-output mappings or how they make sense of spatiotemporal patterns. Additionally, the way in which we learn about and recognize functional patterns is also crucial in the modern world as we look at data – either as scientists or non-scientist decision makers – in trying to understand what function the data reveal. We are interested in all of these aspects of function learning, but will focus first on the latter because it can potentially reveal how people recognize and perform inference about structure very quickly. However, we believe that a strength of our account of compositional inductive biases is that it can account for empirical effects across a diverse set of paradigms.

Donald Broadbent was among the first psychologists to investigate how people learn and control functions between inputs and outputs (Broadbent, 1958). In his experiments, participants controlled functions within an industrial setting called the “sugar factory,” in which they learned the relationship between work force and sugar production. Broadbent showed that participants had difficulty controlling some functions, such as exponential or power functions, but were good at learning others, for example linear functions (Berry & Broadbent, 1984).

Since Broadbent’s pioneering work, further studies have established several empirical regularities (see McDaniel & Busemeyer, 2005, for a review). For example, studies using interpolation judgments—predictions of function outputs for inputs inside the convex hull of training inputs—have found that linear, increasing functions are easier to learn than non-linear, non-monotonic or decreasing functions (Brehmer, 1974a; Brehmer, Alm, & Warg, 1985; Byun, 1995). Presentation order also matters: it is easier to learn functions if the input is ordered by increasing output (DeLosh et al., 1997).

Important constraints on theories of function learning have come from studies of extrapolation judgments—predictions of function outputs for inputs outside the convex hull of training inputs (DeLosh et al., 1997; McDaniel & Busemeyer, 2005). People tend to make linear extrapolations with a positive slope and an intercept of zero (Kalish, Lewandowsky, & Kruschke, 2004; Kwantes & Neal, 2006). This linearity bias holds true even when the underlying function is non-linear; for example, when trained on a quadratic function, average predictions fall between the true function and straight lines fitted to the closest training points (DeLosh et al., 1997).

When Little and Shiffrin (2009) asked participants to produce judgments about the best causal function underlying noisy data, they found that participants also exhibit a simplicity bias, i.e. that a probability distribution over polynomial degrees obtained over all participants tended to put most of its mass on smaller degrees.

Two families of theories have been developed to explain these and other regularities. Rule-based theories (Carroll, 1963; Koh & Meyer, 1991) propose that people learn explicit parametric functions, for example linear, polynomial or power-law functions, or some combination of simpler functions. While rule-based theories have had some success in predicting function learning performance, they have trouble accounting for the aforementioned “order-of-difficulty effects” in interpolation tasks (McDaniel & Busemeyer, 2005), fail to fully predict extrapolation performance (DeLosh et al., 1997), especially for non-linear functions, and are unable to learn a partitioning of the input space (the knowledge partitioning effect; Kalish et al., 2004).

Similarity-based theories (e.g., Busemeyer, Byun, Delosh, & McDaniel, 1997; DeLosh et al., 1997) propose that people learn functions by associating inputs and outputs: if input x is paired with output *y*, then inputs similar to **x** should produce outputs that are similar to *y*. Busemeyer et al. (1997) formalized this intuition using a connectionist model (Associative-Learning Model; ALM) in which inputs activate an array of hidden units representing a range of possible input values; each hidden unit is activated in proportion to its similarity to the current input. Learned associations between the hidden units and the response units map the similarity-based activation pattern to output predictions. ALM successfully captures aspects of interpolation performance, but fails to explain extrapolation and knowledge partitioning phenomena.

In order to overcome some of these problems, hybrid versions of the two approaches have been put forward (McDaniel & Busemeyer, 2005). Hybrid models normally contain an associative learning process that acts on explicitly-represented rules. One such hybrid is EXAM (Extrapolation-Association Model; Busemeyer et al., 1997). EXAM assumes similarity-based interpolation, but extrapolates using a simple linear rule. The model effectively captures the human bias towards linearity, and predicts human extrapolations for a variety of functions, but without accounting for non-linear extrapolation (Bott & Heit, 2004). POLE (population of Linear Experts; Kalish et al., 2004), which approximates functions using piece-wise linear representations, can capture knowledge partitioning and order-of-difficulty effects (McDaniel, Dimperio, Griego, & Busemeyer, 2009). However, it is unclear if a combination of linear functions is able to explain human function learning in more complex, naturalistic contexts, especially given that humans are capable of learning non-linear functions as well (see Bott & Heit, 2004)^1^.

Another area of explaining how people make sense of functional structure could be called “intuitive science” or forecasting. Here, participants are normally confronted with multiple data points at once and then have to predict future or left out points.

Various reviews have established factors that influence participants predictions of visually presented data (see Bolger & Harvey, 1998; Goodwin & Wright, 1993, for example). Broadly speaking, these factors fall within 4 categories. First, people seem to damp trends when they make forecasts from noisy data. This means that their forecasts lie below upward trended lines but above downward ones. Therefore, it appears that forecasters tend to underestimate the steepness of functions (Andreassen & Kraus, 1990; Keren, 1983). Trend damping is greater for downward than for upward trended data, especially when the data representation format is visual rather than in a table (Harvey & Bolger, 1996). Second, forecasts tend to overestimate functions lacking a trend (Eggleton, 1982). Third, people seem to deliberately attach random noise to their forecasts, and add more noise to forecasts from noisier data series. This means that their forecasts appear to represent the way the series will appear once the outcome has occurred (Harvey, Ewart, & West, 1997). Fourth, forecasts for independent series should lie on the series mean, but instead they have been found to lie between the mean and the last revealed data point (Eggleton, 1982), similar to findings in more traditional function learning paradigms.

## Gaussian process regression as a normative theory of function learning

The theories of function learning summarized in the previous section are process models, specifying mechanistic hypotheses about representations and learning rules. However, mechanistic hypotheses do not directly give insight into inductive biases, since different mechanisms may or may not produce the same bias. Thus, if our goal is to understand human inductive biases, we require a computational-level analysis of function learning that is agnostic to mechanism. The GP theory of function learning developed by Lucas et al. (2015) fills this gap.

Formally, a GP is a collection of random variables, any finite subset of which are jointly Gaussian-distributed (see Rasmussen & Williams, 2006; Schulz, Speekenbrink, & Krause, 2017). A GP can be expressed as a distribution over functions. Let 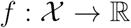 denote a function over input space 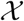 that maps to real-valued scalar outputs.^2^ The function can be modeled as a random draw from a GP:

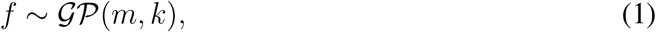

where *m* is a mean function specifying the expected output of the function given input **x**, and *k* is a kernel (or covariance) function specifying the covariance between outputs.

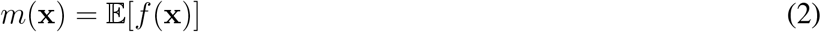

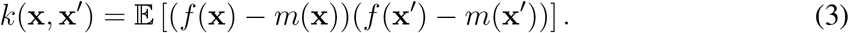

Intuitively, the mean function encodes an inductive bias about the expected shape of the function, and the kernel encodes an inductive bias about the expected smoothness. This does not necessarily imply that distributions of outputs over different input points have to be Gaussian as this would also depend on an added noise term which does not have to come from a Gaussian distribution. To simplify exposition, we follow standard convention in assuming a prior mean of **0**.

Conditional on observed data 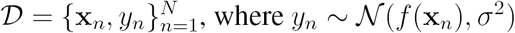 is a noise-corrupted draw from the latent function, the posterior predictive distribution for a new input x_*_ is Gaussian with mean and variance given by:

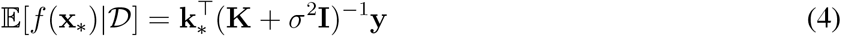

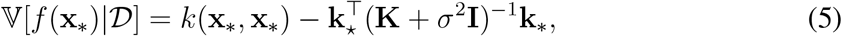

where y = [*y*_1_,…, *y*_*N*_]^T^, K is the *N* × *N* matrix of covariances evaluated at each pair of observed inputs, and k_*_ = [*k*(x_1_, x_*_),…, *k*(x_*N*_, x_*_)] is the covariance between each observed input and the new input x_*_. See Appendix A for further technical details.

As pointed out by Griffiths, Lucas, Williams, and Kalish (2009) (see also Lucas et al. 2015), the predictive distribution can be viewed as an exemplar (similarity-based) model of function learning (DeLosh et al., 1997; McDaniel & Busemeyer, 2005), since it can be written as a linear combination of the covariance between past and current inputs:

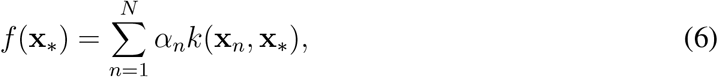

where *α* = (K + *σ*^2^I)^−1^y. Equivalently, by Mercer’s theorem any positive definite kernel can be expressed as an outer product of feature vectors:

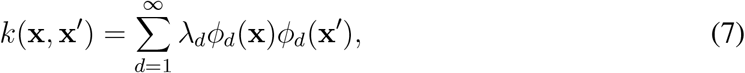

where {*ϕ*_*d*_(x)} are the eigenfunctions of the kernel and {*λ*_*d*_} are the eigenvalues. The posterior predictive mean is a linear combination of the features, which from a psychological perspective can be thought of as encoding “rules” mapping inputs to outputs (Carroll, 1963; Koh & Meyer, 1991). Thus, a GP can be expressed as both a similarity-based model and as a rule-based model, thereby unifying the two dominant families of function learning theories in cognitive science (described in detail by Lucas et al., 2015).

## Understanding human inductive biases with Gaussian processes

Gaussian processes provide a concrete language for constructing and comparing priors over functions, which in turn encode different inductive biases. Priors over functions can be explicitly encoded by the kernel, which means—as we will describe in more detail below—that the kernel can be used to describe expectations about functional regularities (e.g., linear trends, periodic patterns, etc.) directly into the prior, thereby creating inductive biases that make learning these regularities easier.

Intuitively, a GP favors functions that are “smooth” as encoded by the kernel function.^3^ The critical question then is: what kind of smoothness do humans prefer?

Experiments have begun to study this question. One experimental technique, known as *iterated learning,* estimates priors by simulating a Markov chain across multiple people—essentially a “game of telephone”. Under certain assumptions, the Markov chain is guaranteed to converge to a stationary distribution, such that asymptotically it will generate samples from the prior (Griffiths & Kalish, 2007). In a function learning version of this task, participants see a set of points, which they then have to remember and re-draw. The resulting points are presented to another participant, who is asked to repeat the same procedure. Iterating this procedure multiple times will reveal participants’ priors over functions. Kalish, Griffiths, and Lewandowsky (2007) showed that participants consistently converged to linear functions with a positive slope, even when the starting points came from a linear function with a negative slope, quadratic functions, or when they were generated at random.

Lucas et al. (2015) explained the iterated learning results by postulating a GP with a mixture of experts kernel that is mostly dominated by a positively linear kernel ^4^ but can also generate smooth non-linear extrapolations by utilizing a non-linear radial basis function kernel. Additionally, their proposed kernel contained negatively linear and quadratic components. Samples from the GP parameterized in this way tend to be positively linear lines. This parametrization was able to explain a number of findings from the human function learning literature, such as the partitioning of the learning space, the linear-polynomial mixture-like patterns of extrapolations, and the difficulty of learning some functions (e.g., exponential patterns) relative to others (e.g., linear patterns).

Going further, Wilson, Dann, Lucas, and Xing (2015) attempted to infer the “human kernel” by having participants generate extrapolations for different functions sampled from a radial basis kernel and fitting a non-parametric kernel (the spectral mixture defined by Wilson & Adams, 2013, which we will describe in detail below) to their extrapolations. They found that participants expected long-distance correlations between points. These correlations could be viewed as arising from a mixture between linear and radial basis components^5^. In a second experiment, they showed that participants could effectively learn functions sampled from a mixture of a product of linear and spectral kernels.

These studies suggest that participants’ represent complex functions as compositions of simpler ones. In the next section, we will motivate compositionality as a core principle of cognition, and then proceed to formalizing a fully compositional theory of function learning.

## Beyond smoothness: compositionality as a core principle of function learning

Humans can adapt to a wide variety of structural forms (Gershman & Niv, 2010; Kemp & Tenenbaum, 2009); the space of such forms is essentially unbounded, raising the question of how an infinite variety of forms can be represented. One approach (arguably the only tractable approach) is to define a compositional system that can build complex structures through the composition of simpler elements. For example, in functional programming languages, functional primitives can be combined to create more complex functions which can then be re-combined to create even more complex functions (Peyton Jones, 1987). Via re-combinations, compositionality leads to a large increase of productivity—an infinite number of representations can be constructed from a finite set of primitives (Fodor, 1975; Fodor & Pylyshyn, 1988).

One source of evidence for compositionality in cognition comes from studies of rule-based concept learning. In these studies, the rules can be expressed as functions of logical primitives (e.g. Bruner, Goodnow, & George, 1956; Shepard, Hovland, & Jenkins, 1961). By studying participants’ mistakes and the relative learning difficulty of different concepts, researchers tried to unravel the primitives of symbolic thought and how these primitives are combined. This work has led to a rich set of theoretical ideas and empirical constraints on compositionality in concept learning (Goodman et al., 2008; Lake et al., 2015; Nosofsky, Palmeri, & McKinley, 1994; Piantadosi et al., 2016).

Kemp (2012) provided an exhaustive characterization of compositionality in logical domains, showing how a “conceptual universe” can be formed by a rule inference scheme based on minimal description length that explains logical reasoning data across a wide set of domains. Recently, Piantadosi et al. (2016) showed how different sets of structural primitives, embedded in an approximate Bayesian inference framework, can predict distinct learning curves in rule-based concept learning experiments. Based on these and other studies, Lake et al. (2016) argued that compositionality is a necessary requirement for designing algorithms “that learn and think like people.”

Compositionality has also been proposed as an important principle of motor learning, in particular in research on modular decomposition (Wolpert & Ghahramani, 2000). For example, Flanagan et al. (1999) showed that if subjects had to learn a combined transformation of kinematic and dynamic transformations, their reaching errors were smaller if they first learned each transformation separately, suggesting that people have the ability to combine and decompose different internal models of control. Ghahramani and Wolpert (1997) exposed participants to discrepancies between actual and visually perceived hand movements from two different starting positions and found that their movements for intermediate starting positions indicated an adaptive mixture of both individual strategies.

Given the widespread theoretical and empirical support for compositionality in cognitive science, it is natural to ask whether humans make use of compositionality in representing and learning about functions. With a few exceptions (Gershman, Malmaud, & Tenenbaum, 2016; Gershman, Tenenbaum, & Jäkel, 2016), prior work on function learning with GPs has assumed a fixed, non-compositional kernel^6^. Thus, our goal is to formalize a compositional approach to function learning and compare it to alternative non-compositional approaches. We accomplish this by positing two candidate kernel parametrizations that express conceptually different (compositional vs. non-compositional) inductive biases. We will then present a series of experimental tests that pit these kernels against each other.

## Structure learning with Gaussian processes

Broadly speaking, there are two approaches to parametrizing the kernel space: a fixed functional form with continuous parameters, or a combinatorial space of functional forms. These approaches are not mutually exclusive; indeed, the success of the combinatorial approach depends on optimizing the continuous parameters for each form. Nonetheless, this distinction is useful because it allows us to separate different forms of functional complexity. A function might have internal structure such that when this structure is revealed, the apparent functional complexity is significantly reduced. For example, a function composed of many piece-wise linear segments might have a long description length under a typical continuous parametrization (e.g., the radial basis kernel described below), because it violates the smoothness assumptions of the prior. However, conditional on the change-points between segments, the function can be decomposed into independent parts each of which is well-described by a simple continuous parametrization (see Lee & Yuille, 2006, for a discussion of how this strategy is used by the brain for early vision). If internally structured functions are “natural kinds”, then the combinatorial approach may be a good model of human intuitive function representation.

We describe three kernel parametrizations in the rest of this section. The first two are continuous, differing in their expressiveness. The third one is combinatorial, allowing it to capture complex patterns by composing simpler kernels. For all kernels, we take the standard approach of choosing the parameter values that optimize the log marginal likelihood (see Appendix for details).

### Radial basis kernel

The radial basis kernel is a commonly used kernel in machine learning applications, embodying the assumption that the covariance between function values decays exponentially with input distance:

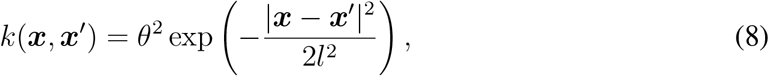

where *θ* is a scaling parameter and *l* is a length-scale parameter determining the speed of the decay over the distance between inputs. This kernel assumes that the same smoothness properties apply globally for all inputs. It provides a standard baseline to compare with more expressive kernels.

### Spectral mixture kernel

The second approach is based on the fact that any stationary kernel^7^ can be expressed as an integral using Bochner’s theorem. Letting 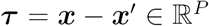, then

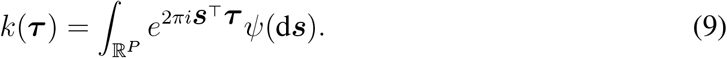

If *ψ* has a density *S*(***s***), then *S* is the spectral density of *k; S* and *k* are thus Fourier duals (Rasmussen & Williams, 2006, see Appendix). This means that a spectral density over the kernel space fully defines the kernel and that furthermore every stationary kernel can be expressed as a spectral density. Wilson and Adams (2013) showed that the spectral density can be approximated by a mixture of *Q* Gaussians, such that

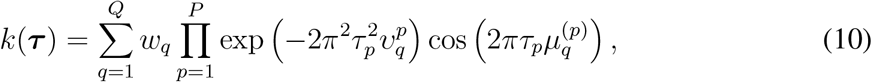

where the *q*th component has mean vector 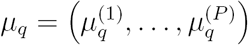 and a covariance matrix 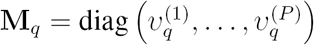. The result is a flexible and expressive parametrization of the kernel, in which complex kernels are approximated by mixtures of simpler ones. Further technical details can be found in Appendix B.

This approach is appealing when simpler kernels (e.g., the radial basis function) fail to capture functional structure. Its main drawback is that because structure is captured implicitly via the spectral density, the building blocks are psychologically less intuitive: humans appear to have preferences for linear (Kalish et al., 2007) and periodic (Bott & Heit, 2004) functions, which are not straightforwardly encoded in the spectral mixture (though of course a mixture can approximate these functions). Since the spectral kernel has been successfully applied to reverse engineering human kernels (Wilson et al., 2015), it is a useful reference of comparison to more structured compositional approaches.

### Compositional kernel

As positive semi-definite kernels are closed under addition and multiplication, we can create richly structured and interpretable kernels from well-understood base components. For example, by summing kernels, we can model the data as a sum of independent functions. Imagine a function that is linearly increasing over time but also shows some seasonal periodicity; then a combination of a linear and a periodic kernel added together might be a good description of that function.

Figure 1 shows an example of how different kernels (radial basis, linear, periodic) can be combined. Our approach, following Duvenaud et al. (2013), is to define a simple grammar over kernels that generates new kernels through summation or multiplication of simpler kernels. Table 1 summarizes the kernels used in our grammar. Given a set of input-output pairs, the task facing the learner is to identify both the function and an underlying parse tree. As with the other kernel parametrizations, the parse tree is chosen to maximize the marginal likelihood.

**Figure 1.**
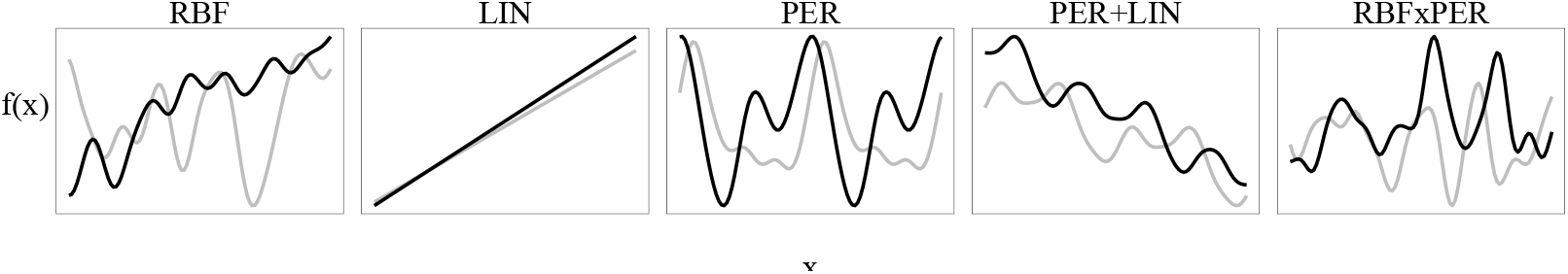
Examples of base and compositional kernels. The base kernels are radial basis (**RBF**), linear (**LIN**), and periodic (**PER**); the composition operators are addition and multiplication. Adding a periodic and a linear kernel creates functions with trends and seasonality. Multiplying a periodic kernel with a radial basis kernel results in more localized periods than a standard periodic kernel would be able to capture.

**Table 1.** Base kernels in the compositional grammar.

| Name | Definition |
| --- | --- |
| Linear | $k(\mathbf{x}, \mathbf{x}') = (\mathbf{x} - \theta_1)(\mathbf{x}' - \theta_1)$ |
| Radial basis | $k(\mathbf{x}, \mathbf{x}') = \theta_2^2 \exp\left(-\frac{(\mathbf{x} - \mathbf{x}')^2}{2\theta_3^2}\right)$ |
| Periodic | $k(\mathbf{x}, \mathbf{x}') = \theta_4^2 \exp\left(-\frac{2 \sin^2(\pi\vert \mathbf{x} - \mathbf{x}'\vert /\theta_5)}{\theta_6^2}\right)$ |

Many other compositional grammars are possible. For example, we could have included a more diverse set of kernels, and other composition operators (e.g., convolution, scaling) that generate valid kernels. However, we believe that our simple grammar is a useful starting point, since the components are intuitive and likely to be psychologically plausible. For tractability, we fix the maximum number of combined kernels to be 3 and do not allow for repetition of kernels in order to restrict the complexity of the inference. The complete set of resulting kernels is shown in Table 2.

**Table 2.**
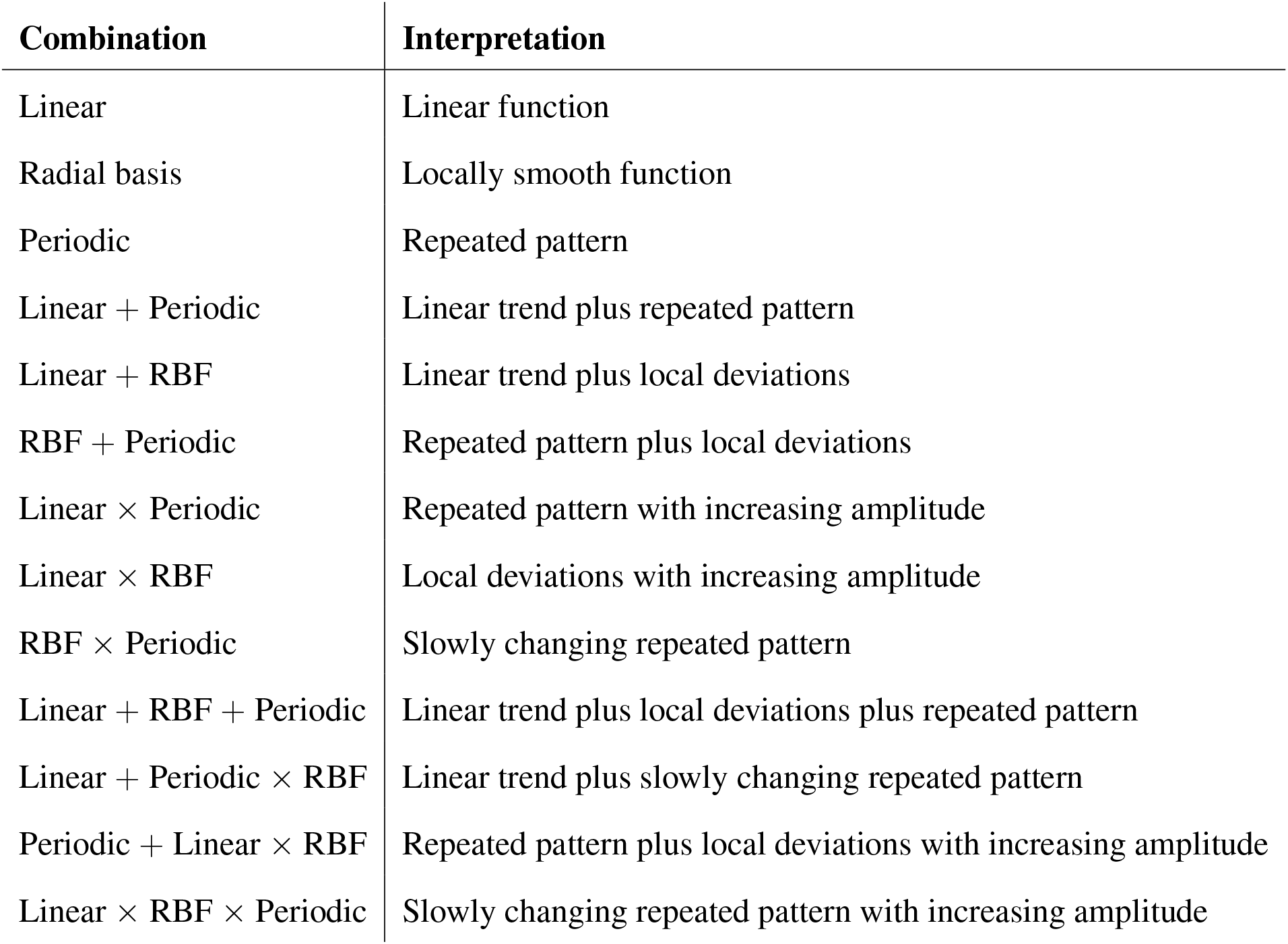
Kernel combinations in the compositional grammar and their interpretations.

| Combination | Interpretation |
| --- | --- |
| Linear | Linear function |
| Radial basis | Locally smooth function |
| Periodic | Repeated pattern |
| Linear + Periodic | Linear trend plus repeated pattern |
| Linear + RBF | Linear trend plus local deviations |
| RBF + Periodic | Repeated pattern plus local deviations |
| Linear × Periodic | Repeated pattern with increasing amplitude |
| Linear × RBF | Local deviations with increasing amplitude |
| RBF × Periodic | Slowly changing repeated pattern |
| Linear + RBF + Periodic | Linear trend plus local deviations plus repeated pattern |
| Linear + Periodic × RBF | Linear trend plus slowly changing repeated pattern |
| Periodic + Linear × RBF | Repeated pattern plus local deviations with increasing amplitude |
| Linear × RBF × Periodic | Slowly changing repeated pattern with increasing amplitude |

We will use this set of kernels as our proposed compositional function class. Given a particular task, we will choose the specific composition from the set that maximizes the marginal likelihood.

## Experiment 1a: Pattern completions of compositional functions

Our first experiment examined whether participants prefer compositional over non-compositional predictions of functions if the ground truth is indeed compositional. Although this mainly tells us whether people make predictions in accordance with the underlying structure (i.e., it is not informative about inductive biases *per se*), Experiment 1b will examine the case where the ground truth is non-compositional.

We used a “pattern completion” paradigm, motivated by prior research on pattern perception as a window into cognitive representations (e.g., Buffart, Leeuwenberg, & Restle, 1981; Kanizsa, 1979). Participants chose among 3 different completions of a partial one-dimensional function.^8^ The candidate completions were generated by the different structure learning approaches described above. Our hypothesis was that if participants have structured, compositional representations of functions, then they should prefer pattern completions generated from the compositional kernel.

### Methods

#### Participants

52 participants (29 women) with an average age of 36.15 (SD = 9.11) were recruited via Amazon Mechanical Turk and received $1 for their participation. The experiment took 5 minutes on average to complete.

#### Design

We preselected 20 different functions^9^ sampled from a Gaussian process parametrized by various compositional kernels within an input space of *x* = [0, 0.1, 0.2, · · ·, 10]. Afterwards, the functional outputs for *x*_learn_ = [0, 0.1, 0.2, · · ·, 7] were used as a training set to which all three approaches were fitted and then used to generate predictions for the test set *x*_test_ = [7.1, 7.2, · · ·, 10]. The hyper-parameters of the radial basis kernel as well as the number of components and the hyper-parameters of the spectral mixture kernel were fitted by optimizing the marginal likelihood given the training set. The best prediction of the compositional kernel approach was found by choosing the composition from the grammar (see Table 2) that produced the best marginal likelihood (following Duvenaud et al., 2013, see Appendix). All of the models were only fitted to the training set (the observed input-output pairs) and used to complete the patterns (predict the functions) for the test inputs.

The different mean predictions were then used to generate 3 plots (one for every kernel approach) that showed the given input as a blue curve and the new predictions (the extrapolation pattern) as a red curve. The procedure was repeated for 20 different compositions, each corresponding to a separate trial.

#### Procedure

Participants were asked to select one of 3 extrapolations (pattern completions) they thought best completed a given function (Figure 2). The position at which a kernel’s predictions appeared was randomized on every trial.

**Figure 2.**
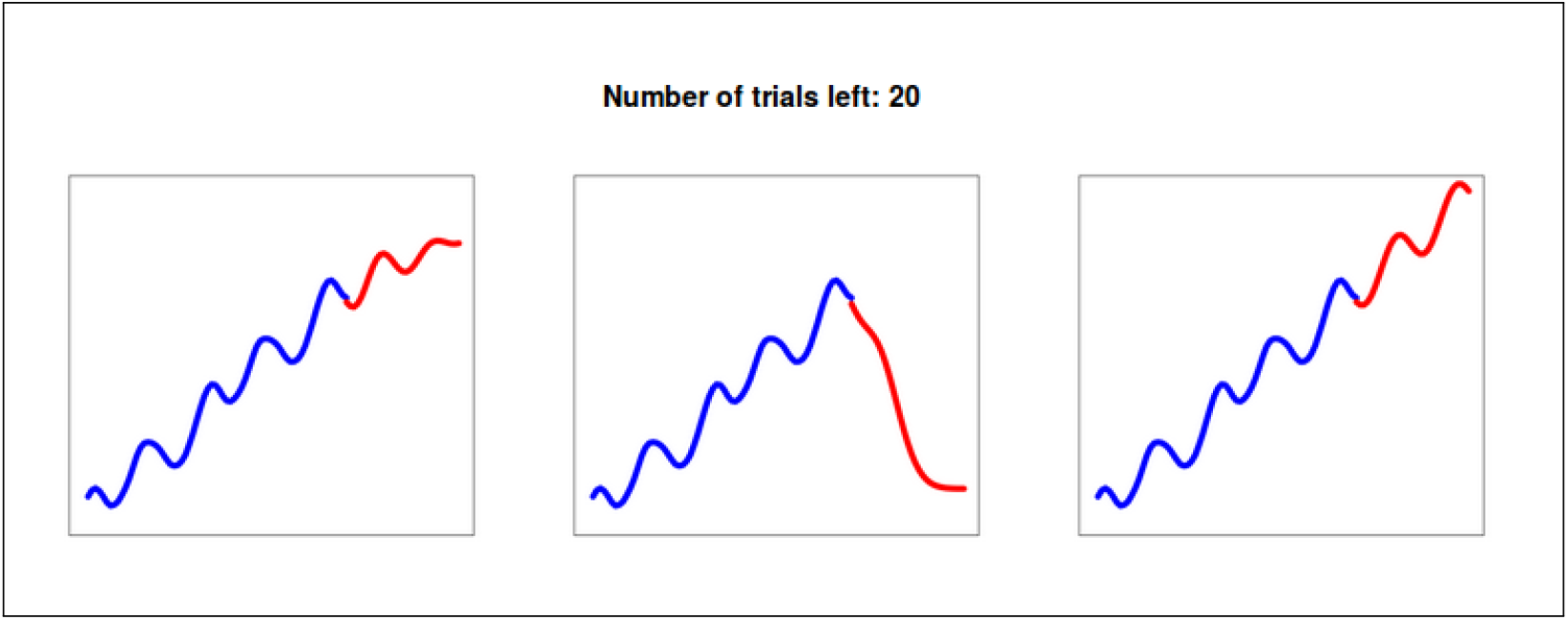
Screen shot of Experiment 1a. Pattern completions (shown in red) were generated by a spectral mixture (left), a radial basis (middle), and a compositional kernel (right).

### Results and discussion

Participants chose the compositional completions on 69% of the trials, the spectral mixture completions on 17%, and the radial basis completions on 14% (Figure 3). Overall, the compositional completions were chosen significantly more often than the other two completions (*χ*^2^(*N* = 1040, *df* = 2) = 591.2, *p* < 0.01). Moreover, assessing for each participant individually whether or not compositional completions were chosen more frequently than the other two based on a *χ*^2^(*N* = 52, *df* = 2)-test, showed that 44 out 52 participants significantly preferred the compositional pattern completions (*α* = 0.05 significance level). The results thus supported our hypothesis that participants will prefer compositional pattern completions when the ground truth is compositional. Using a hierarchal Bayesian model (see Appendix for details) to estimate the posterior probability of choosing the compositional completion led to an estimate of 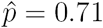 with [0.65, 0.77] as the 95% credible set.

**Figure 3.**
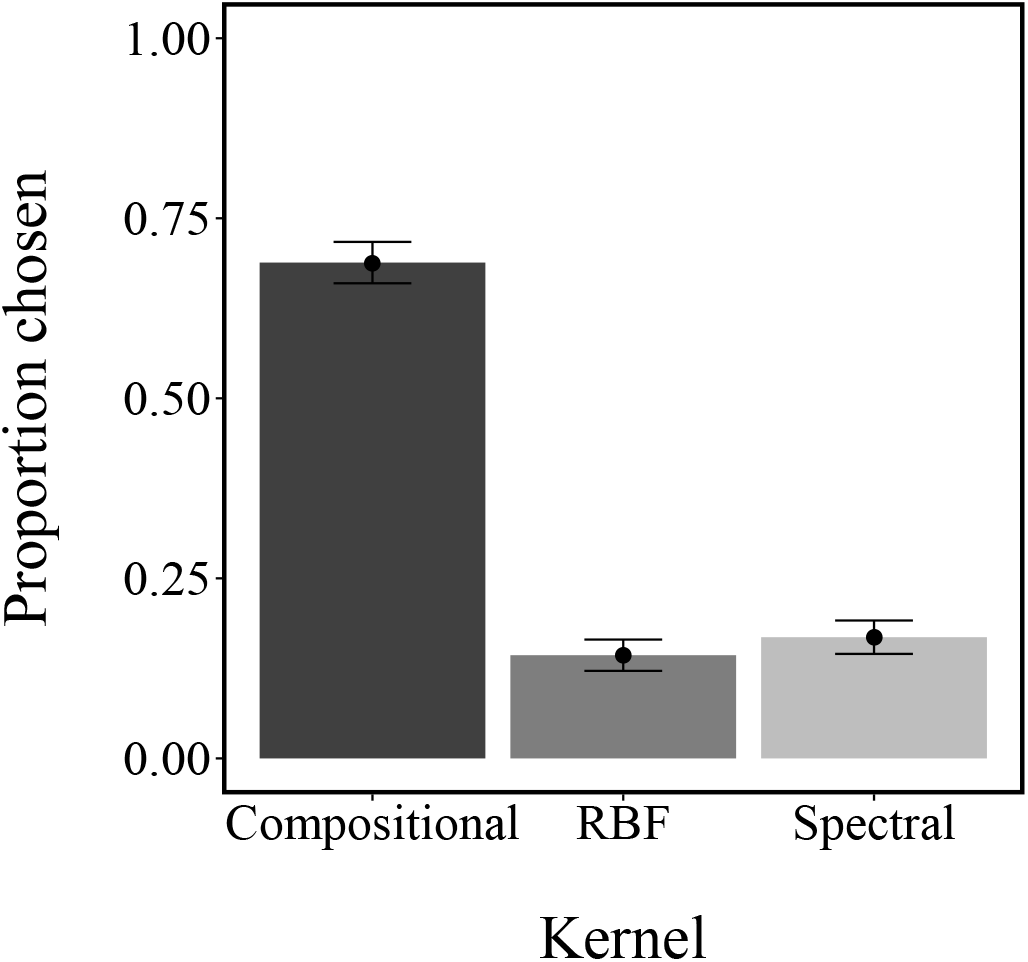
Experiment 1a results: proportion of pattern completion choices for three kernels. Error bars represent the standard error of the mean.

## Experiment 1b: Pattern completions of non-compositional functions

While the results of Experiment 1a suggest a preference for compositionally structured functions, they do not indicate whether humans have an inductive bias for such functions, since the results are perfectly compatible with the possibility that participants adapted to the ground truth structure without a compositional inductive bias. In Experiment 1b, we subject our theory to a stronger test, measuring pattern completion preferences when the ground truth is non-compositional (specifically, functions drawn from a GP with the spectral mixture kernel). If participants prefer compositional completions in this experiment, we can be more confident that the preference for such functions arises from an inductive bias.

### Methods

#### Particpants

65 participants (mean age=30, SD = 9.84, 31 male) were recruited from Amazon’s Mechanical Turk web service and received $0.5 for their participation. The experiment took 4 minutes on average to complete.

#### Design

The design was identical to the one used in Experiment 1a, apart from the fact that in this experiment the underlying functions were sampled (without any further selection) from a GP with the spectral mixture kernel parametrized with a randomly assigned number of components (sampled uniformly between 2 and 5). We also only generated completions for the compositional and the spectral kernel in this experiment. We did not generate completions for the radial basis kernel because in some cases these completions corresponded closely to predictions of the compositional kernel. This correspondence arose due to the fact that there was not much compositional structure for the compositional kernel to capture in samples from the spectral mixture kernel.

#### Procedure

The procedure was as described in Experiment 1a, with the one difference that participants made choices between two (rather than three) completions.

### Results and discussion

As in Experiment 1a, participants chose compositional completions more frequently than non-compositional (spectral mixture) completions (68% vs. 32%, *χ*^2^(*N* = 1300, *df* = 2) = 172.8, *p* < 0.01; Figure 4). Moreover, 41 of 65 participants significantly preferred the compositional over spectral pattern completions at the *α* = 0.05-level. Using a hierarchal Bayesian model to estimate the posterior probability of choosing the compositional completion led to an estimate of 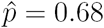 = 0.68 with [0.66, 0.72] as the 95% credible set.

**Figure 4.**
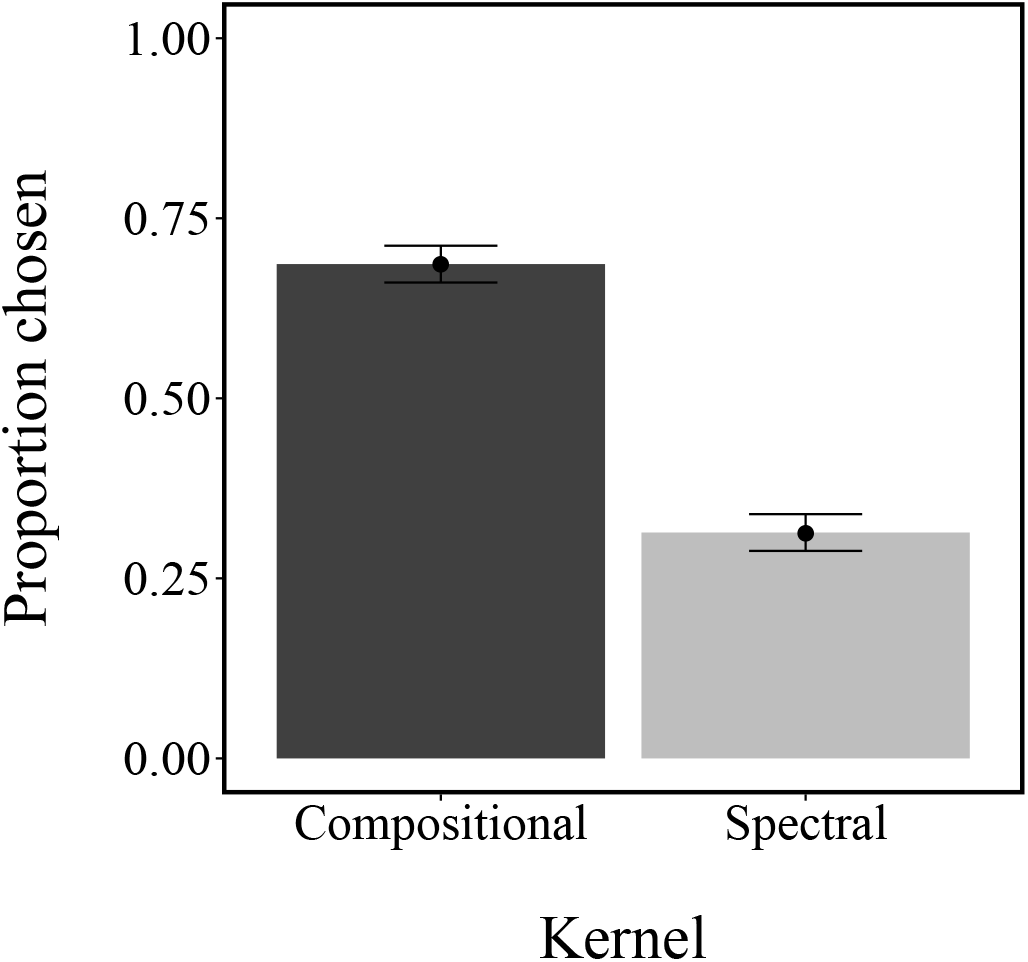
Experiment 1b results: proportions of pattern completion choices. Error bars represent the standard error of the mean.

This finding is consistent with the claim that human inductive biases for functions are compositional—and sufficiently strong to induce preference for compositional completions even when the ground truth is non-compositional.

## Markov chain Monte Carlo with people

In our next set of experiments, we sought to elicit samples from a compositional posterior predictive distribution over functions in order to gain finer-grained insight into the subjective representation of functions. We accomplished this using a technique called *Markov chain Monte Carlo with People* (MCMCP; Sanborn, Griffiths, & Shiffrin, 2010). This technique asks participants to accept or reject proposed hypotheses, effectively simulating a Markov chain whose stationary distribution is the posterior over hypotheses given the presented data. In our experiments, we condition on a training set and ask participants to choose between completions (as in Experiments 1a and 1b); thus the stationary distribution is the posterior distribution over pattern completions.

The basic setup of MCMCP consists of a proposal distribution that generates hypotheses, and an acceptance function that probabilistically accepts or rejects the proposed hypotheses. If the proposal is rejected, the last accepted hypothesis is retained. The proposal distribution is determined by the experimental design, while the acceptance function is a mental construct that we impute to the participant. Under certain assumptions about the acceptance function (given below), Sanborn et al. (2010) showed that this experimental procedure implements a form of Markov chain Monte Carlo known as the Metropolis algorithm. The sequence of hypotheses forms a Markov chain that will eventually (after a “burn-in” period) produce samples from the posterior distribution.

If the proposal distribution is symmetric (the probability of proposing a new state *x*\* from the current state *x* is the same as the probability of proposing *x* from *x*\*), then one psychologically plausible acceptance function is Barker’s acceptance function:

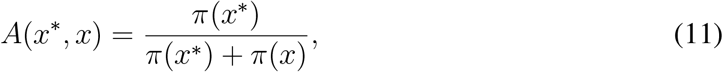

where *π*(*x*) ∝ *P*(*x*). Letting 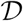 indicate the training set, this leads to the following expression for the acceptance function:

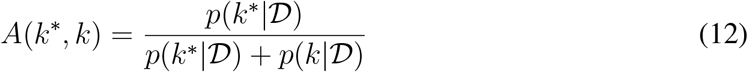

which corresponds to Luce’s choice rule, the most common model for discrete choice distributions in psychology. Thus, under fairly standard assumptions about the choice process, it is possible to elicit samples from the desired distribution, 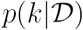, i.e. the posterior distribution over kernels given the data.

This means that MCMCP can be used to gain insights into a restricted distribution (under the assumption that predictions are based on the grammar) over compositional parts by assessing how frequently a given compositional prediction has been preferred over another one. We will therefore use empirical data to assess if participants’ posterior distributions over different kernels approximates sensible compositions.

## Experiment 2a: Compositional ground truth

In the first MCMCP experiment, we sampled the underlying functions from compositional kernels in order to see if the posterior over compositional completions converges to patterns that match the underlying kernel. Using a MCMCP-like design provides us with the opportunity to check if the found restricted posterior distribution corresponds well with the predictions generated by the compositional kernel.

### Methods

#### Design

We generated completions from all possible kernel combinations (up to a maximum of 3 combined kernels), optimizing each kernel’s hyper-parameters on the training set and then generating the completions for the extrapolation set. On each trial, participants chose between their most recently accepted extrapolation and a new proposal.

Eight different functions were sampled from various compositional kernels, the input space was split into training and test sets, and then all kernel combinations (i.e., all kernels described in Table 2) were used to generate completions for the test set. Proposals were sampled uniformly from this set. We mainly focused on combinations containing linear and periodic components, as these provide more interesting structure than the smoothness induced by samples from the radial basis kernel.

#### Participants

51 participants (27 male) with an average age of 32.55 (SD = 8.21) were recruited via Amazon’s Mechanical Turk web service and paid $1. The experiment took 8 minutes on average to complete.

#### Procedure

There were 8 blocks of 30 trials, where each block corresponded to a single extrapolation set. The order of the blocks was randomized for each participant.

### Results and discussion

We calculated the average proportion of accepted kernels over the last 5 trials, as shown in Figure 5. The first 25 trials were excluded to avoid trials on which participants might not have converged yet to their subjective posterior extrapolations, a process commonly called “burn-in”.

**Figure 5.**
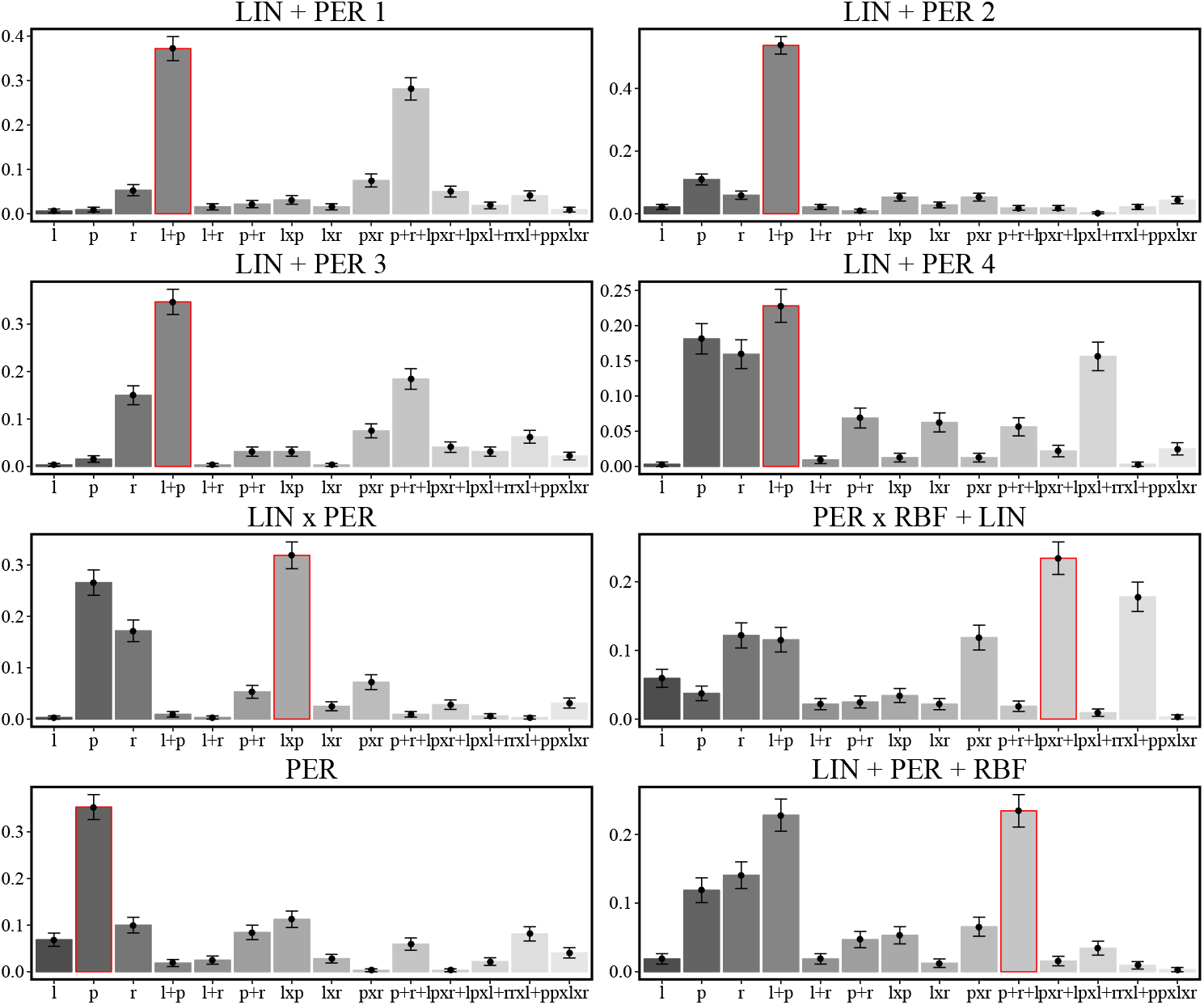
Experiment 1a results: proportions of chosen completions over the last 5 trials. Error bars represent the standard error of the mean. Generating kernel (ground truth) marked in red.

In all cases, participants’ subjective probability distribution over completions placed the greatest mass on the data-generating kernel (marked in red). Furthermore, we observed a strong rank correlation between an approximated posterior probability over completions^10^ and participants’ subjective distribution (*ρ* = 0.91, *p* < 0.01).

This means that the proportions of choices over all compositions produced by the compositional model was very similar to the proportions produced by our subjects.

Thus, participants approximately converged to the underlying posterior when the functions were generated from compositional kernels.

## Experiment 2b: Real-world functions

The second MCMCP experiment assessed what structures people converged to when faced with real-world data, where the ground truth is unknown. We hypothesized that real-world data is often intrinsically compositional (see Duvenaud et al., 2013), and hence human inductive biases may be adapted to such functions.

### Participants

51 participants with an average age of 32.55 (SD = 12.14) were recruited via Amazon Mechanical Turk and received $1 for their participation. The experiment took 7 minutes on average to complete.

### Procedure

We used four real-world time series (Figure 6): airline passenger data, volcano CO2 emission data, the number of gym memberships over 5 years, and the number of times people googled the band “Wham!” over the last 8 years. Some of these functions have been extensively analyzed in the time series modeling literature and all of them showed interesting patterns a priori. Participants were not told any information about the data sets (including input and output descriptions); they were simply shown the unlabeled input-output pairs in the experiment.

**Figure 6.**
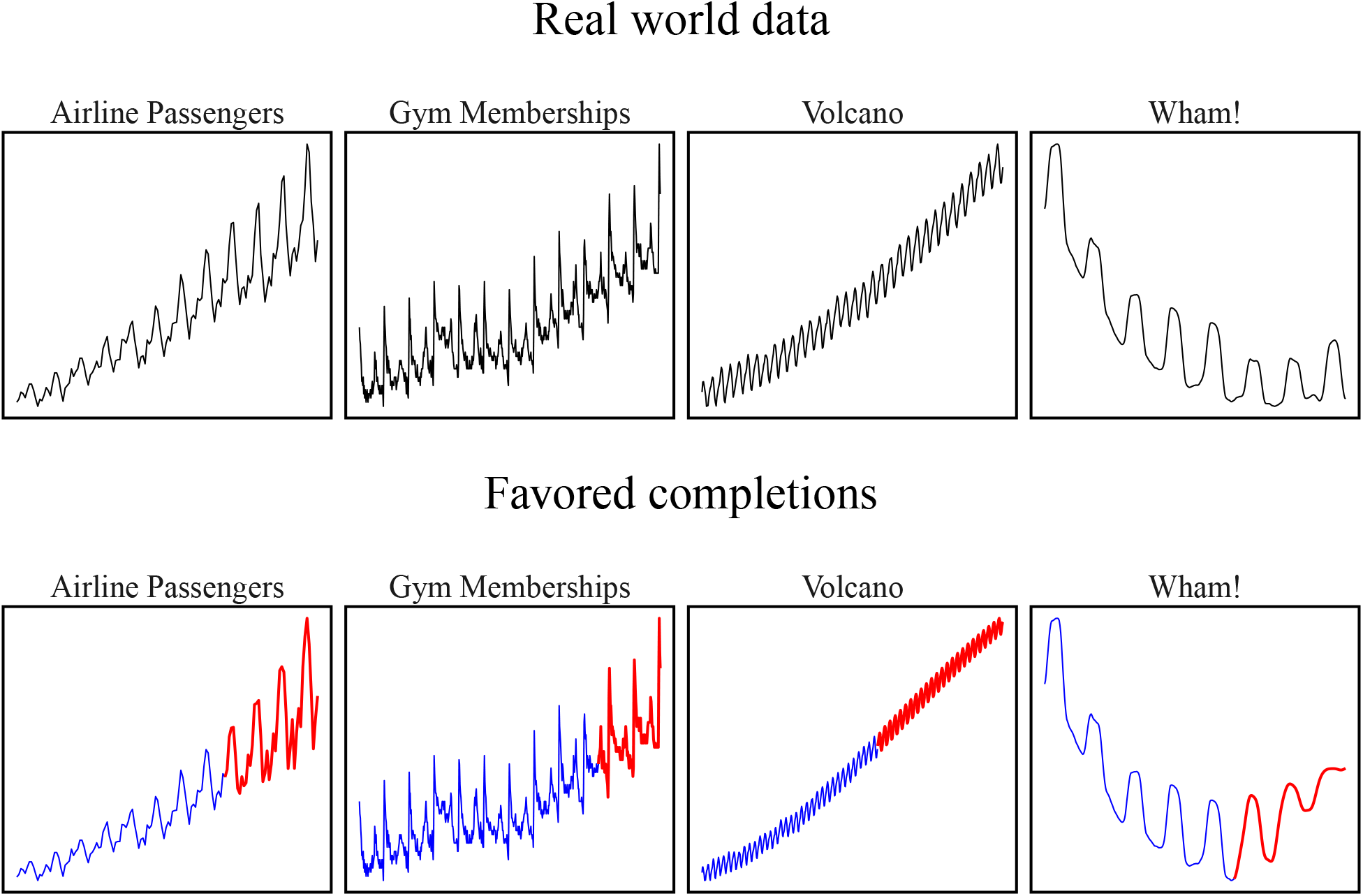
(Top) Real-world data sets used in Experiment 2b. Descriptions and origin of the data were unknown to participants. (Bottom) Participants were shown the region in blue; most frequently selected completions are shown in red. Note that the periodic composition has been adapted by multiplying it with a radial basis function kernel.

The input space was split into training (75% of the data) and test sets (25% of the data), and then all kernel combinations were fitted to the training set and used to generate completions for the test set. Proposals were sampled uniformly from this set. As periodicity in the real world is rarely ever purely periodic, we adapted the periodic component of the grammar by multiplying a periodic kernel with a radial basis kernel, thereby locally smoothing the periodic part of the function.^11^ Apart from the different training sets, the procedure was identical to Experiment 2a.

### Results and discussion

Results (again taken from the last 5 trials) are shown in Figure 7, demonstrating that participants converged to intuitively plausible patterns. In particular, for both the volcano and the airline passenger data, participants converged to compositions resembling those found in previous analyses Duvenaud et al. (2013). The most frequently chosen completions for each data set are shown in Figure 7. The rank correlation between the subjective distributions and the approximated posterior over completions was significantly positive (*ρ* = 0.83, *p* < 0.01), supporting the hypothesis that the compositional pattern completions capture human inferences about functions. This shows again that the proportions of choices over all compositions produced by the compositional model was similar to the proportions produced by our subjects.

**Figure 7.**
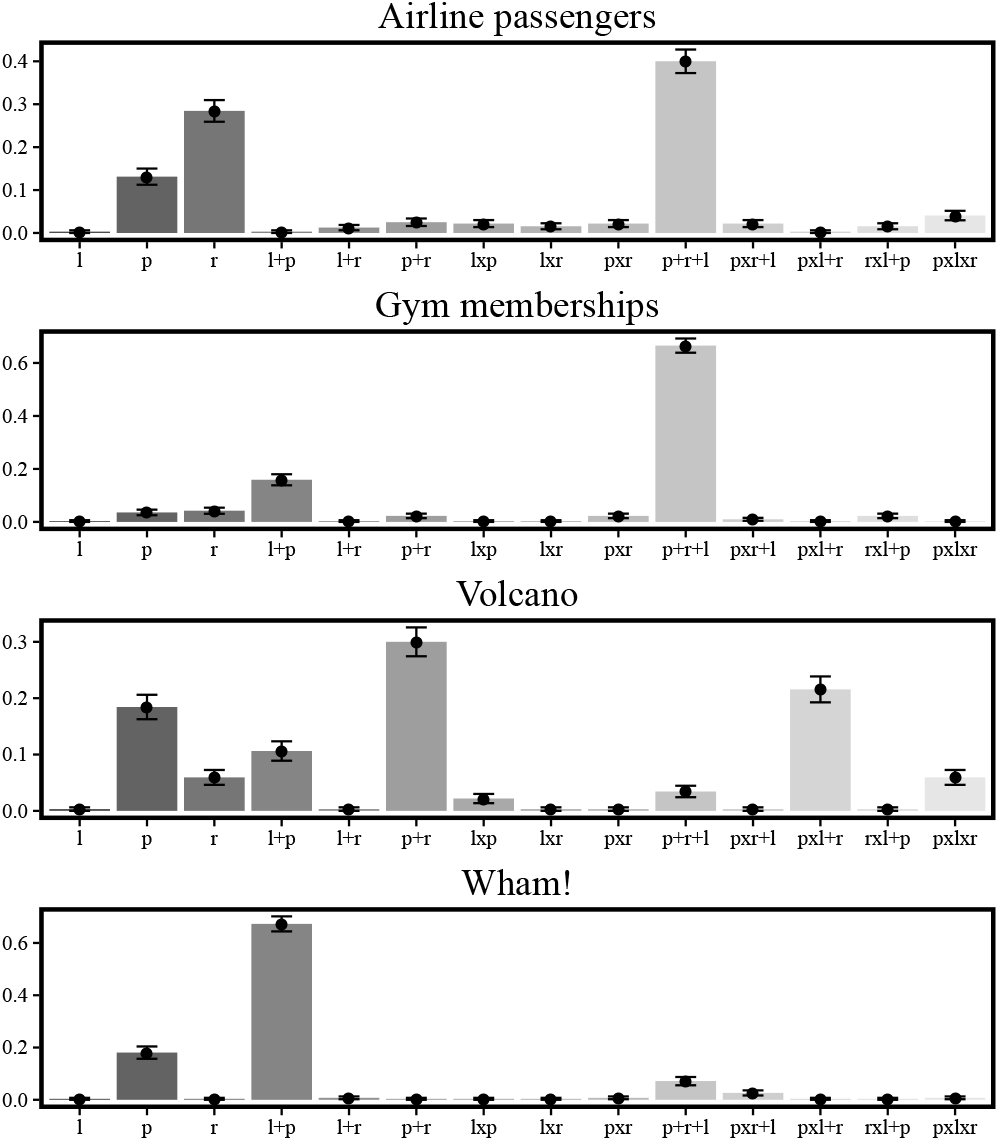
Experiment 2b results: proportions of chosen predictions over last 5 trials.

## Experiment 3: Manual pattern completion

In our previous experiments, we asked participants to make choices between a discrete set of pattern completions. In our next experiment, we measured pattern completion in a less constrained task, by having participants draw the pattern completions manually (see Cox, Kachergis, & Shiffrin, 2012, for related work).

### Methods

#### Design

On each round of the experiment, functions were sampled from the compositional grammar at random, the number of points to be presented on each trial was sampled uniformly between 100 and 200, and the noise variance was sampled uniformly between 0 and 25 and fixed for each function. Finally, the size of an unobserved region of the function (for completion) was sampled to be of a size between 5 and 50. Participants were asked to manually draw the function best describing the observed data and to complete this function within the observed and unobserved regions. A screen shot of the experiment is shown in Figure 8.

**Figure 8.**
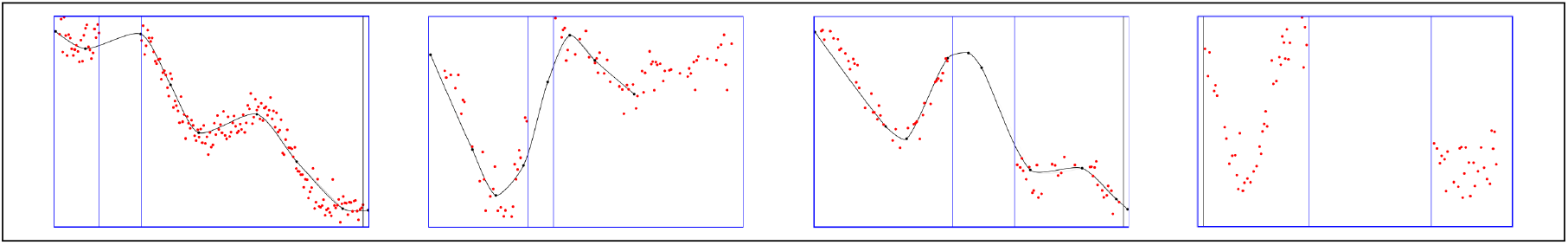
Screen shots of manual pattern completions (Experiment 3). The unobserved region (for completion) is delimited by vertical lines.

#### Participants

36 participants with a mean age of 30.5 (SD = 7.15) were recruited from Amazon Mechanical Turk and received $2 for their participation. The experiment took 12 minutes on average.

#### Procedure

Participants were asked to draw lines in a cloud of dots that they thought best described the given data. To facilitate this process, participants placed black dots into the cloud, which were then automatically connected by a black line based on a cubic Bezier smoothing curve (see Forrest, 1972). The smoothness of the Bezier curve was adapted online to fit the points provided by the participants, thereby making it possible to draw less smooth lines by placing dots closer to each other^12^. Participants were asked to place the first dot on the left boundary and the final dot on the right boundary of the graph. In between, participants were allowed to place as many dots as they liked (from left to right) and could remove previously placed dots. There were 50 trials in total. The dots were sampled from a function that was in turn sampled from a kernel chosen at random out of all possible combinations in our compositional grammar (as shown in Table 2).

### Results and discussion

One concern about our chosen smoothing method was that it might lead to an excessive tendency towards linear interpolations. However, the mean absolute correlation between the input space and participants’ predictions did not significantly differ from the mean absolute correlation between the input space and the true underlying functions. Therefore, the applied smoothing method did not cause participants to prefer linear functions more than required (*t*(36) = 0.09, *p* = 0.92, *d* = 0.02)

We assessed the average root mean squared distance between participants’ predictions (the line they drew) and the mean predictions of each kernel given the data points participants had seen, for both interpolation and extrapolation areas. This means that we only compared the kernel’s predictions given the provided data points with participants’ predictions and did not optimize any parameters in order to achieve better distance measures. As before, we optimized the parameters for the radial basis kernel and the number of components as well as the parameters for the spectral mixture kernel by optimizing the log marginal likelihood. Predictions of the compositional kernel were also generated by choosing the composition that produced the best marginal log-likelihood given the presented dots. Results are shown in Figures 9.

**Figure 9.**
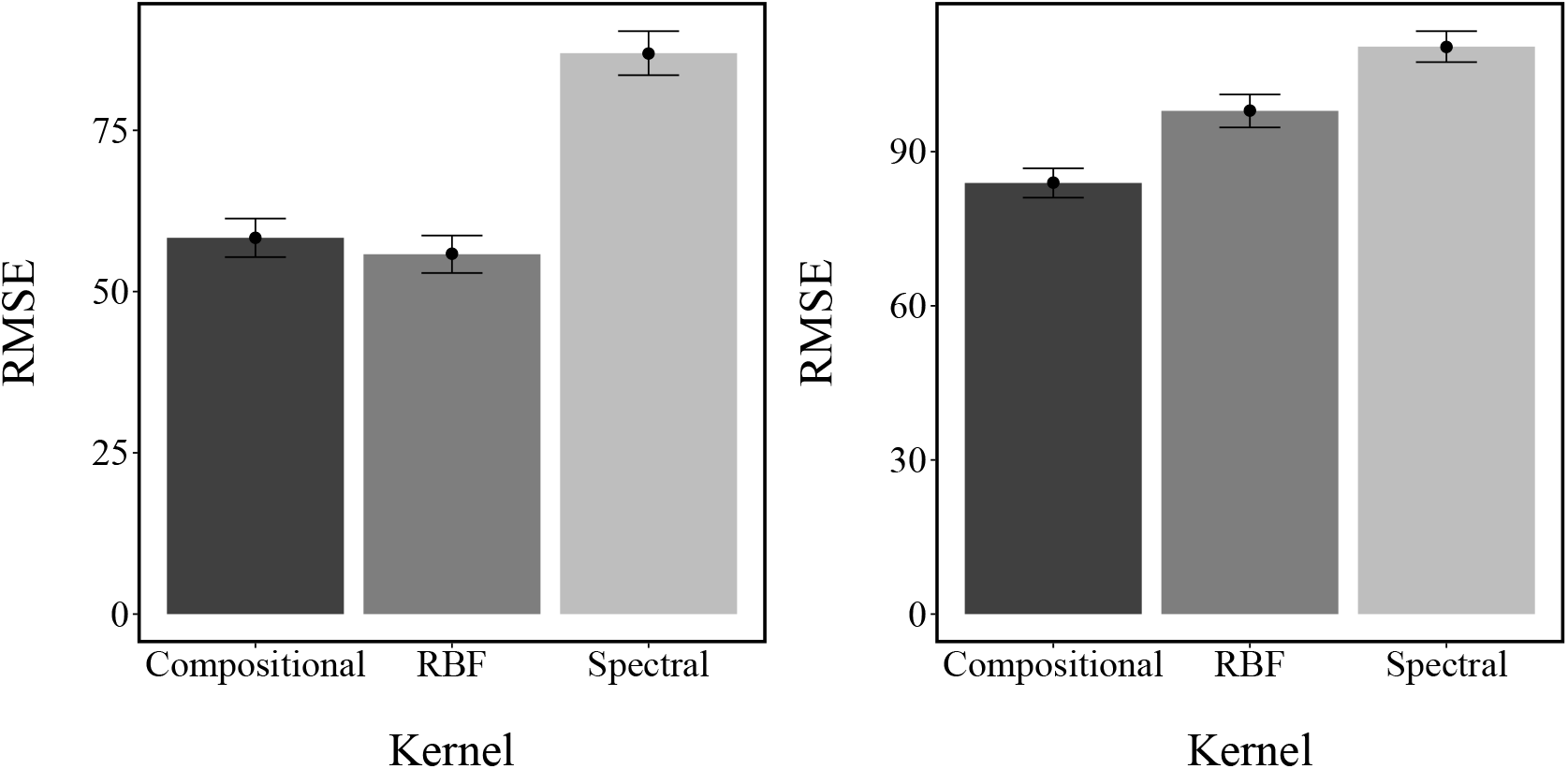
Experiment 3 results: average root mean squared error for interpolation (left) and extrapolation (right) drawings. Error bars show standard error of the mean.

We compared the mean distance between the model predictions and participants’ drawings by performing a hierarchical t-test between the distances produced by the different models while accounting for the nestedness of distance measures within participants and trials; this is essentially the same as performing a mixed-effects regression. All of the mean distances were lower than the distances produced by a random baseline for both interpolation (mean distance of baseline: 142.5) and extrapolation (mean distance of baseline=127.5).

Importantly, the mean distance between predictions and participants’ drawings was significantly higher for the spectral mixture kernel than for the compositional kernel in both interpolation (86.96 vs. 58.33, hierarchical t-test: *t*(676) = −2.28, *p* < 0.05, *d* = 0.35) and extrapolation areas (110.45 vs 83.91, hierarchical *t*-test: *t*(434) = 2.2, *p* < 0.05, *d* = 0.33). The radial basis kernel produced similar distances as the compositional kernel in interpolation (55.8, hierarchical t-test: *t*(649) = −0.55, *p* > 0.05, *d* = 0.03), but predicted participants’ drawings significantly worse in extrapolation areas (97.9, *t*(508) = 1.8, *p* < .05, *d* = 0.17). As extrapolation is normally seen as the best criterion to judge between different models of function learning (DeLosh et al., 1997), the difference in extrapolation judgments provides further evidence for the compositional kernel model.

## Generating comparison functions

One potential concern with the previous experiments is that compositional and non-compositional functions may differ in terms of low-level perceptual characteristics, and these differences (rather than the hypothesized high-level structural differences) are driving the behavioral effects. To address this concern, we need a set of non-compositional functions whose low-level perceptual characteristics (as assessed by the two measures introduced below) are matched with a set of compositional functions.

We generated 500 functions from both compositional and non-compositional kernels. The compositional functions were sampled randomly from our compositional grammar by first randomly sampling a kernel composition and then sampling a function from that kernel, whereas the non-compositional functions were sampled from the spectral mixture kernel, where the number of components was varied between 2 and 6 uniformly. We then calculated the standardized spectral entropy (or “forecastability”) for each function, following Goerg (2013):

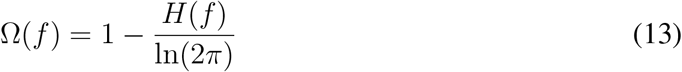

where *H*(·) is the Shannon entropy of the function’s normalized spectral density *S*_*f*_ (see Appendix C for details),

Forecastability can be seen as a measure of how well future output values of a function can be predicted mathematically. It takes values between 0% and 100%, representing the proportional reduction in entropy a given function achieves relative to white noise. Although forecastability technically assumes stationary functions, it has also been shown to produce reliable measures for non-stationary functions in practice (Hyndman, Wang, & Laptev, 2015). Of the 500 non-compositional functions that we generated, we selected those that had a forecastability of higher than 20%, but set the maximum predictability for the compositional kernel functions to be less than 40% and the minimum predictability for the spectral mixture kernel functions to be higher than 40%. This means that theoretically the functions generated from the spectral mixture kernel are more forecastable on average (i.e., contain lower entropy in their spectral density) than the functions sampled from the compositional kernel. Because of this forecastability advantage for non-compositional functions, behavior consistent with the compositional kernel predictions would provide especially strong evidence for our framework.

We verified that all of the functions were matched in terms of low level visual properties, as measured by a similarity measure based on the discrete wavelet Haar transform (Montero, Vilar, et al., 2014, see Appendix C). The basic idea of this similarity measure is to replace the original series by its wavelet approximation coefficients at an appropriate scale, and then to measure similarity between the wavelet coefficients. This measure not only provides a state-of-the art similarity metric between two functions, but also has been used to model human behavior in visual integration tasks (Field, Hayes, & Hess, 1993). Additionally, we validated this measure as a predictor of participants’ similarity judgments when comparing different function in a study described in Appendix E.

We ranked each function based on its wavelet similarity to other functions and selected the top functions (20 compositional and 20 spectral) from this set. These functions, which are used in the experiments reported below, are shown in Figures 10 and 11. We will subsequently refer to these functions as the “matched set.”

**Figure 10.**
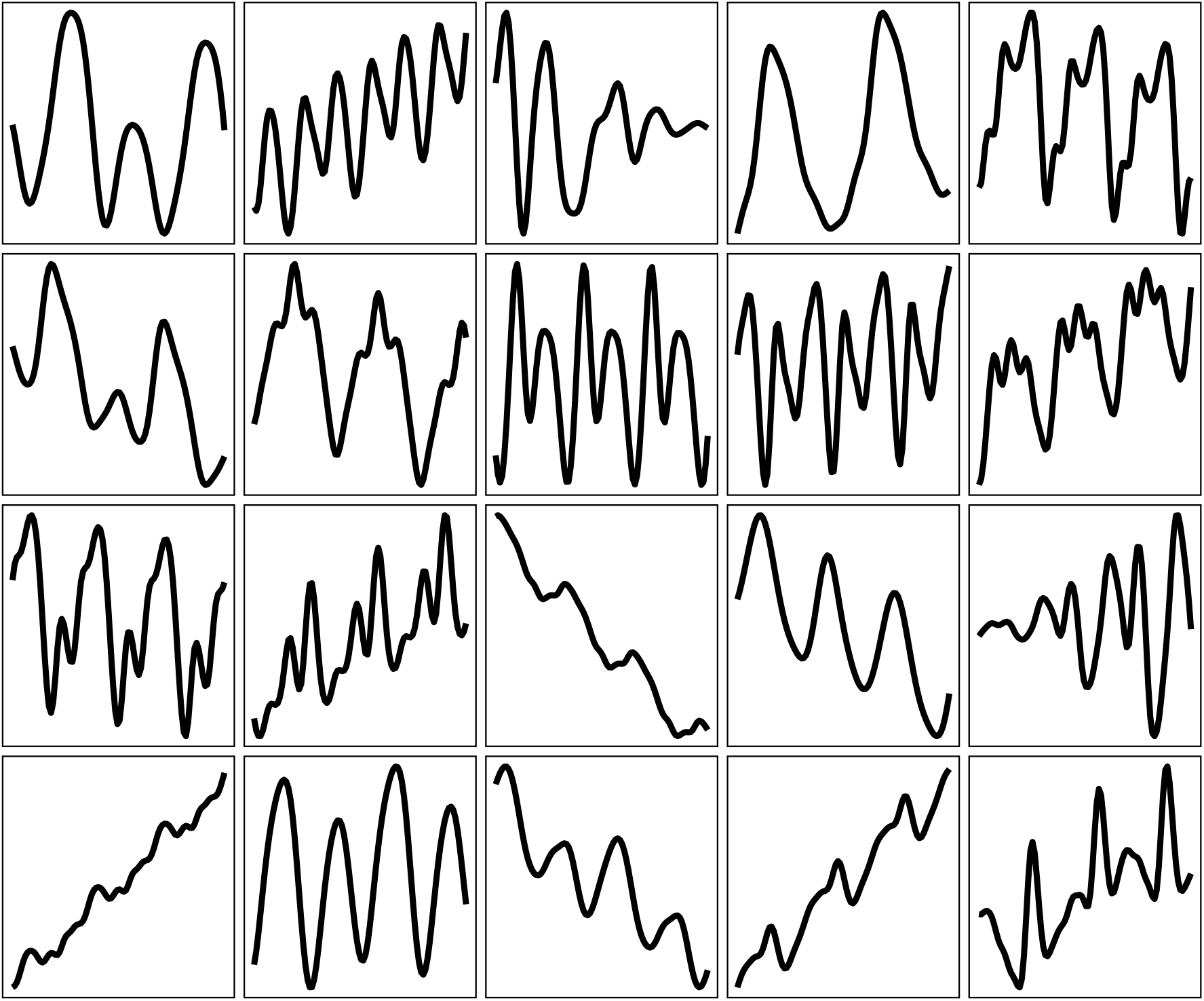
Sampled compositional functions.

**Figure 11.**
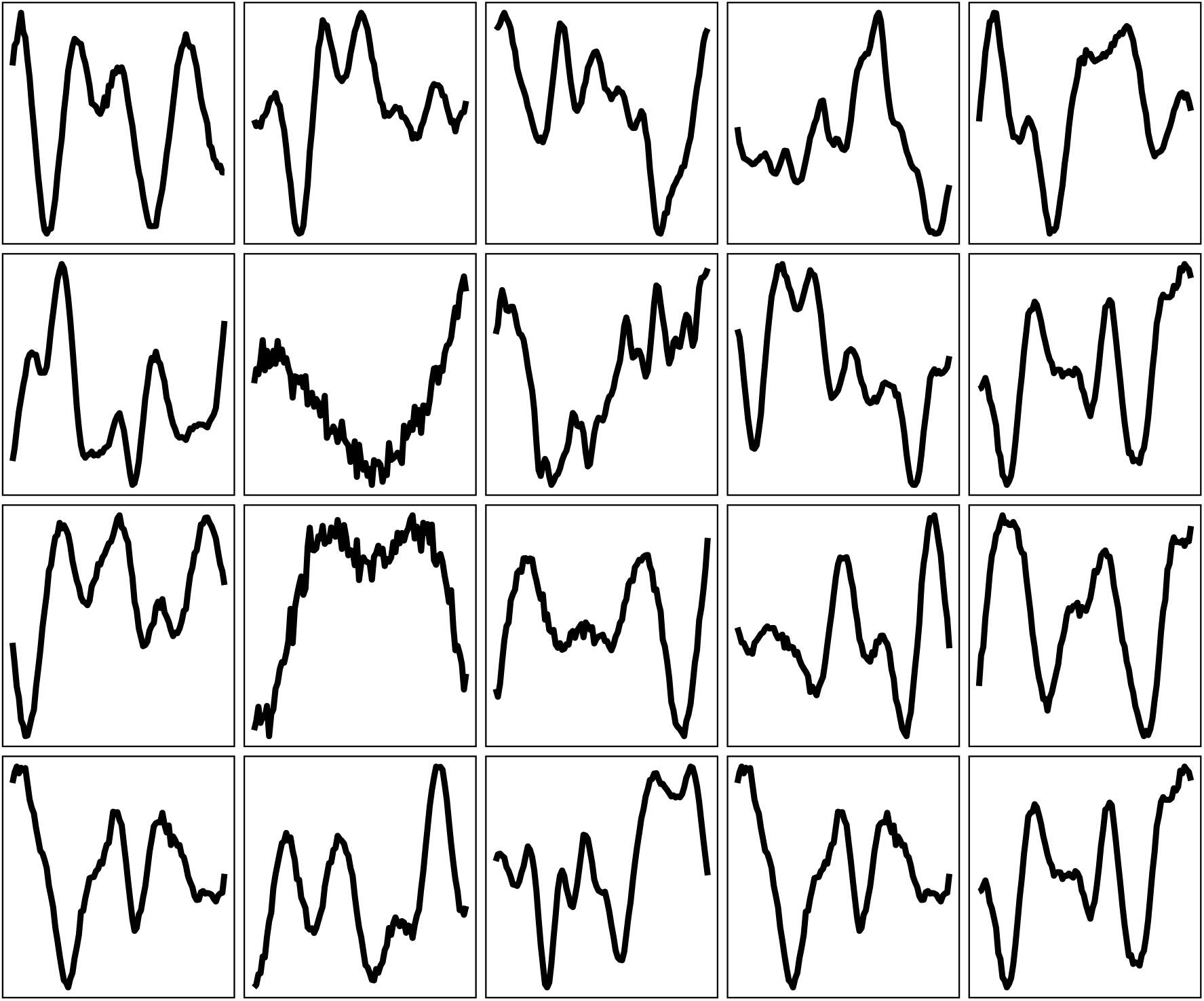
Sampled spectral mixture functions.

## Experiment 4: Assessing predictability

If human inductive biases for functions are inherently compositional, then compositional functions should be perceived as more *predictable*.^13^ In our previous work (Schulz, Tenenbaum, Reshef, Speekenbrink, & Gershman, 2015), we operationalized predictability in terms of generalization error. Formally, predictability is the expected prediction error for a newly-sampled input point from within a (interpolation) set of points. Intuitively, more predictable functions should lead to lower generalization error on new inputs. This analysis identified several key factors determining the predictability of a function; specifically, predictability increases with sample size and smoothness, and decreases with noise.

Here we posit compositionality as another key factor. If inductive biases for functions are compositional, then we expect such functions to be perceived as more predictable. We assess this by collecting both absolute and relative measures of subjective predictability.

### Methods

#### Participants

50 participants (mean age=32, SD=7.2; 32 males) were recruited via Amazon Mechanical Turk and received $0.5 for their participation. The experiment took 9 minutes on average.

#### Procedure

On each trial, one of the “matched” functions defined above was randomly subsampled with a sample size *n*_*j*_ was drawn uniformly from {50,60,…,100}. Participants were asked to judge how well they thought they could predict a newly sampled input point for the function on a scale ranging from 0 (not at all) to 100 (very well). After judging the subjective predictability for all 40 of the matched functions in a randomized order, participants then had to make pairwise comparisons between compositional and non-compositional functions from -100 (function presented on the left is definitely easier to predict) to 100 (function presented on the right is definitely easier to predict). As with the absolute predictability judgments, the sample size *n*_*j*_ was varied randomly, with the constraint that both functions had the same sample size and the position of the functions were counter-balanced over trials.

We did not add noise to the presented functions in order to only assess differences in predictability judgments that can be traced back to differences between the structure of the presented functions. For a treatment of predictability judgments in situations with varying noise levels, we refer the interested reader to our earlier work presented in (Schulz et al., 2015).

Screen shots of the two tasks are shown in Figure 12.

**Figure 12.**
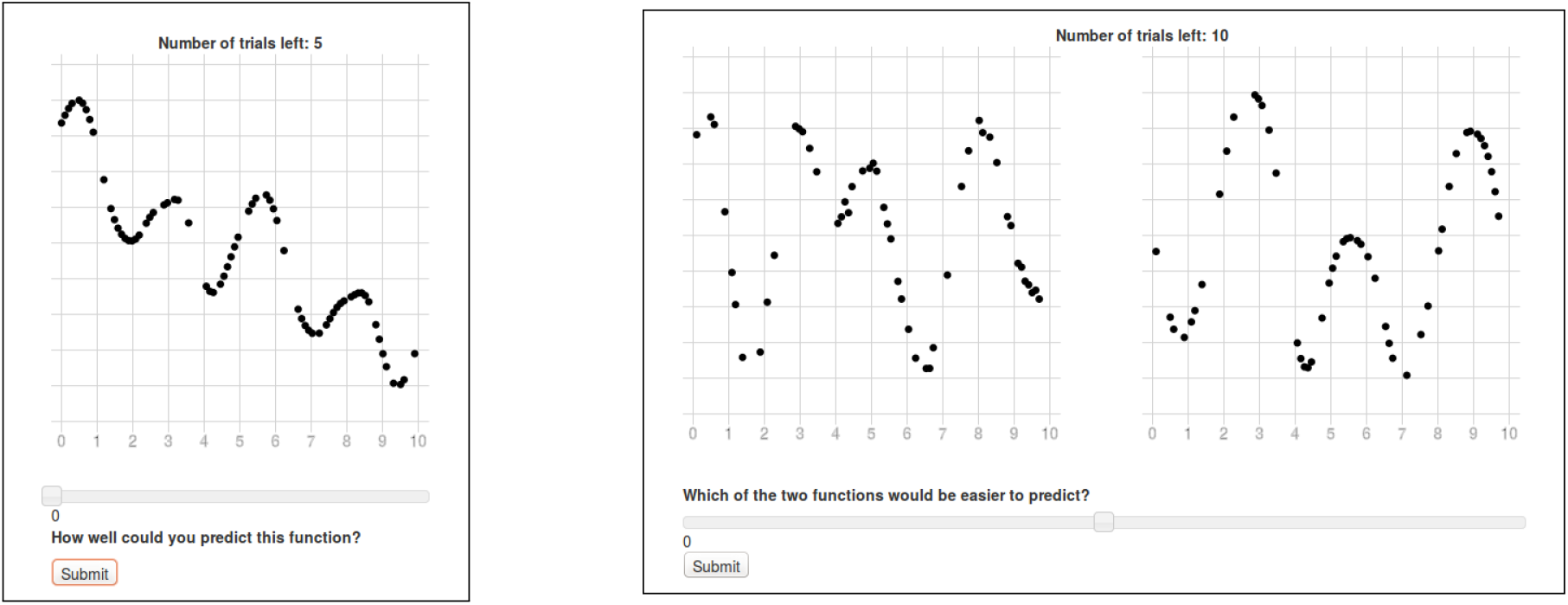
Screen shots of the two predictability judgment tasks (Experiment 4). (Left) Absolute predictability judgments. (Right) Relative predictability judgments.

### Results and discussion

As shown in Figure 13, we found that compositional functions were perceived as more predictable (mean predictability judgment=55.92) than non-compositional functions (mean predictability judgment=37.88 when performing a hierarchical t-test for which judgments were nested within participants and items were treated as a random effect, *t*(31) = 8.35, *p* < 0.001, *d* = 0.73. Moreover, the predictability judgments for compositional judgments were more strongly correlated with sample size (*r* = 0.28, *p* < 0.001) compared to predictability judgments for non-compositional functions (*r* = 0.18, *p* < 0.001; difference test: *z* = 2.36, *p* < 0.05).

**Figure 13.**
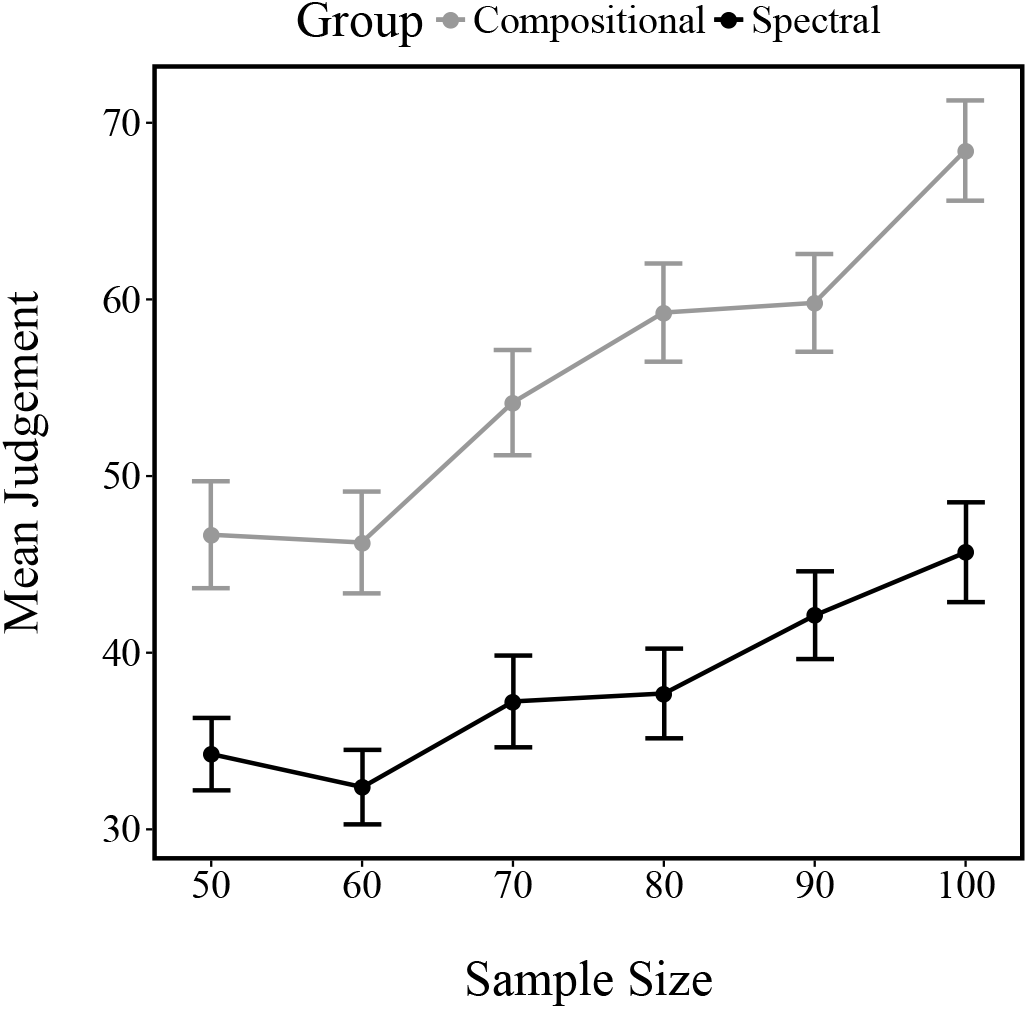
Mean predictability judgments. Error bars represent the standard error of the mean.

To analyze these results further, we fit a hierarchical regression model of subjective predictability (nested within participants) with two predictors: the size of the shown samples and a dummy variable indicating whether or not a presented function was compositional. The fixed effect results of this analysis, summarized in Table 3, demonstrate main effects of both compositionality and sample size.

**Table 3.** Regression model of predictability judgments. Overall model fit: R^2^ = 0.17.

|  | **Estimate** | **Std. Error** | **t-value** | **Pr(>\|t\|)** |
| --- | --- | --- | --- | --- |
| Intercept | 13.06 | 3.50 | 3.72 | 0.0002 |
| Compositional | 17.44 | 1.53 | 11.39 | 0.0000 |
| Sample size | 0.34 | 0.04 | 7.42 | 0.0000 |

The relative predictability judgments tell a similar story (Figure 14). As with the absolute judgments, compositional functions were perceived to be more predictable relative to non-compositional functions (*t*(499) = 13.502, *p* < 0.001, *d* = 0.63), and this difference increased with sample size (*r* = 0.14, *p* < 0.001).

**Figure 14.**
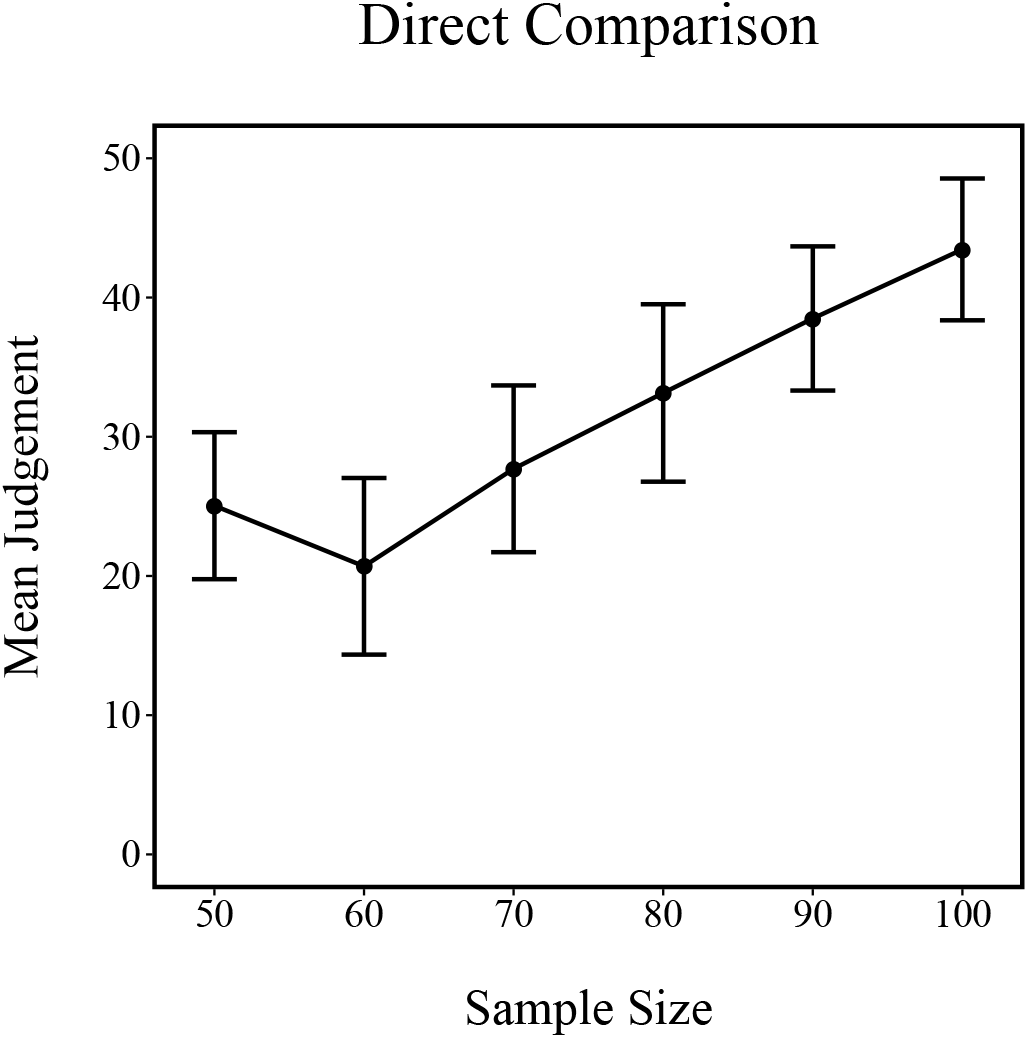
Relative predictability judgments. Positive values indicate that the compositional function was judged to be more predictable than the non-compositional function. Error bars represent the standard error of the mean.

To understand how subjective probability judgments match theoretical predictability, we approximated the generalization error using the average squared error for a randomly sampled new input point. We computed this approximation for every sample that a participant saw under different kernel choices: linear, RBF, spectral mixture, or a compositional kernel. To generate predictions from each kernel, we chose the hyper-parameters (including the composition for the compositional kernel) that maximized the log marginal likelihood. Figure 15 shows the average correlation between each kernel’s generalization error and participants’ predictability judgments, averaged over all trials and participants.

**Figure 15.**
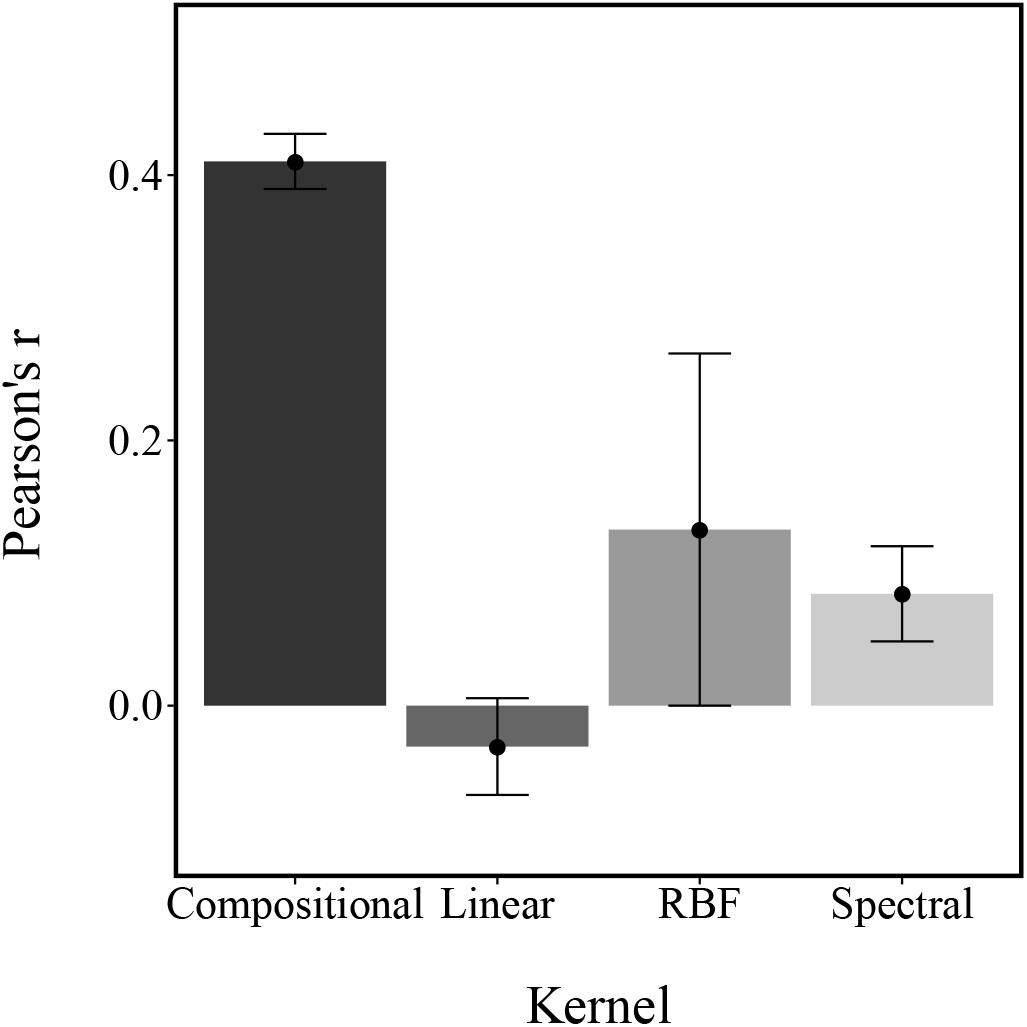
Average correlation between model predictions and participants’ predictability judgments. Predictions were derived from models’ average generalization error. As generalization error and predictability are inversely related, the inverse of the generalization error was used to produce the correlations. Error bars represent the standard error of the mean.

The spectral mixture kernel (t-test against a correlation of 0: *t*(49) = −2.351, *p* < 0.05, *d* = 0.33), the compositional kernel (*t*(49) = −19.73, *p* < 0.001, *d* = 2.79) as well as the RBF kernel (*t*(49) = −4.3, *p* < 0.01, *d* = 0.61) all described participants’ judgments better than chance. The linear (*t*(49) = 0.846, *p* > 0.1, *d* = 0.12) kernel did not predict participants’ judgments better than chance. Crucially, the compositional kernel showed a significantly higher correlation than either the spectral mixture kernel (*t*(49) = −7.86, *p* < 0.001, *d* = 1.57) or the radial basis function kernel (*t*(49) = −9.92, *p* < 0.001, *d* = 1.49).

In summary, compositional functions were perceived as more predictable, even though they were generated to be theoretically less “forecastable” than the non-compositional functions sampled from the spectral mixture kernel. The generalization error produced by the compositional structure learning approach matched participants’ predictability judgments more closely than any of the alternative models.

## Experiment 5a: A short version of the traditional function learning paradigm

The experiments presented so far have mostly focused on visual pattern completions, either by choosing completions in forced choice or MCMCP tasks or by manual pattern completions. However, psychological experiments on function learning have traditionally focused on a different task, in which participants produce output predictions given inputs presented one at a time. For example, Carroll (1963) asked participants to predict the height of one bar given another bar’s height. After each prediction, participants received feedback about how close their prediction was to the actual output.

We sought to compare compositional and non-compositional functions using the traditional function learning paradigm. Additionally, we used this as an opportunity to assess which account of function learning best captures participants’ trial-by-trial learning.

### Methods

#### Participants

46 participants (mean age=31, SD=11; 30 males) were recruited via Amazon Mechanical Turk and received $0.5 for their participation. The experiment took 12 minutes on average to complete.

#### Procedure

Participants were asked to predict an output indicated by the height of a red bar given the current input indicated by the height of a blue bar (Figure 16). On each trial, they saw the current input (a blue bar) and had to indicate their predictions by adjusting the height of an orange bar using a slider. After submitting their prediction, the actual output appeared marked as a red bar directly next to the orange bar. Additionally, participants also saw the actual numbers of the current input, their prediction and –after they submitted their predictions– the output, as well as the difference between their prediction and the actual output. It was explicitly stated to participants that the input is related to the output by an underlying function and that they had to learn that function in order to produce the smallest possible difference between their predictions and the outputs.

**Figure 16.**
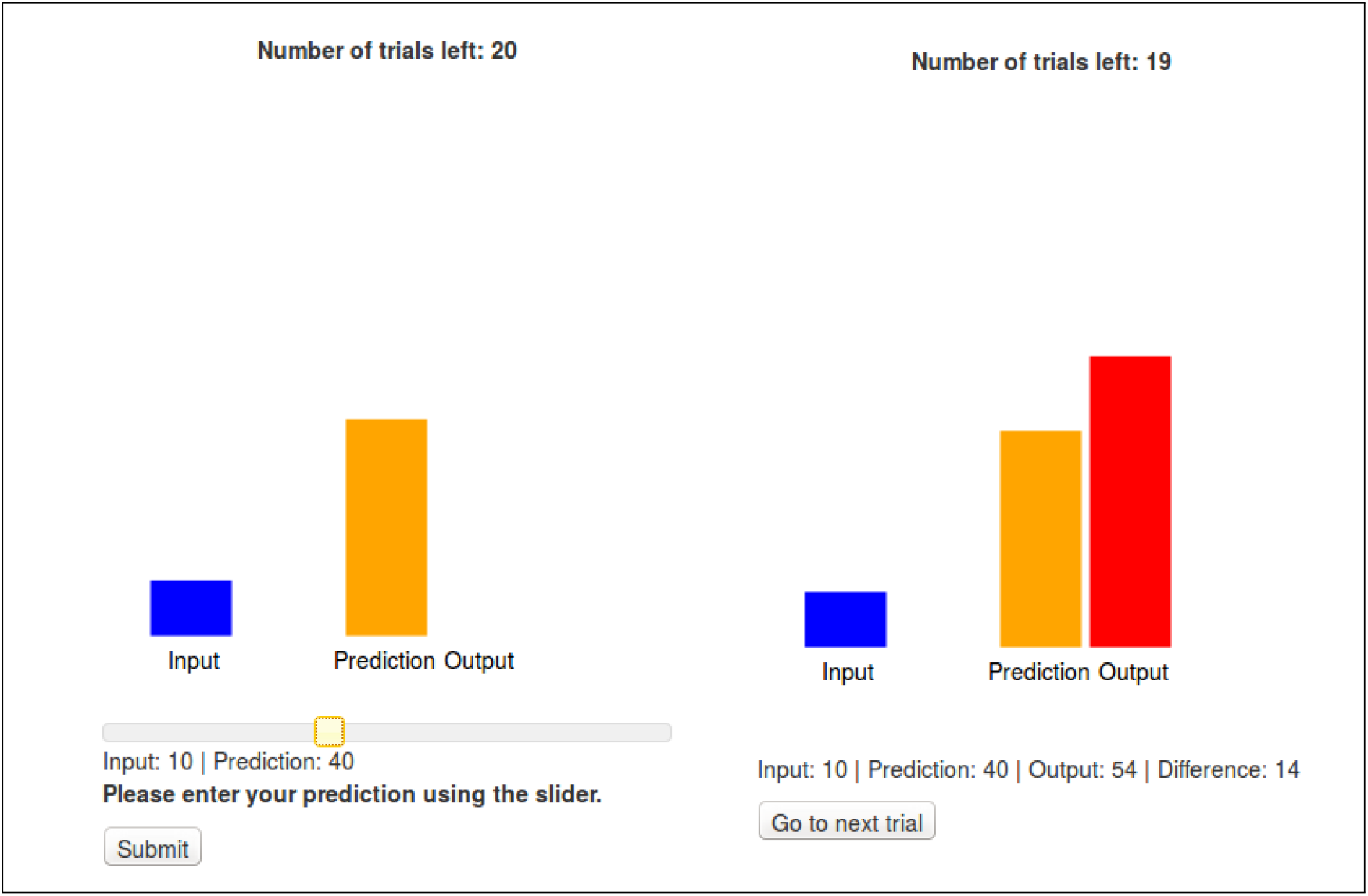
Screen shots of the input-output learning experiment. The height of the blue bar indicates the input. The height of the orange output marks the prediction and can be adjusted by using the slider. After the prediction was submitted, the actual output, marked by the height of the red bar, appeared and participants were told the absolute difference between their prediction and the actual outcome.

Participants learned 4 different functions over 4 blocks. Each block consisted of 20 trials (input-output pairs). The input was randomly sampled to be within 0 and 100, and the output was also transformed to range between 0 and 100. Out of the 4 different functions participants had to learn, 2 were sampled from the set of matched compositional functions and 2 were sampled from the set of matched non-compositional functions.

### Results and discussion

On average, participants’ predictions were only slightly more accurate for the compositional functions than for the non-compositional functions as shown when performing a hierarchical t-test for which predictions were nested within participants and items were treated as a random effect (*t*(34) = 1.6, *p* < 0.1, *d* = 0.18; Figure 17). Performing a linear regression with the absolute difference between participants’ predictions and the actual outputs as the dependent variable and a factor variable encoding whether or not the underlying function was compositional as well as the trial number (how many predictions participants had made before) resulted in the model summarized in Table 5.

**Figure 17.**
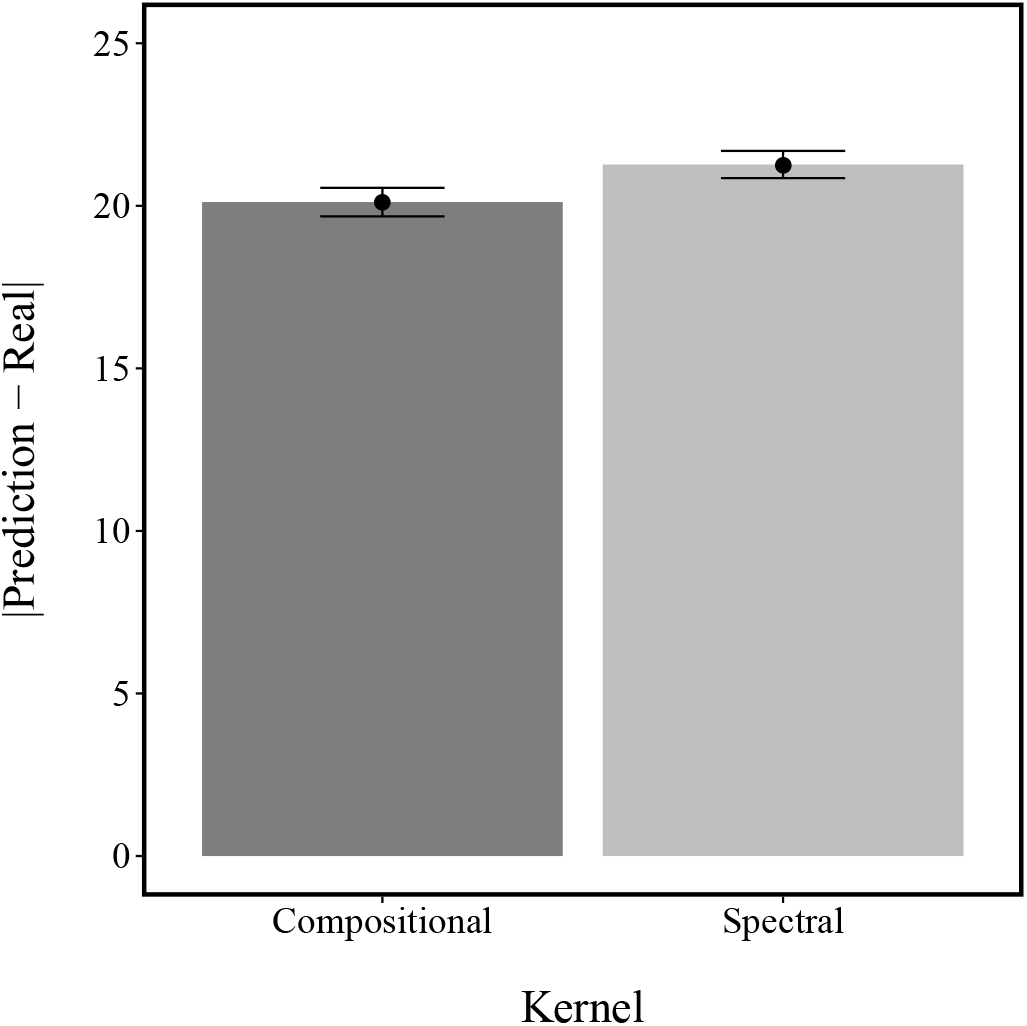
Mean difference between predictions and outcomes for compositional and spectral kernel in Experiment 5a. Error bars represent the standard error of the mean.

**Table 5.**
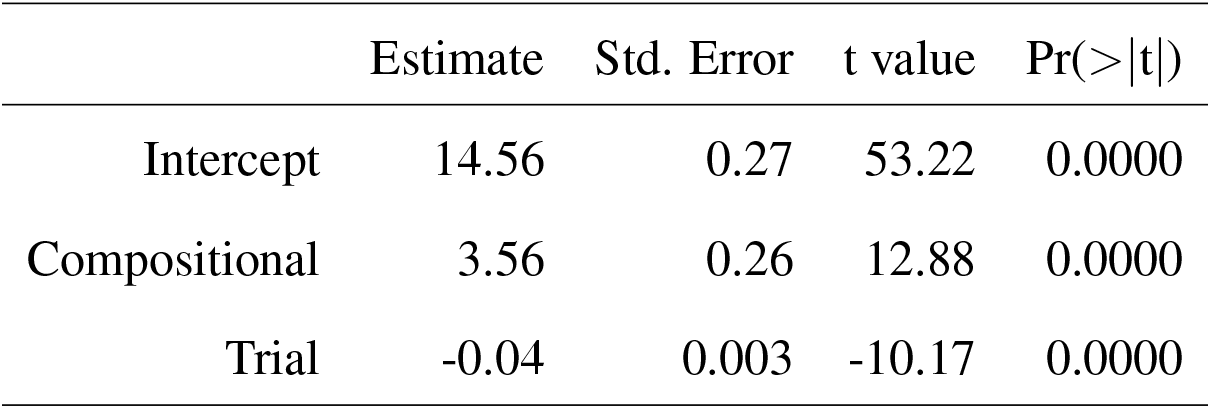
Regression model of the extended traditional function learning paradigm. Overall model fit: R^2^ = 0.14.

|  | Estimate | Std. Error | t value | Pr(>\|t\|) |
| --- | --- | --- | --- | --- |
| Intercept | 14.56 | 0.27 | 53.22 | 0.0000 |
| Compositional | 3.56 | 0.26 | 12.88 | 0.0000 |
| Trial | -0.04 | 0.003 | -10.17 | 0.0000 |

Participants clearly improved over the course of each block, as evidenced by the main effect of trial. Even though compositional functions are on average predicted more easily *(β* = –1.56), this effect turned out to be rather small, perhaps because both compositional and non-compositional functions are non-linear, which are notoriously hard to learn in the traditional function learning paradigm (Byun, 1995) and also because they were created to be quite similar when we designed the matched set.

**Table 4.** Regression model of the traditional function learning paradigm. Overall model fit: R^2^ = 0.14.

|  | Estimate | Std. Error | t value | Pr(>\|t\|) |
| --- | --- | --- | --- | --- |
| Intercept | 23.01 | 0.72 | 32.02 | 0.0000 |
| Compositional | -1.56 | 0.62 | -2.52 | 0.0118 |
| Trial | -0.30 | 0.0535 | -5.65 | 0.0000 |

To further disentangle the different models, we created the trial-by-trial predictions for the linear, compositional, spectral mixture and RBF kernels, and assessed how well each model’s predictions for the *n* + 1 trial after having seen *n* outputs matched participants’ predictions. As shown in Figure 18, the compositional kernel described participants’ predictions best (*r* = 0.54, *p* < 0.01) followed by the linear kernel (*r* = 0.21, *p* < 0.01). The correlation for both the RBF kernel (*r* = 0.03) and the spectral mixture kernel did not differ significantly from zero (*r* = 0.04). Importantly, the compositional kernel predicted participants’ predictions significantly better than the linear kernel (*t*(45) = 9.38, *p* < 0.001, *d* = 1.14).

**Figure 18.**
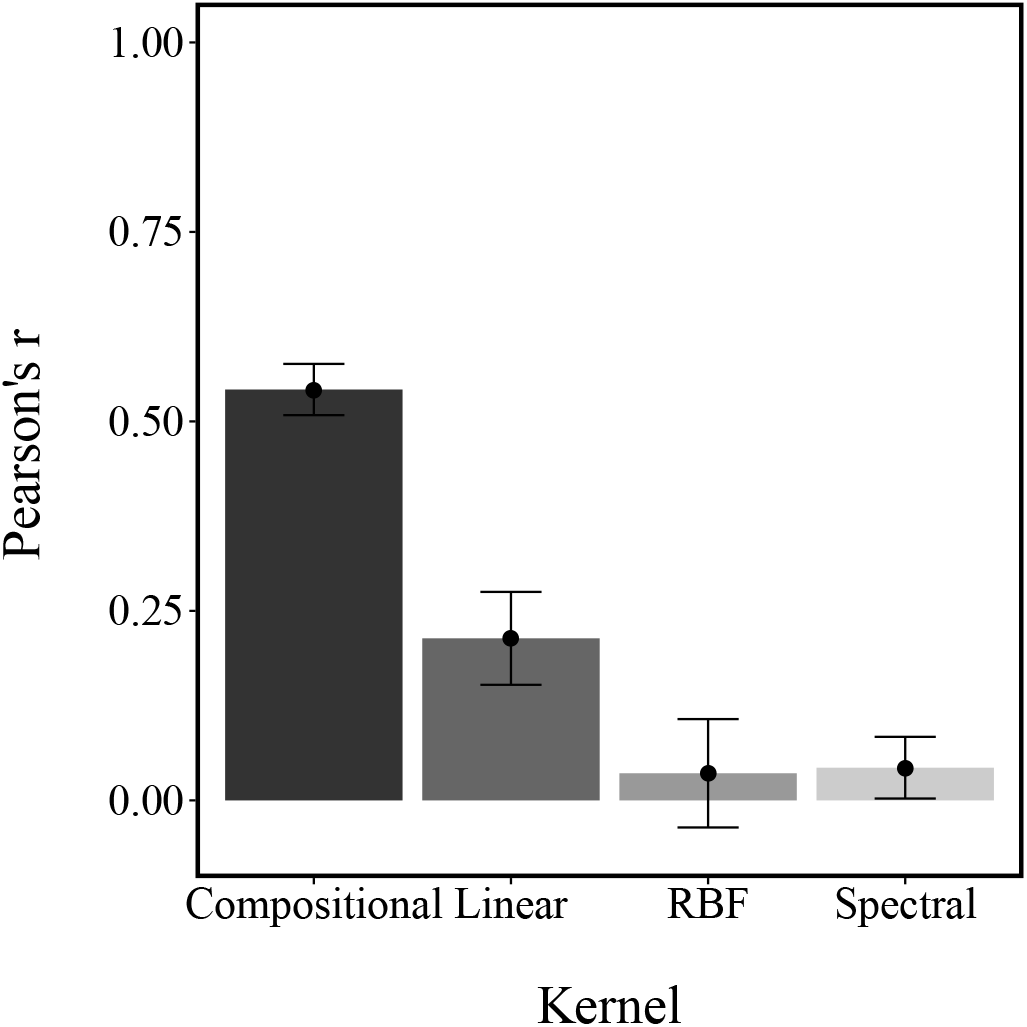
Model fit (Pearson correlation coefficient) for predicting participants’ next predictions given the current input over all trials. Error bars represent the standard error of the mean.

Additionally, more simplistic strategies such as simply matching the height of the input bar (*r* = 0.14, *p* < 0.01) or entering as the next input the output of the previous trial (*r* = 0.15, *p* < 0.01) did not predict participants’ trial-by-trial behavior better than the compositional model. Thus, the compositional kernel appears to provide a better account of trial-by-trial learning in the traditional function learning paradigm over shorter trial numbers.

## Experiment 5b: An extended version of the traditional function learning paradigm

In order to further probe the different accounts of function learning in a trial-by-trial function learning paradigm, we ran another experiment with longer learning horizons as well as an additional stage of both interpolation and extrapolation judgments. This experiment tried to adhere to the design described in DeLosh et al. (1997).

### Methods

#### Participants

89 participants (mean age=38, SD=7.7; 44 females) were recruited from Amazon Mechanical Turk and received $0.5 for their participation as well as a performance-dependent bonus of up to $1.8. The experiment took 35 minutes on average.

#### Procedure

As in Experiment 5a, participants had to predict outputs (indicated by the height of a red bar) given different inputs (indicated by the height of a blue line) by adjusting a slider and thereby changing the height of an orange bar. However, this time the experiment consisted of only one block for which –unknown to participants— one function was sampled from the matched set (i.e., one of the 40 functions described above) and used as the mapping between inputs and outputs. Participants were told that they had to learn and predict how different inputs relate to different outputs over 120 trials.^14^ After the 120 trials were over, participants were asked to make predictions for 30 more trials but without receiving immediate feedback about the output after each prediction. We divided the input space into equidistant points from *x* = [0, 1, 2, 3,…, 100]. For the first 120 trials, participants had to predict inputs for values sampled randomly within the boundaries *x* = [20, 80]. For the 30 remaining trials, participants had to predict inputs for 10 inputs from within the previously learned range that they had not seen before, i.e. interpolation predictions, 10 for inputs sampled from the range [0, 20] and 10 sampled from the range [80, 100], i.e. extrapolation predictions. Participants gained between 0 and 10 points for the quality of their predictions (measured by the absolute distance to the output) over the first 120 trials, and between 0 and 20 for the last 30 trials. After participants had finished the experiment, their total points were converted into money such and they received $0.10 for every 100 points gained during the experiment.

### Results and discussion

Participants gained $2.02 on average. This time, the spectral functions were predicted better than the compositional functions over all of the 120 trials (hierarchical t-test: *t*(87) = −2.38, *p* < 0.05, *d* = 0.52). This can also be seen when performing a similar regression as in Experiment 5a.

This means that —over extended experience— participants learn to predict inputs from within the provided sets slightly better for the spectral than for the compositional functions. This is not that surprising as these functions were set up to be more forecastable when we designed the “matched set”. However, a crucial comparison is then whether compositional or spectral functions are easier to predict both within the interpolation and extrapolation set. We found that whereas spectral functions are easier to predict in interpolation judgments (*t*(87) = −1.89, *p* < 0.1, *d* = 0.41), compositional functions are easier to predict in extrapolation judgments (*t*(87) = 2.1, *p* < 0.05, *d* = 0.46). This means that compositional structure is more difficult to learn than the relatively unstructured but smooth spectral functions but leads to better out-of-sample predictions over all (Figure 19).

**Figure 19.**
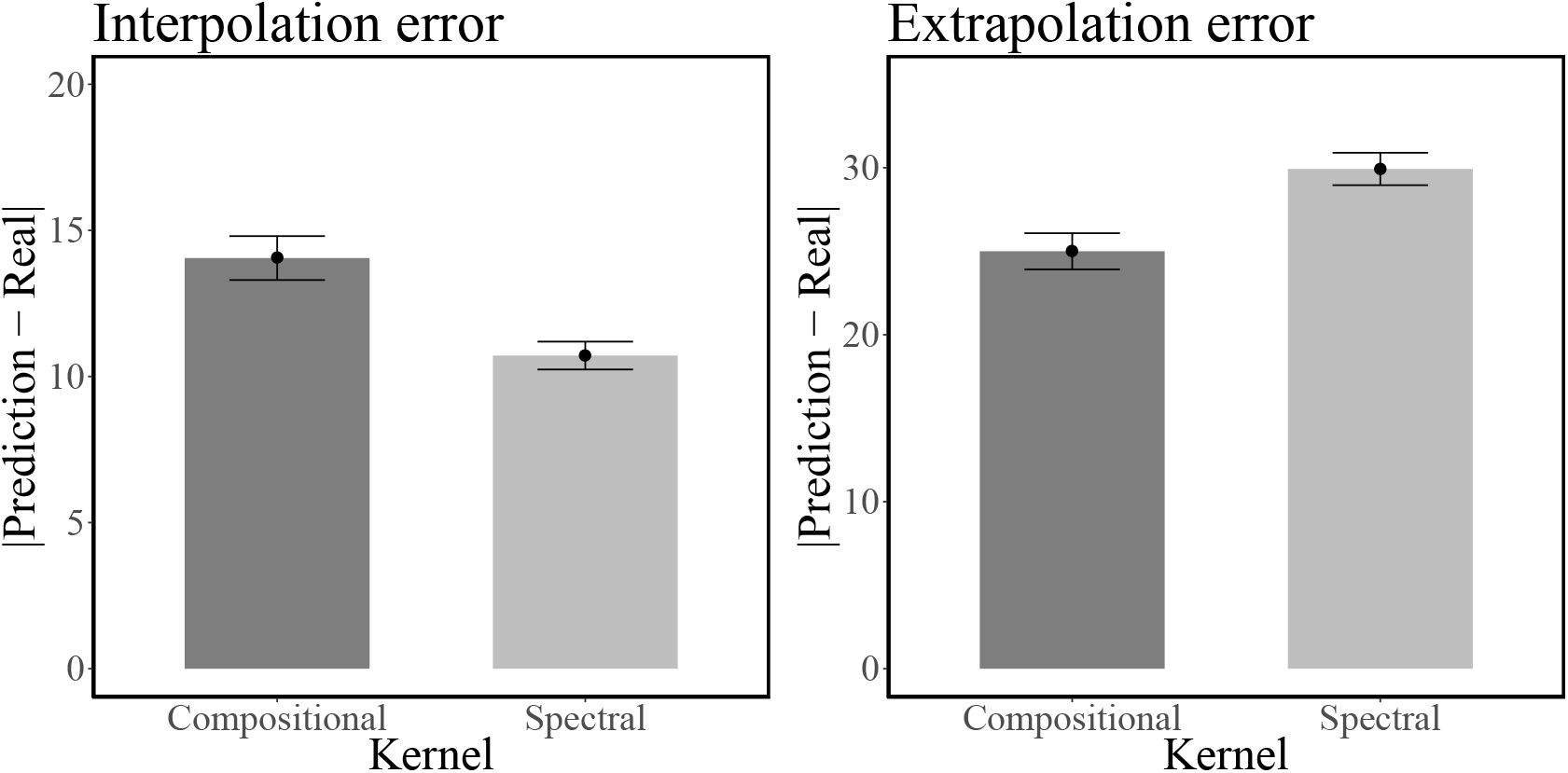
Experiment 5b results: average error for interpolation (left) and extrapolation (right) predictions. Error bars show the standard error of the mean.

Next, we again created trial-by-trial predictions for the linear, compositional, spectral mixture and RBF kernels, and assessed how well each model’s predictions for the *n* + 1 trial after having seen *n* outputs matched participants’ predictions. Moreover, we also assessed how well each of the kernels was able to capture participants’ predictions for both the interpolation and extrapolation trials. Results are shown in Figure 20.

**Figure 20.**
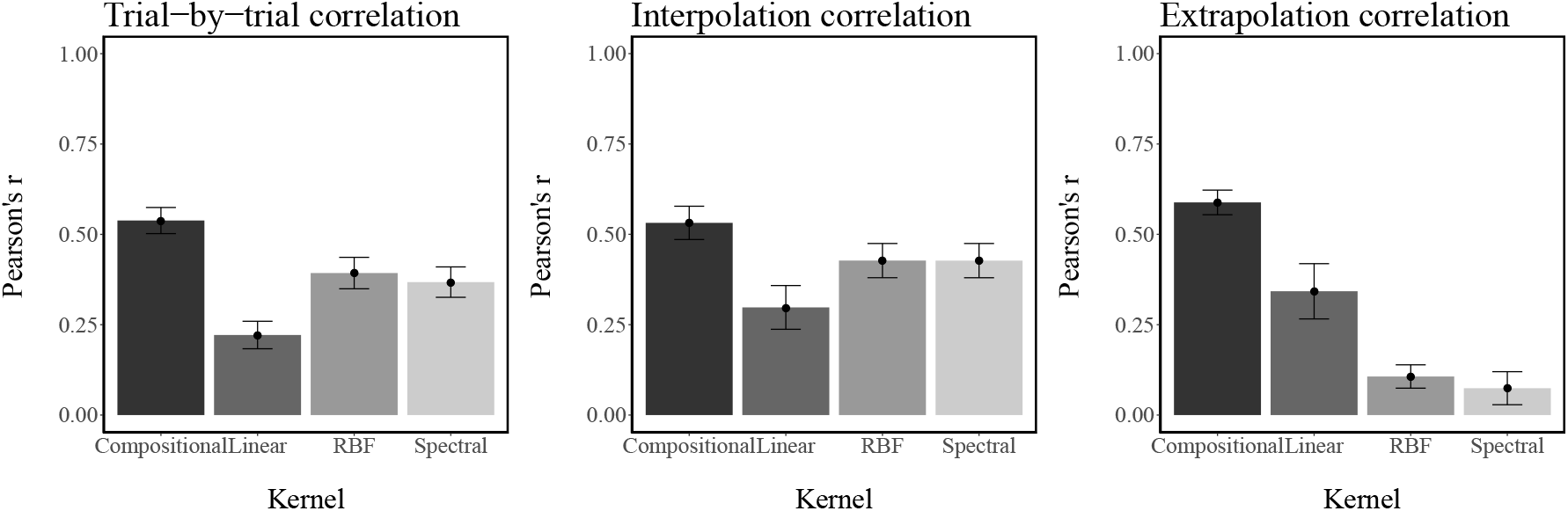
Experiment 5b results: average correlation between model and participants’ predictions for trial-by-trial learning (left), interpolation (center) and extrapolation (right). Error bars show the standard error of the mean.

Overall, all models generated better trial-by-trial predictions than what would be expected at chance level. For the trial-by-trial predictions, the compositional kernel generated better predictions than the linear kernel (*t*(87) = 6.99, *p* < 0.001, *d* = 0.74), the RBF kernel (*t*(87) = 6.85, *p* < 0.001. *d* = 0.73) as well as the spectral mixture kernel (*t*(87) = 8.23, *p* < 0.001, *d* = 0.87). Both the RBF (*t*(87) = 3.19, *p* < 0.01, *d* = 0.34) as well as the spectral mixture kernel (*t*(87) = 2.73, *p* < 0.01, *d* = 0.29) led to better predictions than the linear kernel. A similar picture emerged for the interpolation predictions during the test trials, where the compositional kernel again generated better predictions than the linear (*t*(87) = 5.11, *p* < 0.001, *d* = 0.54), the RBF (*t*(87) = 4.84, *p* < 0.001 *d* = 0.52), or the spectral mixture kernel (*t*(87) = 4.85, *p* < 0.001. *d* = 0.51). As before, both the RBF ((87) = 2.36, *p* < 0.05, *d* = 0.25) as well as the spectral mixture kernel (*t*(87) = 2.36, *p* < 0.05, *d* = 0.25) led to better predictions than the linear kernel. The effects for interpolation predictions were generally smaller than for the trial-by-trial predictions.

Interestingly, some of the results reversed when comparing performance for the extrapolation predictions, where the linear kernel led to better performance than either the RBF (*t*(87) = 3.16, *p* < 0.01, *d* = 0.34) or the spectral mixture kernel *t*(87) = 3.00, *p* < 0.01, *d* = 0.32). However, the compositional kernel still performed better than the RBF (*t*(87) = 14.34, *p* < 0.001, *d* = 1.53), the spectral (*t*(87) = 10.80, *p* < 0.001, *d* = 1.15), or the linear (*t*(87) = 3.91, *p* < 0.001, *d* = 0.42) kernel. The finding that the compositional kernel can match both interpolation and extrapolation judgments resonates well with previous studies. For example, DeLosh et al. (1997) proposed a model that applied smooth trial-by-trial and interpolation learning and simple linear rules for extrapolation. As the compositional kernel can approximate both smooth and linear function, it is able to capture both behavioral phenomena (as also proposed by Lucas et al., 2015).

## Experiment 6: Assessing numerosity

In our next series of experiments, we explore the broader implications of compositional function representation for 3 domains of cognition: numerosity perception, change detection, and short-term memory. We begin with a standard numerosity judgment task, in which participants estimate the number of dots appearing briefly on a screen. Zhao and Yu (2016) showed that, in the absence of explicit grouping cues, structured configurations of dots led to lower estimates compared to random configurations. This effect of structure on perceived numerosity appears to arise from the fact that dots belonging to structured configurations are more likely to be perceptually grouped together; these groups then become the units which are enumerated. In a related study, Zhao, Ngo, McKendrick, and Turk-Browne (2011) showed that the accuracy of numerosity judgments improves when statistical regularities are removed, indicating a direct relationship between statistical learning and numerosity perception.

If compositional functions are perceived as more “structured” than non-compositional functions (as suggested indirectly by our predictability experiment), then a natural hypothesis is that dot configurations generated from compositional functions will be perceived as less numerous. We tested this prediction using a standard numerosity judgment task.

### Methods

#### Participants

91 participants (mean age=37.84, SD=7.87, 48 female) were recruited via Amazon Mechanical Turk and received $0.5 for their participation. The experiment took 7 minutes on average to complete.

#### Procedure

Each participant completed a total of 40 trials. On each trial, participants were presented with a function drawn from the set of matched functions, discretely sampled into 100 equidistant gray dots on the screen. This configuration was displayed for one second, after which the dots vanished. Out of the 100 dots, *n* = [5, 6, 7,…, 15] randomly selected dots were marked as red. Participants were asked to estimate the number of red dots (between 0 and 20) after all the dots had vanished. A screen shot of the experiment is shown in Figure 21.

**Figure 21.**
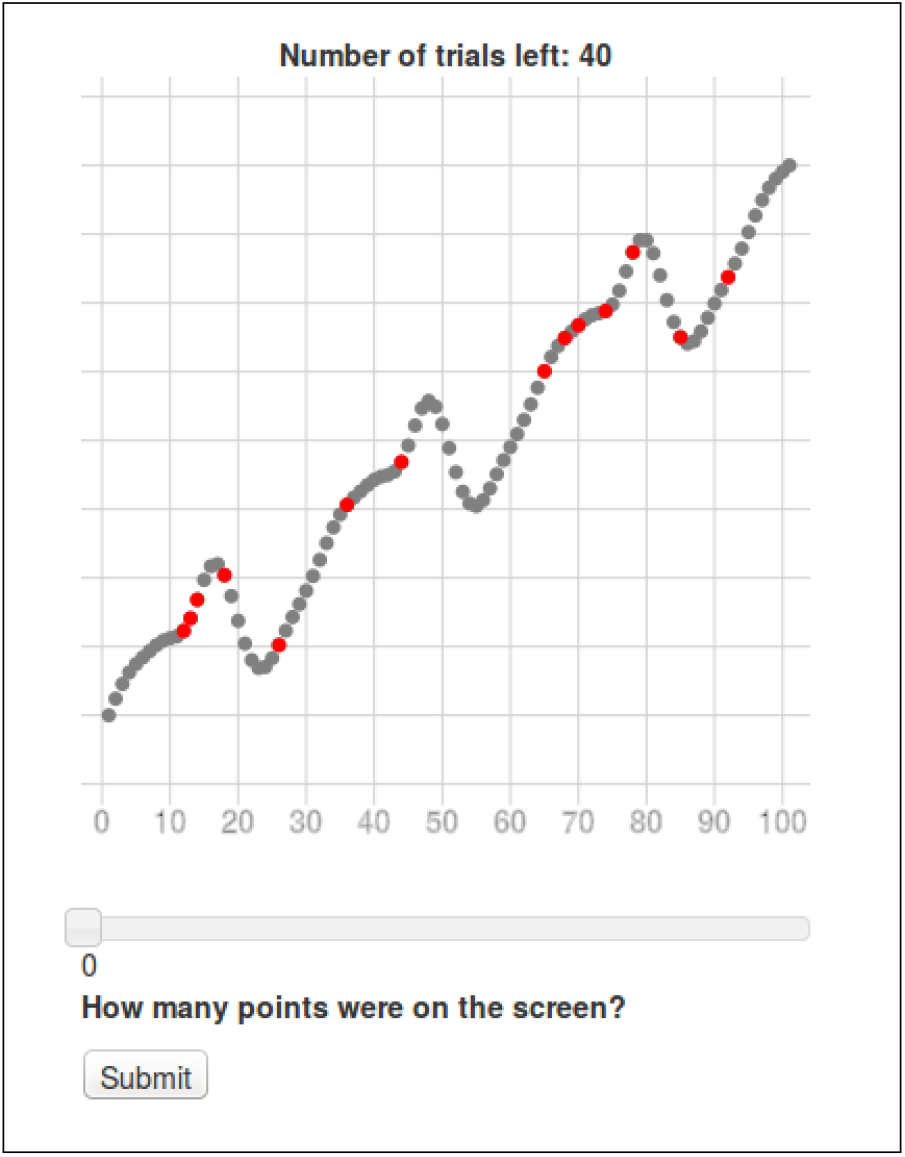
Screen shot of the numerosity experiment. Dots stayed on the screen for 1000ms and then vanished before the slider could be used.

### Results and discussion

Figure 22 shows the effect of the number of dots on participants’ numerosity judgments, averaged over both compositional and non-compositional functions, demonstrating that increasing the number of red dots led to greater underestimation of the actual number, consistent with the idea that perceived numerosity diminishes as structure becomes more discernible.

**Figure 22.**
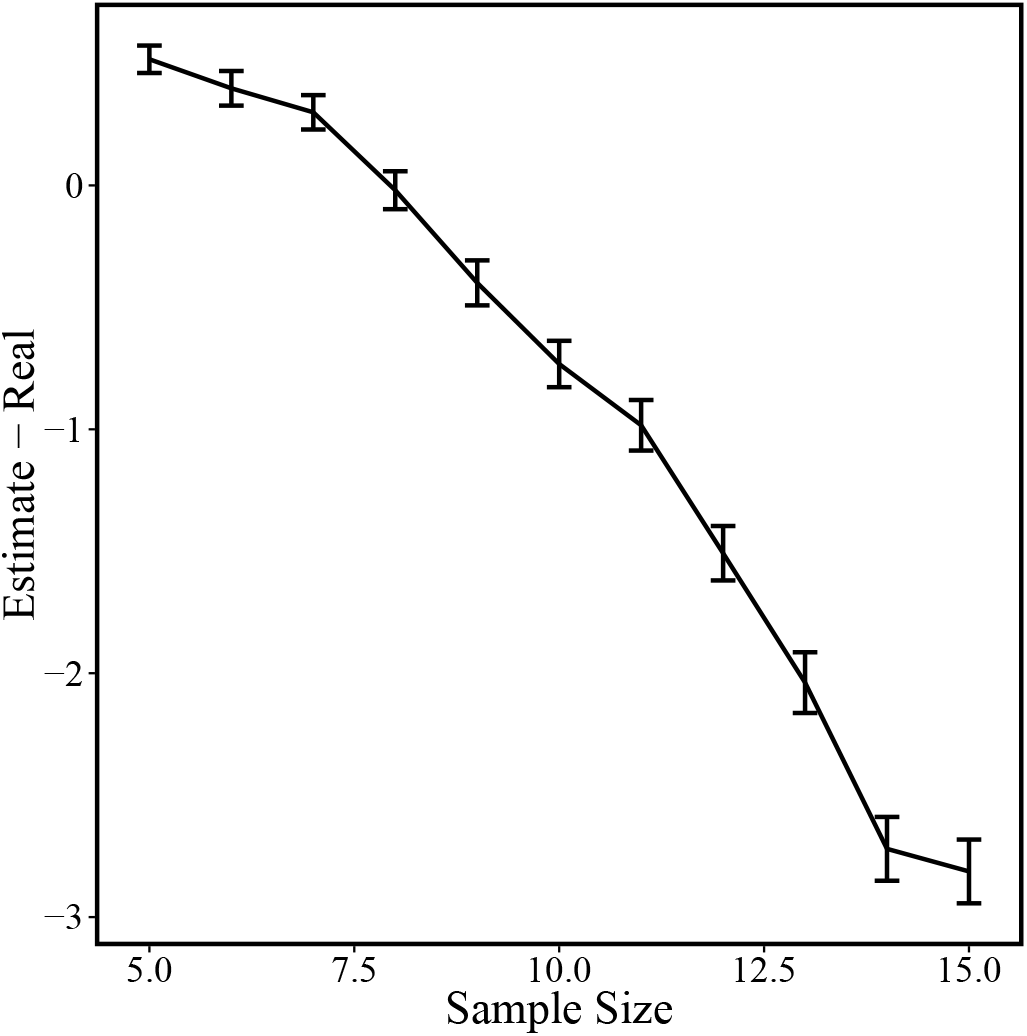
Mean difference between presented and estimated number of red dots over number of shown dots. Error bars represent the standard error of the mean. As more red dots are presented, participants tend to under-estimate the actual number more.

We hypothesized that participants would underestimate the red dots if they were superimposed on more structured functions (functions sampled from a compositional kernel) as compared to relatively unstructured functions (sampled from the spectral mixture kernel). Figure 23 shows the direct comparison of numerosity estimates for compositional and non-compositional patterns. Overall, participants underestimated the number of dots more for compositional than for non-compositional patterns(−0.74 vs. −0.53, hierarchical t-test: *t*(38) = −2.2, *p* < 0.05, *d* = 0.43), consistent with our hypothesis.

**Figure 23.**
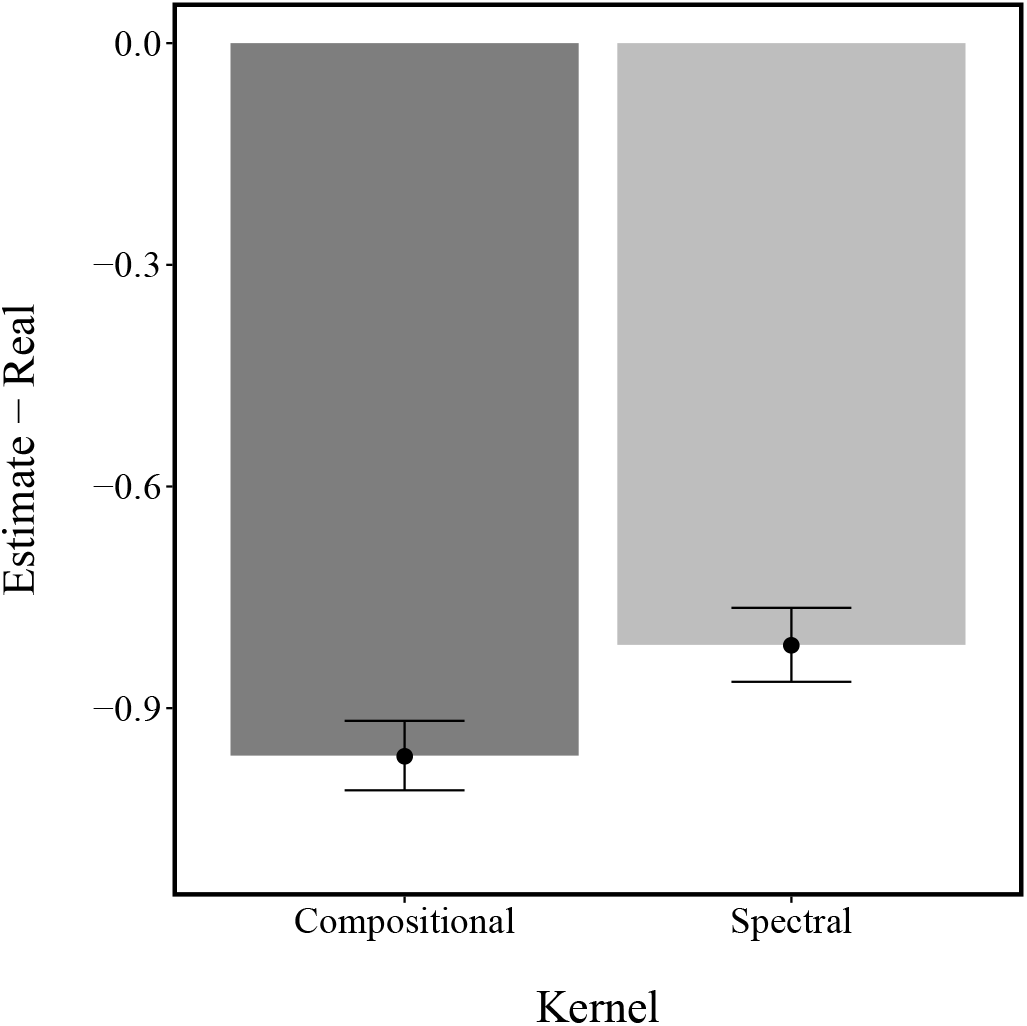
Comparative judgments. Lower values mean that participants underestimated the number of red dots. Error bars represent the standard error of the mean.

We performed a regression analysis to account for both the number of dots and the function type. The results (presented in Table 6) showed that with an increasing number of red dots, participants showed a stronger tendency to underestimate numerosity (*β* = −0.39). Additionally, we found a main effect of function type (*β* = −0.17). There was no interaction effect between the number of red dots and function type (*β* = −0.02).

**Table 6.** Parameter estimates from numerosity judgments regression analysis.

|  | Estimate | Std. Error | t-value | Pr(>\|t\|) |
| --- | --- | --- | --- | --- |
| Intercept | 3.23 | 0.12 | 27.631 | <2e-16 |
| Number of dots | -0.39 | 0.01 | -35.845 | <2e-16 |
| Compositional | -0.17 | 0.07 | -2.43 | 0.015 |
| Compositional × Number of dots | -0.02 | 0.02 | -0.76 | 0.446 |

Taking together, our results indicate that perceived numerosity is reduced to a greater extent by compositional functions compared to non-compositional functions, a finding that agrees with the results of Zhao and Yu (2016): structural regularity distorts the units of perception, making them appear less numerous. Even though the non-compositional functions we used were in fact structured (as measured by forecastability), they produced a weaker effect on numerosity relative to compositional functions, arguably because their structure is less “intuitive” (i.e., they lack the inductive bias needed to easily perceive this structure).

## Experiment 7: Change detection with functions

We next examined whether compositional structure influences change detection performance (Pashler, 1988; Rouder, Morey, Morey, & Cowan, 2011). In a typical change detection paradigm, participants judge whether two stimuli presented rapidly in sequence are the same or different. It has been suggested that structured representations facilitate change detection by allowing a summary representation of the stimulus to be stored in short-term memory (Brady & Tenenbaum, 2013). When the stimulus consists of multiple items, those items that are not assimilated into the summary representation are encoded as “outliers”. However, because memory is capacity-limited, only a small number of outliers can be encoded. Thus, the summary representation frees up encoding resources for specific items. The more structure in the display that can be encoded, the more resources will be available for encoding specific items.

The critical question concerns the nature of this structure—what are the inductive biases that constrain short-term memory representations? Brady and Tenenbaum (2013) used Markov random fields to encode information about object features; several other structural assumptions have been explored in the literature (Lew & Vul, 2015; Mathy & Feldman, 2012; Orhan & Jacobs, 2013). Here we explore the possibility that the representation of functions in short-term memory is compositional, leading to the prediction that change detection will be more accurate with compositional functions compared to with non-compositional functions. This prediction links the compressibility of functions directly to their generalizability. In fact, recent attempts in Bayesian machine learning have shown that models providing a good description of data can also be used for lossy compressions of the same data (see Ghahramani, 2015).

### Participants

66 participants (mean age=30, SD=8.2, 42 males) were recruited from Amazon Mechanical Turk and received $0.5 for their participation. The experiment took 10 minutes on average.

### Procedure

Participants judged whether or not two consecutively displayed patterns were the same (by pressing “J”) or different (by pressing “L”; see Figure 24). As in the numerosity experiment, we used functions from the matched set (20 compositional, 20 non-compositional, sampled without noise) and displayed them on the screen as 100 horizontally equidistant gray dots. On each trial, participants saw the original structure for 1000ms, followed by an interstimulus interval (500ms) and then a test probe (1000ms). Unbeknownst to participants, the probe had a 50% chance of being the same or different. Change probes were constructed by randomly selecting *N*∈{2,3,…,5} dots and permuting them (under the constraint that no point ends up at the same position as before).

**Figure 24.**
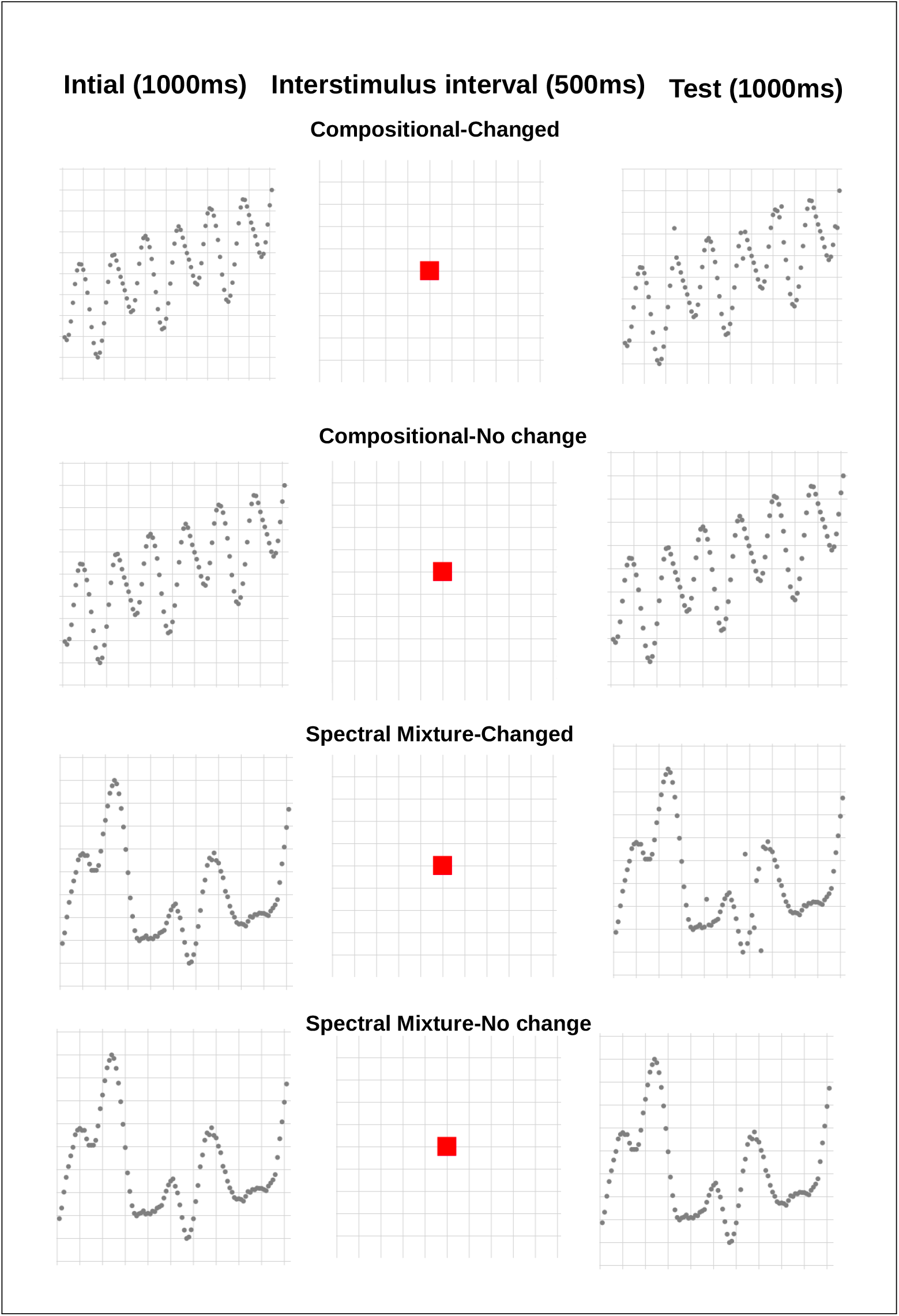
Schematic of change detection task. Initial stimulus was presented for 1000ms, drawn either from the compositional or from the non-compositional (spectral mixture) set. After an interstimulus interval (500ms), a test probe was presented and participants had to make a same/different judgment.

### Results and discussion

Participants responded correctly on 82% of no-change trials for the compositional functions and on 81% of the no-change trials for the non-compositional functions. Thus, there was no significant difference in correct rejection rate for the two types of functions (*χ*^2^(*N* = 1185, *df* = 1) = 0.21, *p* > 0.05). However, 77% of the changed compositional functions were correctly identified as having changed, whereas that proportion was only 67% for the changed non-compositional functions. Therefore, change is more easily detected for compositional functions (*χ*^2^(*N* = 1188, *df* = 1) = 12.13, *p* < 0.001). Moreover, 49 of 66 participants correctly identified change of compositional functions more frequently than change of non-compositional functions (*χ*^2^(*N* = 66, *df* = 1) = 9.67, *p* < 0.01).

Using a Bayesian hierarchical model to assess the posterior distribution of correctly identifying a probe revealed an estimate of 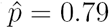 with a 95% credible set of [0.76, 0.81] for the compositional probes, whereas the same posterior was only 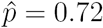 with a 95% credible set of [0.68, 0.75] for the non-compositional functions.

We created an indicator variable that was set to 1 if a participant responded correctly, and 0 otherwise. Figure 25 shows the mean proportions of correct responses for compositional and non-compositional trials across different levels of change: “no change”, “small change” (between 1 and 2 dots permuted) and “large change” (between 3 and 5 dots permuted). This analysis demonstrated that change detection performance was superior for compositional across levels of change.

**Figure 25.**
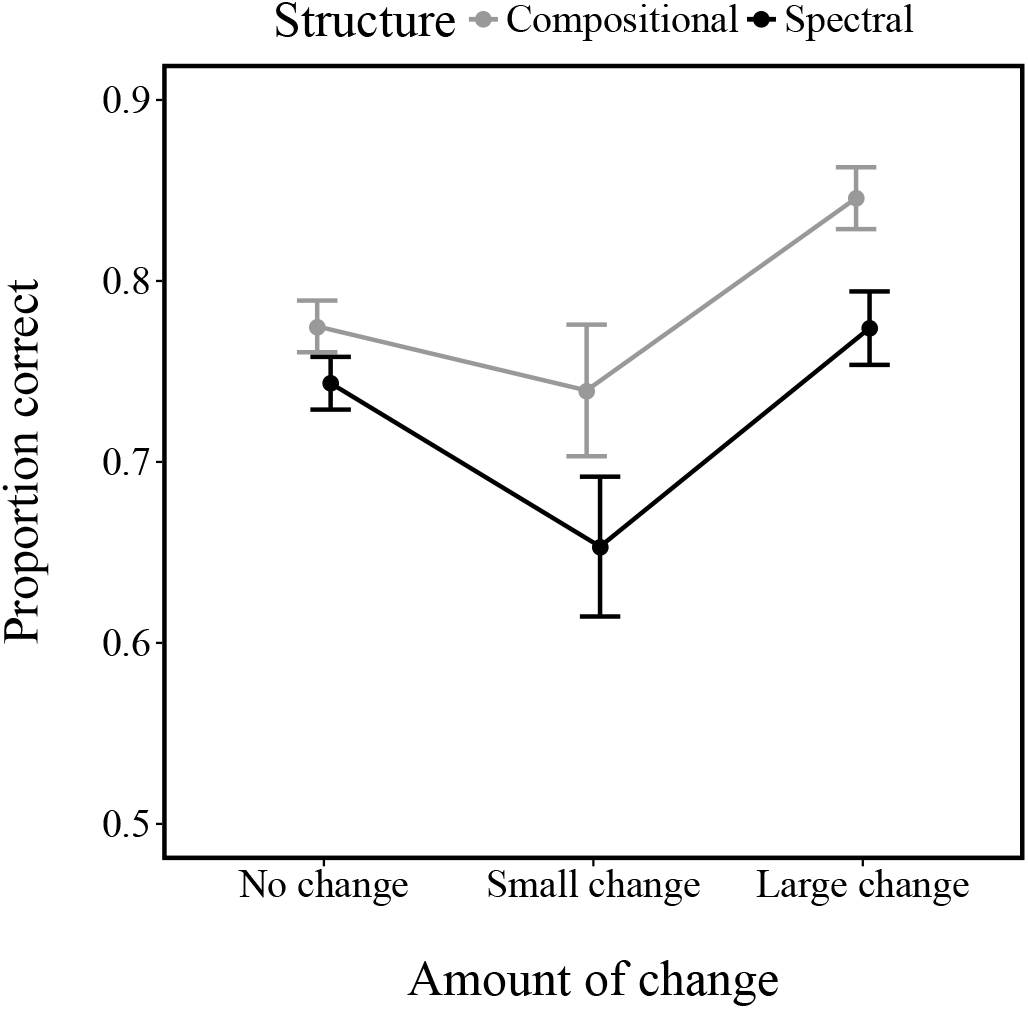
Proportion of correctly detected changes. Participants were more likely to detect changes in compositional functions. Error bars represent the standard error of the mean.

In order to quantitatively capture these results, we developed a Bayesian theory of functional change detection. Let 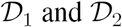 denote the first and second stimuli (input-output pairs), respectively, and let 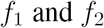 denote the corresponding functions. The task facing participants is to determine the posterior probability that the two functions are different:

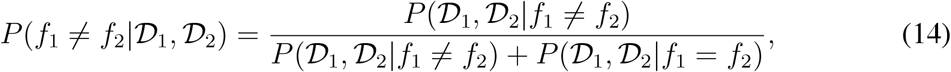

where for simplicity we have assumed that the prior probability of change is 0.5. The change 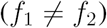 likelihood is given by:

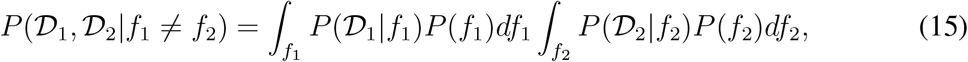

and the no-change 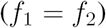 likelihood is given by:

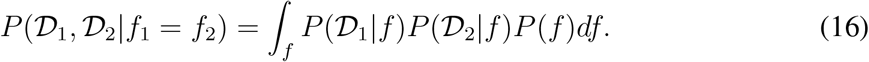

We used this probabilistic model to make trial-by-trial predictions of performance on the change detection task. As in our other analyses, we chose hyper-parameters that optimized the marginal likelihood for a given kernel. For the spectral kernel, we treated the number of mixture components as a free hyper-parameter (ranging from 1 to 6).

To allow for noise in the decision process, we modeled the binary responses using a logistic function of the model predictions, with separate predictors for compositional and non-compositional predictions. The results of this logistic regression are summarized in Table 7. The compositional model was a significant predictor of human change detection performance (*β* = 0.0122, *p* < 0.001), whereas the spectral mixture model was only marginally predictive (*p* = 0.05).

To show this result in a different way, we computed the point biserial correlation between the model predictions and human responses (Figure 26). The average correlation per participant for the compositional model was significantly higher than the average correlation for the spectral mixture model (*r* = 0.68 vs. *r* = 0.15, *t*(132) = 20.5, *p* < 0.001, *d* = 3.54).

**Table 7.** Result of change detection logistic regression.

|  | Estimate | Std. Error | z value | Pr(>\|z\|) |
| --- | --- | --- | --- | --- |
| Intercept | 1.3018 | 0.0918 | 14.19 | 0.0000 |
| Compositional prediction | 0.0122 | 0.0006 | 19.61 | 0.0000 |
| Spectral mixture prediction | 0.0012 | 0.0006 | 1.95 | 0.0517 |

**Figure 26.**
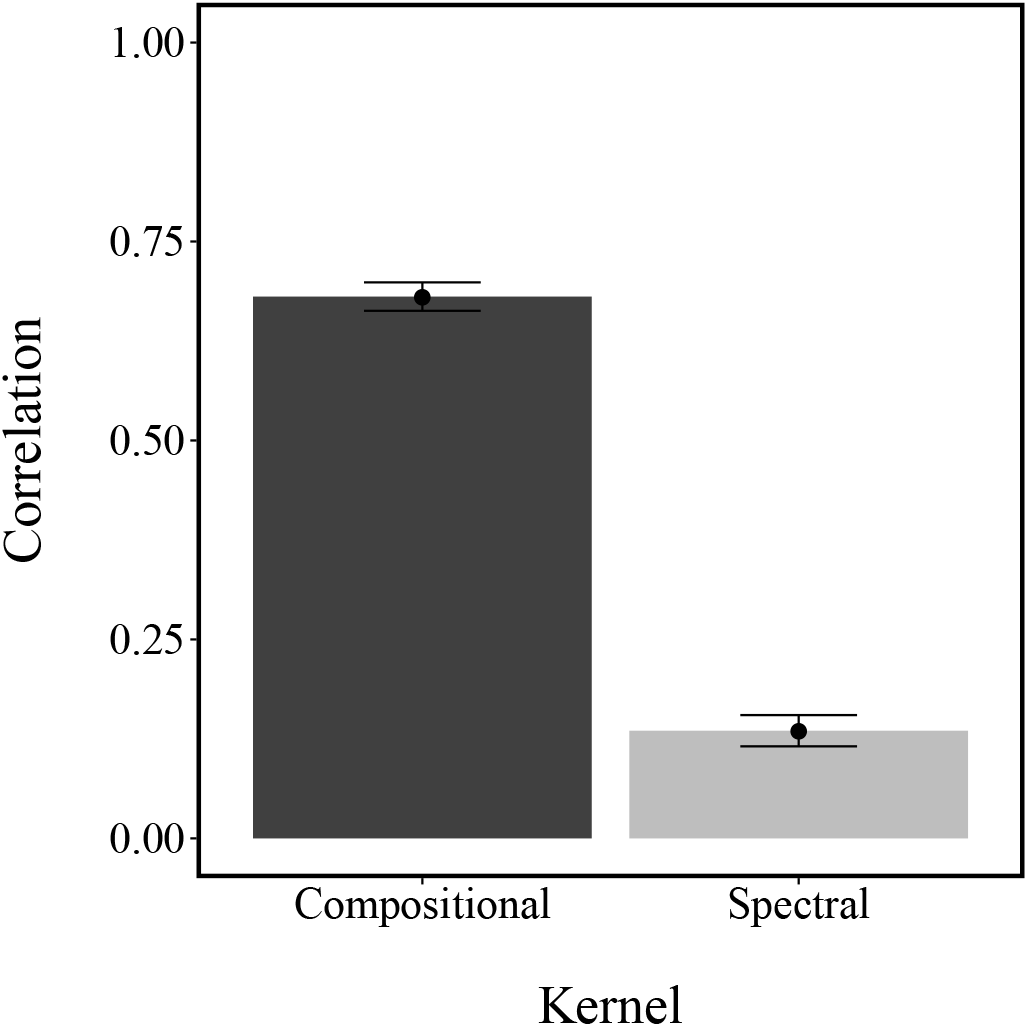
Mean point biserial correlation between model predictions and participants’ responses in the change detection task. Error bars represent the standard error of the mean.

In summary, we found that changes are more easily detected in compositional functions compared to non-compositional functions, consistent with the idea that compositional functions are more efficiently encoded into a summary representation. Moreover, a GP theory of change detection with a compositional kernel allowed us to quantitatively predict human change detection performance.

## Experiment 8: Compositional chunking in short-term memory

The results of Experiment 7 suggested that change detection is facilitated by compositional summary representations. A closely related idea is that structural regularities increase memory capacity because they are “compressible” (Brady, Konkle, & Alvarez, 2011; Mathy & Feldman, 2012). For example, if multiple items can be chunked together, then a greater number of items can be stored in memory (Miller, 1956). Chunking has been posited as the basis of exceptional expert memory (Chase & Simon, 1973; Gobet & Simon, 1998) and story comprehension (Thorndyke, 1977).

In our next experiment, we pursue this idea further, asking whether compositional functions are more compressible (and hence more memorable) than non-compositional functions. We used a standard short-term memory task (the Sternberg paradigm; Sternberg et al., 1966), in which participants are shown a rapid sequence of items (functions in this case) followed by an old/new judgment of a probe item. This task additionally allowed us to measure the interplay between compositionality and set size.

### Methods

#### Participants

133 participants (mean age=31.05, SD=8.19, 71 male) were recruited via Amazon Mechanical Turk and received $0.5 for their participation. The experiment took 9 minutes on average.

#### Procedure

Participants were shown between 2 and 6 functions sampled randomly from the matched set. Each function appeared on the screen for 1000ms. A 500ms intertrial interval succeeded the final item, followed by a probe item. Participants were asked to judge as quickly as possible whether the probe item was old (i.e., appeared in the preceding set) or new. There were 15 trials in total (3 trials for each set size). The probe was randomly selected to be either compositional or non-compositional, and probes were old on half of the trials. A schematic of the experiment is shown in Figure 27.

**Figure 27.**
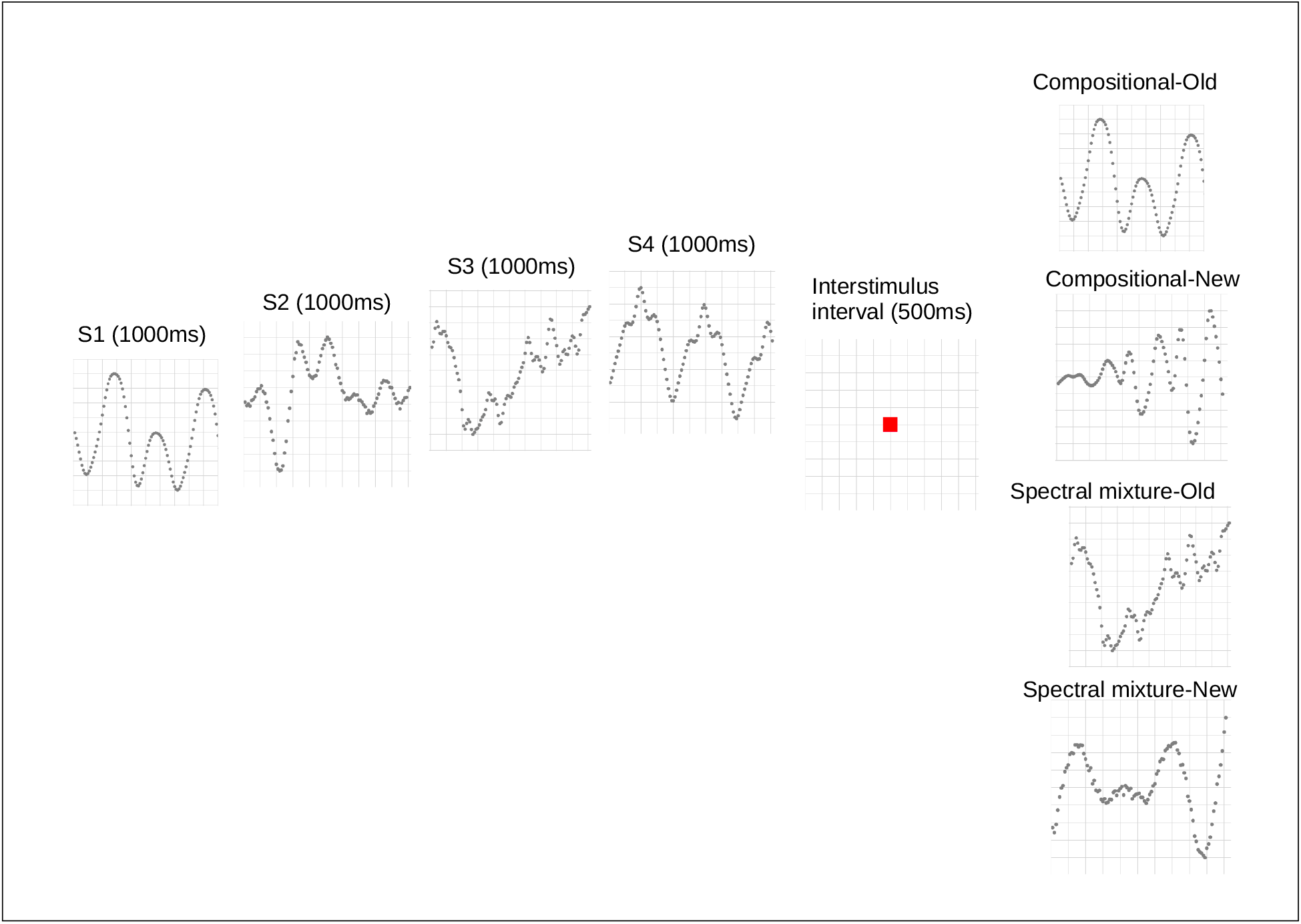
Schematic of the memory experiment. A sequence of stimuli was sampled from the matched set of functions. Every stimulus was presented for 1000ms, followed by an interstimulus interval (500ms), and then a probe (speeded old/new judgment).

### Results and discussion

We excluded 23 participants who failed to respond correctly for more than half of the trials, and removed all trials that took longer than 5 seconds or less than 500ms.

Participants responded correctly on 77.8% of the compositional probes and on 66.7% of the non-compositional probes—a significant difference in accuracy (*χ*^2^(*N* = 1456, *df* = 1) = 14.252, *p* < 0.001). 71 of 110 participants responded correctly more frequently to the compositional probes than to the non-compositional probes (*χ*^2^(*N* = 110, *df* = 1) = 9.3, *p* < 0.01).

This result can be decomposed further into a 75% hit rate for compositional probes, compared to a 67% hit rate for the non-compositional probes (a significant difference between hit rates: *χ*^2^(*N* = 709, *df* = 1) = 5.19, *p* < 0.05). 78 of 110 participants showed a higher hit rat for compositional probes as compared to non-compositional probes (*χ*^2^(*N* = 110, *df* = 1) = 19.2, *p* < 0.01).

Finally, there was also a significant difference between the correct rejection rate (78% for compositional probes vs. 65% for non-compositional probes;

*χ*^2^(*N* = 747, *df* = 1) = 8.87, *p* < 0.01). 76 of 110 participants showed a higher correct rejection rate for compositional probes as compared to non-compositional probes (*χ*^2^(*N* = 110, *df* = 1) = 17.92, *p* < 0.01).

Using a Bayesian hierarchical model to assess the posterior distribution of correctly identifying a probe, we found that this posterior was 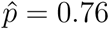 for the compositional probes with a 95% credible set of [0.74,0.78] and only 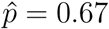 for the non-compositional probes with a 95% credible set [0.64,0.69].

To disentangle compositionality and set size, we ran a logistic regression to predict the probability of a correct response from compositionality (i.e., compositional vs. non-compositional probe) and set size factors. As shown in Table 8, the probability of responding correctly decreases with set size (*β* = −0.22), and compositional probes are more likely to be correctly identified (*β* = −0.37). We also found a significant interaction, whereby the compositional advantage decreases with set size (*β* = −0.18; Figure 28). This might be due to an increased guessing rate that could potentially obscure the compositional advantage.

**Table 8.** Results of logistic regression analysis of the memory experiment.

|  | Estimate | Std. Error | z-value | Pr(>\|z\|) |
| --- | --- | --- | --- | --- |
| Intercept | 2.04 | 0.29 | 7 | <2.57e-12 |
| Set size | -0.22 | 0.06 | -3.33 | <0.001 |
| Compositional | 1.24 | 0.37 | 3.33 | <0.001 |
| Set size × Compositional | - 0.18 | 0.08 | 2.2 | 0.02 |

**Figure 28.**
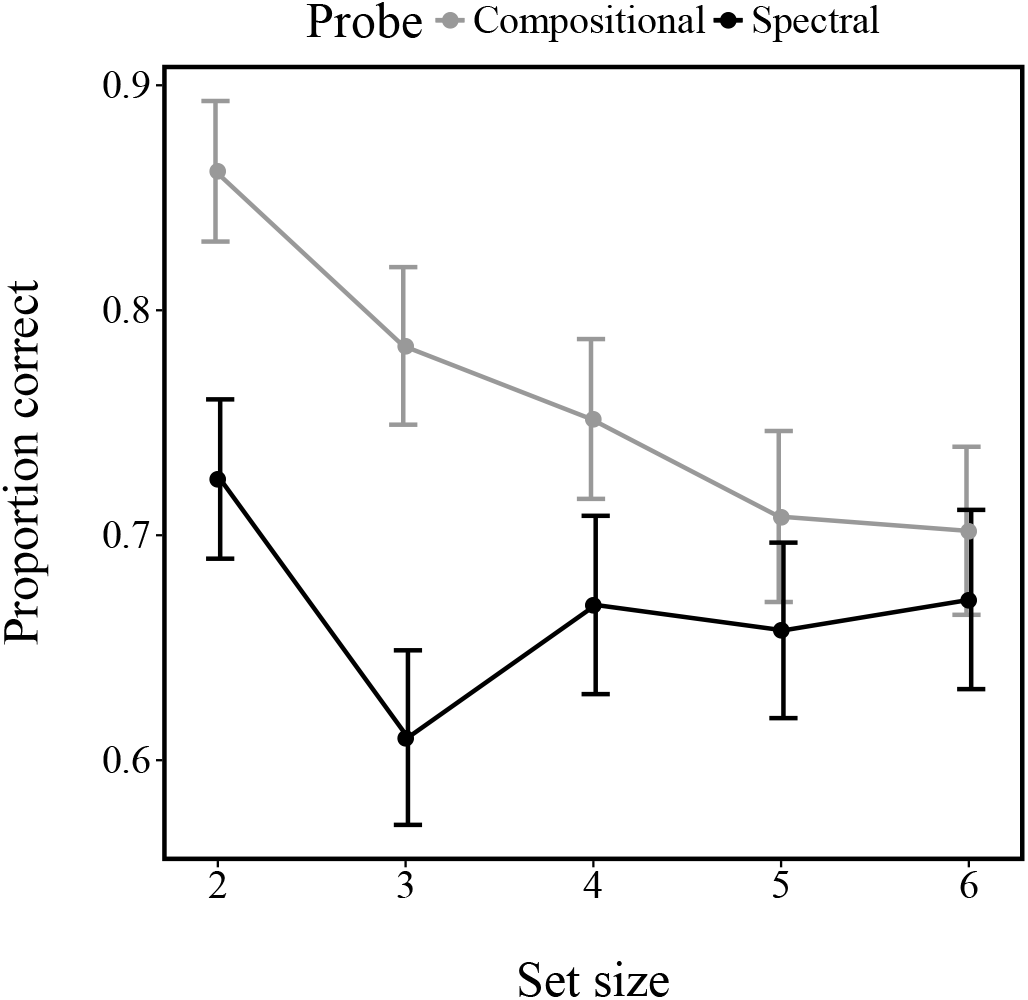
Proportion of correctly identified probes as a function of set size. Error bars represent the standard error of the mean.

We next developed a Bayesian model of performance in the task, adapting the same basic framework that we applied to the change detection task. A participant is exposed to a study list (sequence of independent input-output datasets), denoted by 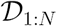, generated by underlying functions 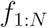. The task is to compute the posterior probability that a new dataset (the test probe), denoted by 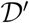, was generated by one of the functions in the study list:

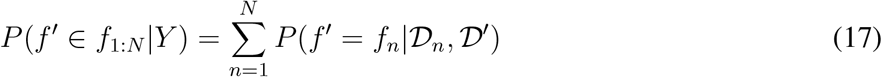

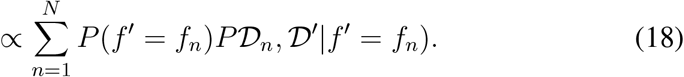

Following the structure of the experiment, we will assume that the probability of an “old” trial, 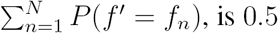. The marginal likelihood is given by:

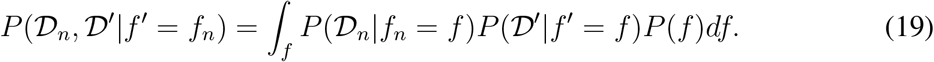

Our general optimization procedure is the same as described in the change detection model.

We entered the model predictions for both the compositional and spectral mixture models into a logistic regression to predict participant’s old/new responses. Results of the logistic regression analysis are summarized in Table 9. Only the compositional model was a significant predictor of responses (*β =* 0.054, *p* < 0.001), whereas the spectral mixture model was not a significant predictor (*β =* 0.01, *p =* 0.07).

**Table 9.**
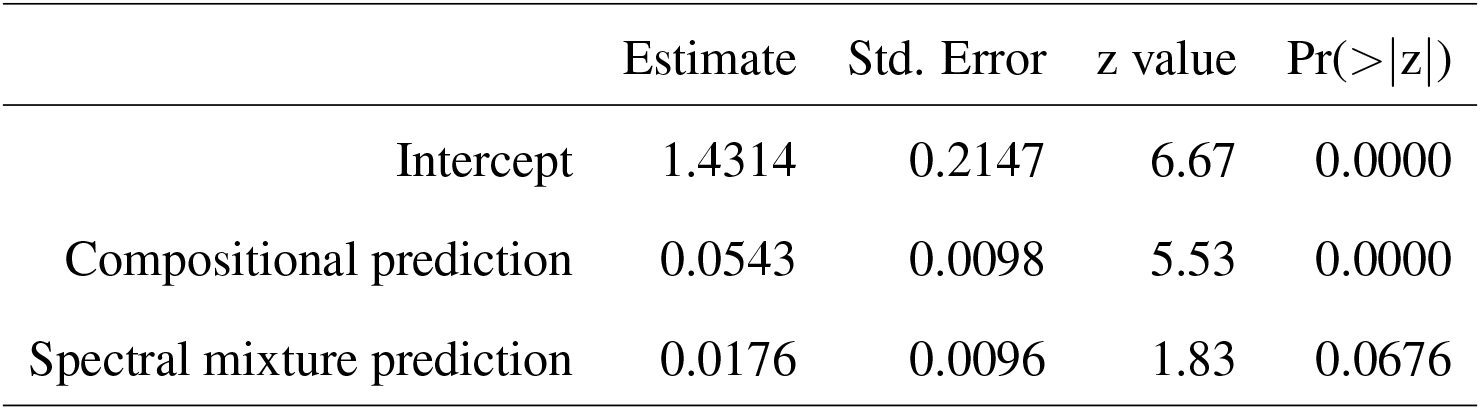
Results of logistic regression analysis of the memory experiment.

Finally, we calculated the correlation between each model’s predictions and participants’ responses. As shown in Figure 29, the compositional model produced a superior correlation compared to the non-compositional model (*r* = 0.35 vs. *r* = 0.09; *t*(262) = 9.97, *p* < 0.001, *d* = 0.34). Moreover, assessing two more simplistic models, we found that neither simply calculating a mean squared distance between the probe and all functions in the set (*r* = 0.094), nor counting the number of types within the set and then stating new if the probe is of the same type as most functions within the set and old otherwise (*r* = 0.04) led to better predictions of participants’ responses.

**Figure 29.**
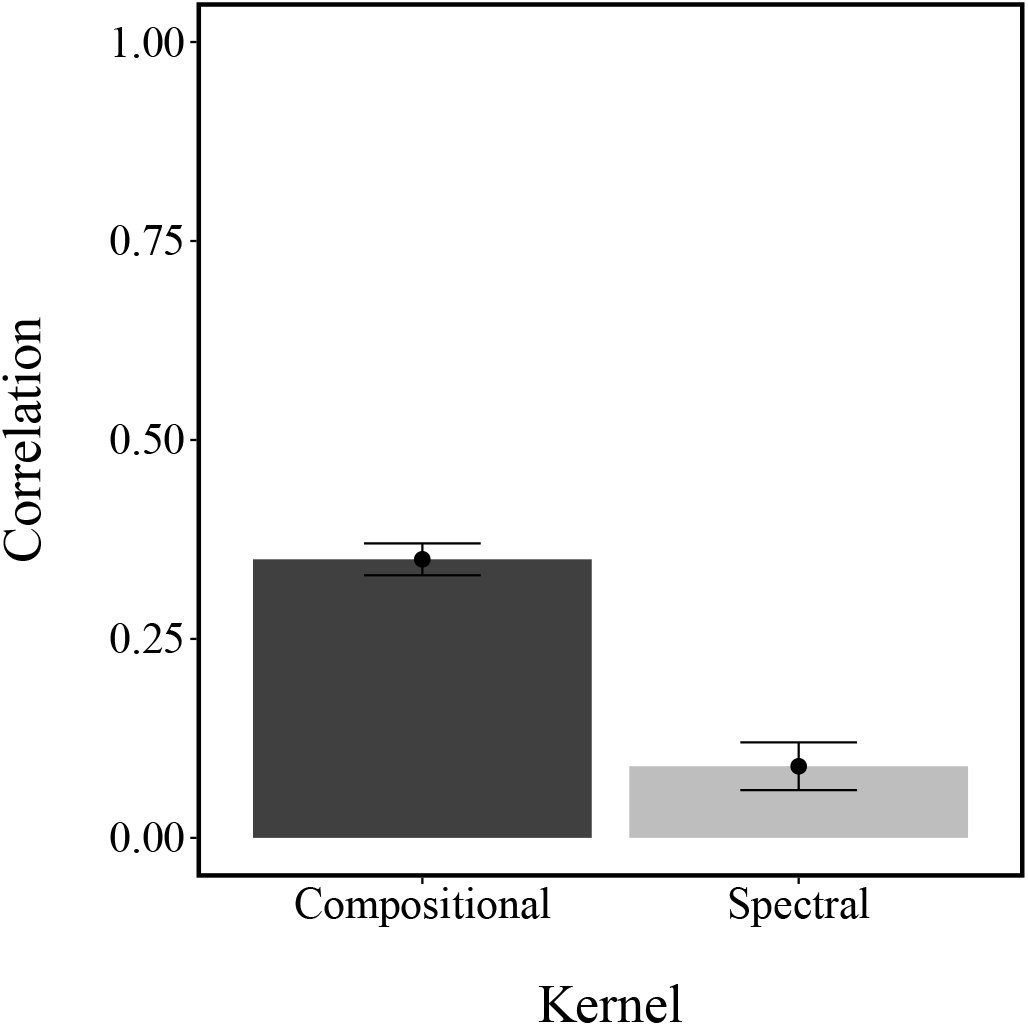
Mean point biserial correlation for the compositional and the spectral mixture models. Error bars represent the standard error of the mean.

We conclude that compositional functions are more easily remembered than non-compositional functions in a version of the Sternberg task. Participants’ old/new judgments were well-captured by a Bayesian model of short-term memory that used a compositional kernel.

## General discussion

In this paper, we pursued the hypothesis that human inductive biases for functional pattern recognition are compositionally structured—that is, humans prefer to represent complex functions as compositions of simpler building blocks. We formalized this idea using a compositional kernel within a Gaussian Process regression framework, and assessed human inductive biases across a diverse range of experiments. The first set (Experiments 1, 2, 3 and 5) attempted to directly measure inductive biases using extrapolation and interpolation judgments, finding that participants preferred compositional over non-compositional pattern completions. The second set of experiments examined the broader implications of compositionality, finding that compositional functions are perceived as more predictable (Experiment 4) and memorable (Experiment 8) compared to non-compositional functions. Furthermore, discrete displays of items are perceived as less numerous (a signature of statistical regularities; Experiment 6), and changes in such displays are more easily detected (Experiment 7). Taken together, our experimental findings provide strong support for the compositional hypothesis.

There are several reasons why intelligent agents might exhibit compositional inductive biases. First of all, if the world obeys a similar compositional structure, then compositional inductive biases support learning and generalization. This conjecture could be further assessed by testing if participants’ compositional priors over different domains track the structure within those domains, similar to what has been found in judgments about durations and other quantities (Griffiths & Tenenbaum, 2006). Moreover, compositionality can improve sample complexity and approximation accuracy under finite resource constraints (Mhaskar & Poggio, 2016). In principle, allowing for compositionality only ever costs a few extra bits of information (i.e. storing the compositional priors), but if structure exists that a grammar can express, then one can save an unbounded number of bits by detecting that structure.

Another reason for compositional inductive biases could be that compositionality helps memorizing structure by providing naturally occurring chunks. As seen in Experiment 8, participants memorized compositional structure more easily and a Bayesian model of compositional short-term memory for functions described their responses well. Thus, these biases might support the encoding and retrieval of structure in the real world. Finally, compositions might also make it easier to describe different structures and therefore support the social transmission of functions.

### Related work

The work presented here is connected to several lines of previous work. Most relevant are the seminal work of Griffiths et al. (2009) and Lucas et al. (2015) on Gaussian processes for function learning in general, and Wilson et al. (2015) more recent attempts to reverse-engineer the human kernel using a non-parametric kernel in particular. We see our work as complementary to this research. If we want to find out how people perceive and reason about functional structure then we need both a bottom-up theory to describe how people make sense of structure as well as a top-down method to indicate what the final structure might look like when represented as a kernel. Additionally, implementing a structure search algorithm as a parse tree as we have done here has recently been shown to be statistically efficient (Xu, Honorio, & Wang, 2017).

Our approach here also ties together neatly with past attempts to model compositional structure in other cognitive domains. Of course, language (e.g., Chomsky, 1965) and object perception (e.g., Biederman, 1987) have long traditions of emphasizing compositionality. More recently, these ideas have been extended to other domains. For example, Gershman, Tenenbaum, and Jäkel (2016) showed how hierarchical motion perception could be understood as a kind of vector analysis, using compositional GPs to model the combination of motion flow fields. This approach has also been applied to decision making; Gershman, Malmaud, and Tenenbaum (2016) used GPs to model utility functions over tree-structured objects (e.g., meals in a restaurant). In both motion perception and decision making, simpler non-compositional models failed to explain human performance.

### Limitations and future directions

While our experiments established the importance of compositional representations for functional pattern recognition, we do not believe that we have fully characterized the base functions and composition architecture. We restricted our framework to a small number of components primarily for practical purposes. Thus, an important direction for future work will be to systematically investigate the boundaries of function composition. Fortunately, the GP formalism can accommodate a wide variety of compositional structures (Duvenaud et al., 2013), so we expect that this task can be accomplished without deviating too far from the analytical framework laid out in this paper.

Another limitation is that we currently optimize the hyper-parameters of the kernels (for example, in the pattern completion experiments) a priori, whereas agents learning about functions in the real world naturally have to learn about both the underlying compositions as well as their hyper-parameters. One way to build a continuously updating model of compositional function learning could take inspiration from previously proposed models such as ALM (Busemeyer et al., 1997). ALM essentially implements a learning algorithm similar to a GP with an RBF kernel, but instead of using a Bayesian learning algorithm it is based on simple delta rule learning. Building a delta-learning based compositional function learning model and comparing it to other models of function learning provides a promising avenue for future research.

In this work we probed in what sense (if any) human functional inductive biases can be seen as compositional within a single prediction task. However, composing structures learned across different tasks could provide even more powerful learning strategies. Future work could investigate the composition of structure across tasks (as for example shown by Hwang, Tong, & Choi, 2016), for example in one-shot learning experiments.

There are many other potential implications of compositional inductive biases for functions. For example, these inductive biases could shape active learning (i.e., tasks in which participants can choose the next data point). We believe that active learning model comparisons constitute an interesting framework to pit the structured and unstructured approaches against each other (Parpart, Schulz, Speekenbrink, & Love, 2015).

Related issues arise in reinforcement learning tasks, where agents must balance exploration and exploitation. Such tasks have already been modeled using GPs (Schulz, Konstantinidis, & Speekenbrink, 2016; Wu, Schulz, Speekenbrink, Nelson, & Meder, 2017), suggesting their amenability to analysis with compositional kernels. This approach could also be applied to designing nonparametric value function approximators, which have been proposed as cognitively and neurally plausible solutions to reinforcement learning problems (Gershman & Daw, 2015). New experiments will be required to discern whether reinforcement learning exploits compositional inductive biases.

Finally, the development of compositional inductive biases for functions is another important and open question. Very little is known in general about the development of function learning. The studies described in this paper could potentially be run in children, which would provide insight into the origin of compositional inductive biases.

### Conclusions

We proposed that people exhibit compositional inductive biases in function learning and assessed a compositional model of function learning across 8 experimental paradigms. Our model not only provides a good account of participants’ completion, learning, perception, and memorization of functional structure, but also resonates well with a general trend in the cognitive sciences focusing on the importance of compositionality for intelligent behavior (Fodor & Pylyshyn, 1988; Lake et al., 2015; Piantadosi et al., 2016; Tenenbaum, Kemp, Griffiths, & Goodman, 2011). Taken together, our results show how people manage to construct richly structured representations from simple functional building blocks.

## Appendix A Gaussian Process Inference

In the case that there is only one test point 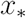, the posterior mean 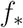, can be calculated as:

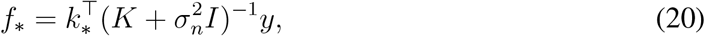

where 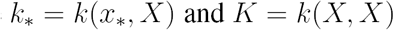. The posterior variance of the prediction is given by:

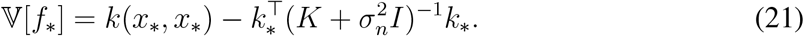

The marginal likelihood is given by integrating over the latent function *f*:

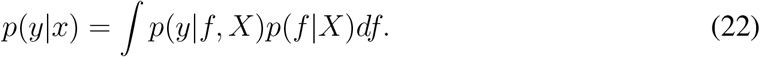

The log marginal likelihood for a GP with hyper-parameters *θ* is given by:

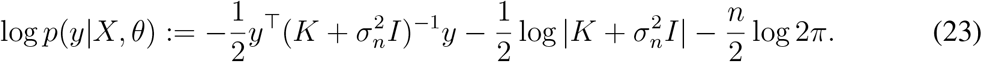

where the dependence of *K* on *θ* is left implicit. The hyper-parameters are chosen to maximize the log-marginal likelihood, using gradient-based optimization.

## Appendix B Spectral mixture kernel

**Bochner’s theorem**

A stationary kernel is a function of *τ* = *x — x′.* A complex-valued function 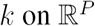 is the covariance function of a weakly stationary mean-square continuous complex-valued random process on 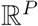 if and only if it can be represented as:

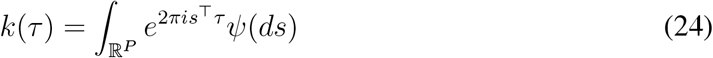

where *ψ* is a positive finite measure. The density *S(s)* of *ψ* is called the spectral density of *k; k* and *S* are Fourier duals:

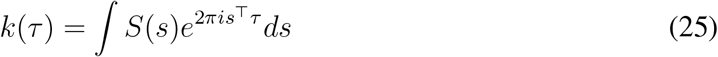

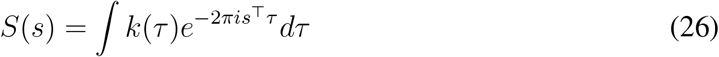

Therefore, every kernel can also be represented by a distribution over the spectral density space.

**Mixture of Gaussians spectral kernel**

Our treatment of the spectral kernel follows Wilson et al. (2015), and we refer the reader to that paper for more details. The spectral density modeled with a single Gaussian can be expressed as:

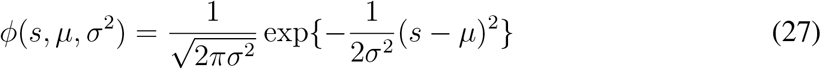

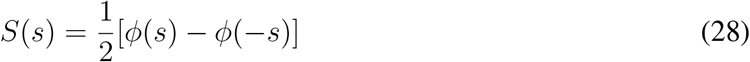

The resulting kernel is given by:

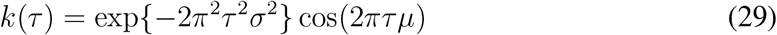

Extending this result to a mixture of *Q* Gaussians results in:

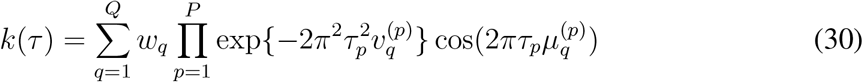

where the *q*th component has the mean 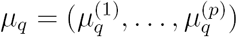 and covariance matrix 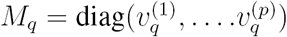, where the inverse mean represents the component periods and the inverse standard deviation the length scales.

## Appendix C Hierarchical model

The Bayesian model used to assess the posterior probabilities in Experiments 1, 7, and 8 is based on a hierarchical binomial model with the hyper-prior *κ* ~ Pareto(1, 1.5), prior 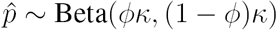; and the likelihood 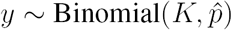. Inference was performed using MCMC as implemented in Stan (Stan Development Team, 2016).

## Appendix D Assessing the Bezier smoothing technique

We ran simulations to assess whether the Bezier smoothing technique applied in Experiment 3 forced participants’ drawings to be smoother than intended. We sampled 1000 functions (100 points for each) from an RBF kernel and 1000 functions from a less smooth Ornstein-Uhlenbeck kernel (cf. Schulz et al., 2015), added normally distributed noise 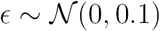 to every observation, and submitted these functions to the Bezier smoothing algorithm. Afterwards, we used the extracted (Bezier-smoothed) function and performed the same model comparison as in Experiment 3, assessing which of the two kernels led to better out-of-sample predictions given different ground truths of the underlying smoothness. Whereas the RBF kernel was always favored when the ground truth was sampled from a RBF kernel (mean squared prediction error: 4.94 vs. 4.52), the Ornstein-Uhlenbeck kernel was preferred for 505 of 1000 samples and led to a lower prediction error if the ground truth was sampled from an Ornstein-Uhlenbeck kernel (mean squared prediction error: 3.72 vs. 3.74). Thus, even though the Bezier smoothing technique has a tendency to prefer smoother functions, it is nonetheless able to pick up on the intention to draw less smooth functions.

## Appendix E Details on generation of comparison functions

**Forecastibility**

The Shannon entropy of a function *f* can be measured by the uncertainty of its spectrum *S*_*f*_:

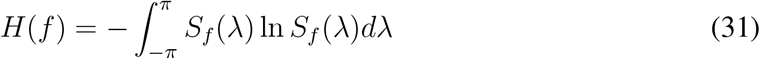

For any stationary function,

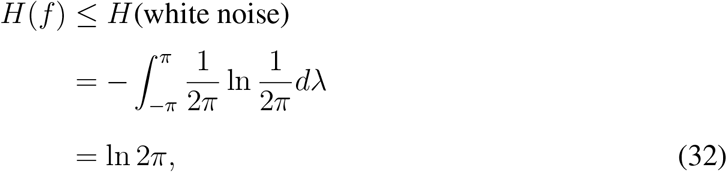

with equality if and only if *f* is white noise, the least forecastable signal with a uniform spectrum. The forecastibility measure Ω(*f*) is then defined as

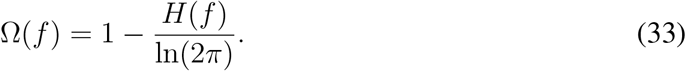

We follow Goerg (2013) and estimate Ω(*f*) by first estimating the spectrum *S*_*f*_, normalizing it so that it integrates to 1, and then plugging it back into Eq. 31. We use the periodogram as an unbiased estimator of *S*_*f*_ (see Goerg, 2013, for details).

**Wavelet transform similarity measure**

A discrete wavelet Haar Transform performs a scale-wise decomposition of the time series in such a way that most of the energy of the time series can be represented by a few coefficients. The main idea is to replace the original series by its wavelet approximation coefficients **a** in an appropriate scale, and then to measure the dissimilarity between the wavelet approximations. A detailed description of wavelet methods for time series analysis can be found in Percival and Walden (2006). We used the R-package `TSclust` (Montero et al., 2014) to find the appropriate scale of the transform. We then measured the dissimilarity between two series *x*_1_ and *x*_2_ by the Euclidean distance at the selected scale: *d*(*x*_1_, *x*_2_) = ||a_1_ — a_2_||.

**Is the Haar transform an intuitive measure of similarity?**

To assess whether the wavelet Haar transform is an intuitive measure of functional similarity, we sampled 50 functions from a radial basis function kernel and 50 functions from an Ornstein-Uhlenbeck kernel and asked 50 MTurkers (mean age=32.94, SD=14.04, 19 female, flat payment of $ 0.3) to rate the similarity between pairs of randomly selected functions from this pool for 20 consecutive trials on a scale ranging from 0 (not at all similar) to 100 (very similar). The results of this experiment revealed that the Wavelet transform similarity measure was predictive of participants’ judgments with an average correlation between similarity judgments and the wavelet Haar distance between the two functions of *r* = −0.49, *p* < 0.001 over all comparisons, *r* = −0.60, *p* < 0.001 when only comparing functions sampled from the radial basis function kernel, and *r* = −0.62, *p* < 0.001, when only comparing functions from the Ornstein-Uhlenbeck kernel.

## Footnotes

1 Even though the ability to extrapolate more complex functions such as periodic patterns has been debated (see Kalish, 2013), and participants seem to have more difficulty learning cyclic instead of quadratic functions (Byun, 1995), they generally seem to be able to learn non-linear functions (Busemeyer et al., 1997), especially if the representation format supports encoding (DeLosh, 1995).

2 In general *χ* can be multidimensional and discrete, but in this paper we focus on functions with one-dimensional, continuous inputs since these have received the most attention in the function learning literature (but see Juslin, Olsson, & Olsson, 2003; Lewandowsky, Kalish, & Ngang, 2002).

3 A mathematically precise characterization of smoothness induced by a given kernel function is given by the theory of reproducing kernel Hilbert spaces (Schölkopf & Smola, 2002).

4 Generating functions with a positive linear trend can be achieved by using the linear kernel shown in Table 1 and combining it with a non-zero mean function that is increasing.

5 A mixture of linear and radial basis function extrapolations can lead to longer-distance dependencies between points than a radial basis function alone.

6 This is also true for approaches utilizing a mixture of expert kernel, which is fixed *a priori*, but see Kalish (2013) for further discussion.

7 A stationary kernel is a function of x – x′. Thus, it is invariant to translation of the inputs.

8 This is essentially a form of extrapolation judgment, but unlike typical extrapolation paradigms that test input-output pairs one at a time, pattern completion asks participants to consider a set of input-output pairs.

9 Most of these functions were generated from two composed kernels and contained either a linear or a periodic component.

10 As there is currently no reliable way to generate an exact posterior over compositional parts, the normalized marginal likelihood of each composition was used as a rough approximation.

11 See the following page for an example: http://learning.eng.cam.ac.uk/carl/mauna.

12 Most participants found this set-up to be very intuitive. The source code for this experiment can be found online at https://github.com/ericschulz/drawfunctions. See also Appendix D for further analysis on the effect of smoothing the input points.

13 Predictability is closely related to forecastability, insofar as both measures can be characterized in terms of the eigenspectrum (Sollich, 2001). Here we focus on predictability since it has a more intuitive interpretation.

14 We avoided explicitly mentioning the term “function” in this experiment in order to discourage participants from imposing familiar functional forms on the data.

## References

Andreassen, P. B., & Kraus, S. J. (1990). Judgmental extrapolation and the salience of change. Journal of forecasting, 9, 347–372.

Berry, D. C., & Broadbent, D. E. (1984). On the relationship between task performance and associated verbalizable knowledge. The Quarterly Journal of Experimental Psychology, 36, 209–231.

Biederman, I. (1987). Recognition-by-components: a theory of human image understanding. Psychological Review, 94, 115–147.

Bolger, F., & Harvey, N. (1998). Heuristics and biases in judgmental forecasting.

Bott, L., & Heit, E. (2004). Nonmonotonic extrapolation in function learning. Journal of Experimental Psychology: Learning, Memory, and Cognition, 30, 38–50.

Brady, T. F., Konkle, T., & Alvarez, G. A. (2011). A review of visual memory capacity: Beyond individual items and toward structured representations. Journal of Vision, 11, 4–4.

Brady, T. F., & Tenenbaum, J. B. (2013). A probabilistic model of visual working memory: Incorporating higher order regularities into working memory capacity estimates. Psychological Review, 120, 85.

Brehmer, B. (1974a). The effect of cue intercorrelation on interpersonal learning of probabilistic inference tasks. Organizational Behavior and Human Performance, 12, 397–412.

Brehmer, B. (1974b). Hypotheses about relations between scaled variables in the learning of probabilistic inference tasks. Organizational Behavior and Human Performance, 11, 1–27.

Brehmer, B., Alm, H., & Warg, L.-E. (1985). Learning and hypothesis testing in probabilistic inference tasks. Scandinavian Journal of Psychology, 26, 305–313.

Broadbent, D. E. (1958). The effects of noise on behaviour.

Bruner, J. S., Goodnow, J. J., & George, A. (1956). A study of thinking. New York: John Wiley & Sons, Inc, 14, 330.

Buffart, H., Leeuwenberg, E., & Restle, F. (1981). Coding theory of visual pattern completion. Journal of Experimental Psychology: Human Perception and Performance, 7, 241.

Busemeyer, J. R., Byun, E., Delosh, E. L., & McDaniel, M. A. (1997). Learning functional relations based on experience with input-output pairs by humans and artificial neural networks.

Byun, E. (1995). Interaction between prior knowledge and type of nonlinear relationship on function learning.

Carroll, J. D. (1963). Functional learning: The learning of continuous functional mappings relating stimulus and response continua. Educational Testing Service.

Chase, W. G., & Simon, H. A. (1973). Perception in chess. Cognitive Psychology, 4, 55–81.

Chomsky, N. (1965). Aspects of the theory of syntax. MIT press.

Cox, G. E., Kachergis, G., & Shiffrin, R. M. (2012). Gaussian process regression for trajectory analysis. In Proceedings of the 34th Annual Conference of the Cognitive Science Society (pp. 1440–1445).

DeLosh, E. L. (1995). Hypothesis testing in the learning of functional concepts. Unpublished master’s thesis, Purdue University, West Lafayette, IN.

DeLosh, E. L., Busemeyer, J. R., & McDaniel, M. A. (1997). Extrapolation: The sine qua non for abstraction in function learning. Journal of Experimental Psychology: Learning, Memory, and Cognition, 23, 968.

Duvenaud, D., Lloyd, J. R., Grosse, R., Tenenbaum, J. B., & Ghahramani, Z. (2013). Structure discovery in nonparametric regression through compositional kernel search. Proceedings of the 30th International Conference on Machine Learning, 1166–1174.

Eggleton, I. R. (1982). Intuitive time-series extrapolation. Journal of Accounting Research, 68–102.

Field, D. J., Hayes, A., & Hess, R. F. (1993). Contour integration by the human visual system: evidence for a local âĂIJassociation fieldâĂİ. Vision research, 33, 173–193.

Flanagan, J. R., Nakano, E., Imamizu, H., Osu, R., Yoshioka, T., & Kawato, M. (1999). Composition and decomposition of internal models in motor learning under altered kinematic and dynamic environments. Journal of Neuroscience, 19, RC34–RC34.

Fodor, J. A. (1975). The language of thought (Vol. 5). Harvard University Press.

Fodor, J. A., & Pylyshyn, Z. W. (1988). Connectionism and cognitive architecture: A critical analysis. Cognition, 28, 3–71.

Forrest, A. R. (1972). Interactive interpolation and approximation by Bézier polynomials. The Computer Journal, 15, 71–79.

Gershman, S. J., & Daw, N. D. (2015). Reinforcement learning and episodic memory in humans and animals: An integrative framework. Annual Review of Psychology, 68.

Gershman, S. J., Malmaud, J., & Tenenbaum, J. B. (2016). Structured representations of utility in combinatorial domains. Decision.

Gershman, S. J., & Niv, Y. (2010). Learning latent structure: carving nature at its joints. Current Opinion in Neurobiology, 20, 251–256.

Gershman, S. J., Tenenbaum, J. B., & Jäkel, F. (2016). Discovering hierarchical motion structure. Vision Research.

Ghahramani, Z. (2015). Probabilistic machine learning and artificial intelligence. Nature, 521, 452.

Ghahramani, Z., & Wolpert, D. M. (1997). Modular decomposition in visuomotor learning. Nature, 386, 392.

Gobet, F., & Simon, H. A. (1998). Expert chess memory: Revisiting the chunking hypothesis. Memory, 6, 225–255.

Goerg, G. (2013). Forecastable component analysis. In Proceedings of the 30th International Conference on Machine Learning (ICML-13) (pp. 64–72).

Goodman, N. D., Tenenbaum, J. B., Feldman, J., & Griffiths, T. L. (2008). A rational analysis of rule-based concept learning. Cognitive Science, 32, 108–154.

Goodwin, P., & Wright, G. (1993). Improving judgmental time series forecasting: A review of the guidance provided by research. International Journal of Forecasting, 9, 147–161.

Griffiths, T. L., Chater, N., Kemp, C., Perfors, A., & Tenenbaum, J. B. (2010). Probabilistic models of cognition: Exploring representations and inductive biases. Trends in Cognitive Sciences, 14, 357–364.

Griffiths, T. L., & Kalish, M. L. (2007). Language evolution by iterated learning with Bayesian agents. Cognitive Science, 31, 441–480.

Griffiths, T. L., Lucas, C., Williams, J., & Kalish, M. L. (2009). Modeling human function learning with gaussian processes. In Advances in Neural Information Processing Systems (pp. 553–560).

Griffiths, T. L., & Tenenbaum, J. B. (2006). Optimal predictions in everyday cognition. Psychological science, 17, 767–773.

Harvey, N., & Bolger, F. (1996). Graphs versus tables: Effects of data presentation format on judgemental forecasting. International Journal of Forecasting, 12, 119–137.

Harvey, N., Ewart, T., & West, R. (1997). Effects of data noise on statistical judgement. Thinking & Reasoning, 3, 111–132.

Hwang, Y., Tong, A., & Choi, J. (2016). Automatic construction of nonparametric relational regression models for multiple time series. In Proceedings of the 33rd international conference on machine learning.

Hyndman, R. J., Wang, E., & Laptev, N. (2015). Large-scale unusual time series detection. In 2015 IEEE International Conference on Data Mining Workshop (ICDMW) (pp. 1616–1619).

Juslin, P., Olsson, H., & Olsson, A.-C. (2003). Exemplar effects in categorization and multiple-cue judgment. Journal of Experimental Psychology: General, 132, 133.

Kalish, M. L. (2013). Learning and extrapolating a periodic function. Memory & cognition, 41, 886–896.

Kalish, M. L., Griffiths, T. L., & Lewandowsky, S. (2007). Iterated learning: Intergenerational knowledge transmission reveals inductive biases. Psychonomic Bulletin & Review, 14, 288–294.

Kalish, M. L., Lewandowsky, S., & Kruschke, J. K. (2004). Population of linear experts: knowledge partitioning and function learning. Psychological Review, 111, 1072.

Kanizsa, G. (1979). Organization in vision: Essays on gestalt perception. Praeger Publishers.

Kemp, C. (2012). Exploring the conceptual universe. Psychological Review, 119, 685.

Kemp, C., & Tenenbaum, J. B. (2009). Structured statistical models of inductive reasoning. Psychological Review, 116, 20–58.

Keren, G. (1983). Cultural differences in the misperception of exponential growth. Perception & psychophysics, 34, 289–293.

Koh, K., & Meyer, D. E. (1991). Function learning: Induction of continuous stimulus-response relations. Journal of Experimental Psychology: Learning, Memory, and Cognition, 17, 811–836.

Kwantes, P. J., & Neal, A. (2006). Why people underestimate y when extrapolating in linear functions. Journal of Experimental Psychology: Learning, Memory, and Cognition, 32, 1019.

Lake, B. M., Salakhutdinov, R., & Tenenbaum, J. B. (2015). Human-level concept learning through probabilistic program induction. Science, 350, 1332–1338.

Lake, B. M., Ullman, T. D., Tenenbaum, J. B., & Gershman, S. J. (2016). Building machines that learn and think like people. Behavioral and Brain Sciences.

Lee, T. S., & Yuille, A. L. (2006). Efficient coding of visual scenes by grouping and segmentation. Bayesian Brain: Probabilistic Approaches to Neural Coding, 141–185.

Lew, T. F., & Vul, E. (2015). Ensemble clustering in visual working memory biases location memories and reduces the weber noise of relative positions. Journal of Vision, 15, 10–10.

Lewandowsky, S., Kalish, M., & Ngang, S. (2002). Simplified learning in complex situations: Knowledge partitioning in function learning. Journal of Experimental Psychology: General, 131, 163.

Little, D. R., & Shiffrin, R. (2009). Simplicity bias in the estimation of causal functions. In Proceedings of the cognitive science society (Vol. 31).

Lucas, C. G., Griffiths, T. L., Williams, J. J., & Kalish, M. L. (2015). A rational model of function learning. Psychonomic Bulletin & Review, 22, 1193–1215.

Mathy, F., & Feldman, J. (2012). What’s magic about magic numbers? Chunking and data compression in short-term memory. Cognition, 122, 346–362.

McDaniel, M. A., & Busemeyer, J. R. (2005). The conceptual basis of function learning and extrapolation: Comparison of rule-based and associative-based models. Psychonomic Bulletin & Review, 12, 24–42.

McDaniel, M. A., Dimperio, E., Griego, J. A., & Busemeyer, J. R. (2009). Predicting transfer performance: A comparison of competing function learning models. Journal of Experimental Psychology: Learning, Memory, and Cognition, 35, 173.

Mhaskar, H. N., & Poggio, T. (2016). Deep vs. shallow networks: An approximation theory perspective. Analysis and Applications, 14, 829–848.

Miller, G. A. (1956). The magical number seven, plus or minus two: Some limits on our capacity for processing information. Psychological Review, 63, 81–97.

Mitchell, T. M. (1980). The need for biases in learning generalizations. Department of Computer Science, Laboratory for Computer Science Research, Rutgers Univ. New Jersey.

Montero, P., Vilar, J. A., et al. (2014). Tsclust: An r package for time series clustering. Journal of Statistical Software.

Nosofsky, R. M., Palmeri, T. J., & McKinley, S. C. (1994). Rule-plus-exception model of classification learning. Psychological Review, 101, 53–79.

Orhan, A. E., & Jacobs, R. A. (2013). A probabilistic clustering theory of the organization of visual short-term memory. Psychological Review, 120, 297–328.

Parpart, P., Schulz, E., Speekenbrink, M., & Love, B. C. (2015). Active learning as a means to distinguish among prominent decision strategies. In Proceedings of the 37th Annual Meeting of the Cognitive Science Society (pp. 1829–1834).

Pashler, H. (1988). Familiarity and visual change detection. Perception & Psychophysics, 44, 369–378.

Percival, D. B., & Walden, A. T. (2006). Wavelet methods for time series analysis (Vol. 4). Cambridge University Press.

Peyton Jones, S. L. (1987). The implementation of functional programming languages (prentice-hall international series in computer science). Prentice-Hall, Inc.

Piantadosi, S. T., Tenenbaum, J. B., & Goodman, N. D. (2016). The logical primitives of thought: Empirical foundations for compositional cognitive models.

Rasmussen, C., & Williams, C. (2006). Gaussian processes for machine learning. MIT Press.

Rouder, J. N., Morey, R. D., Morey, C. C., & Cowan, N. (2011). How to measure working memory capacity in the change detection paradigm. Psychonomic Bulletin & Review, 18, 324–330.

Sanborn, A. N., Griffiths, T. L., & Shiffrin, R. M. (2010). Uncovering mental representations with Markov chain Monte Carlo. Cognitive Psychology, 60, 63–106.

Schölkopf, B., & Smola, A. J. (2002). Learning with kernels: support vector machines, regularization, optimization, and beyond. MIT press.

Schulz, E., Konstantinidis, E., & Speekenbrink, M. (2016). Putting bandits into context: How function learning supports decision making. bioRxiv, 081091.

Schulz, E., Speekenbrink, M., & Krause, A. (2017). A tutorial on gaussian process regression: Modelling, exploring, and exploiting functions. bioRxiv. Retrieved from https://www.biorxiv.org/content/early/2017/10/10/095190 doi: 10.1101/095190

Schulz, E., Tenenbaum, J. B., Reshef, D. N., Speekenbrink, M., & Gershman, S. J. (2015). Assessing the perceived predictability of functions. In Proceedings of the 37th Annual Meeting of the Cognitive Science Society (pp. 2116–2121).

Shepard, R. N., Hovland, C. I., & Jenkins, H. M. (1961). Learning and memorization of classifications. Psychological Monographs: General and Applied, 75, 1.

Sollich, P. (2001). Gaussian process regression with mismatched models. arXiv preprint cond-mat/0106475.

Stan Development Team. (2016). RStan: the R interface to Stan. Retrieved from http://mc-stan.org/ (R package version 2.14.1)

Sternberg, S., et al. (1966). High-speed scanning in human memory. Science, 153, 652–654.

Tenenbaum, J. B., Kemp, C., Griffiths, T. L., & Goodman, N. D. (2011). How to grow a mind: Statistics, structure, and abstraction. Science, 331, 1279–1285.

Thorndyke, P. W. (1977). Cognitive structures in comprehension and memory of narrative discourse. Cognitive Psychology, 9, 77–110.

Wilson, A. G., & Adams, R. P. (2013). Gaussian process kernels for pattern discovery and extrapolation. arXiv preprint arXiv:1302.4245.

Wilson, A. G., Dann, C., Lucas, C., & Xing, E. P. (2015). The human kernel. In Advances in Neural Information Processing Systems (pp. 2836–2844).

Wolpert, D. M., & Ghahramani, Z. (2000). Computational principles of movement neuroscience. Nature neuroscience, 3, 1212.

Wu, C. M., Schulz, E., Speekenbrink, M., Nelson, J. D., & Meder, B. (2017). Exploration and generalization in vast spaces. bioRxiv, 171371.

Xu, Y., Honorio, J., & Wang, X. (2017). Statistical efficiency of compositional nonparametric prediction. arXiv preprint arXiv:1704.01896.

Zhao, J., Ngo, N., McKendrick, R., & Turk-Browne, N. B. (2011). Mutual interference between statistical summary perception and statistical learning. Psychological Science.

Zhao, J., & Yu, R. Q. (2016). Statistical regularities reduce perceived numerosity. Cognition, 146, 217–222.

